# Transcriptomic rejuvenation during intestinal and liver regeneration in mice

**DOI:** 10.64898/2026.10.11.758291

**Authors:** Sharif Iqbal, Vadim N. Gladyshev

## Abstract

Aging and regeneration are fundamental biological processes, but the relationship between them is poorly understood. During intestinal regeneration, adult epithelial cells reactivate developmental programs that support repair. To investigate if this plasticity alters tissue biological age, we developed and validated intestinal tissue and epithelium-specific transcriptomic aging clocks. Across various regeneration models, epithelial states were predicted by these clocks to be younger than controls. Their expression changes were also negatively associated with intestinal stem-cell and progenitor aging signatures. The resulting younger state was characterized by YAP-associated regeneration and reduced mature enterocyte identity. In a separate irradiation model, the predicted epithelial age first increased, then fell at the peak of regeneration and later increased again during recovery. In a chemotherapy-induced injury model, epithelium in both young and old animals also reversibly transitioned through this younger state, returning to its preinjury state during recovery, although regenerative and fetal-like gene expression differed with host age. Further spatial transcriptomic analyses linked lower epithelial age with regeneration in selected injured and tumor-associated regions. In addition, liver regeneration after partial hepatectomy in old animals was also accompanied by decreased transcriptomic age, which persisted to the last sampled time point, and by pathway changes negatively associated with aging signatures. Overall, these findings suggest that mammalian regeneration is associated with a transient reduction in transcriptomic age, whose timing and persistence depend on tissue and the course of repair.

## Introduction

Regeneration restores tissue after injury and, in some animals, can also reverse features of aging. In aged sexual planarians, amputation and regeneration improve physiological function and reduce age-associated cellular and transcriptional changes in both regenerated and retained tissues (Dai *et al*, 2025). Newts can repeatedly regenerate lenses that retain youthful expression features (Eguchi *et al*, 2011; Sousounis *et al*, 2015). Mammalian repair can also alter molecular features associated with age. Regenerated mouse muscle can show lower epigenetic age (Chambers *et al*, 2025), and partial hepatectomy reverses selected age-associated chromatin changes in mouse liver (Yang *et al*, 2023). These observations raise a broader question: how do molecular features of aging change as mammalian tissues enter regeneration and then restore their differentiated functions?

Intestinal injury allows the molecular effects of regeneration to be examined in a continuously renewing epithelium. During homeostasis, *Lgr5*⁺ stem cells continuously replenish the differentiated epithelium (Barker *et al*, 2007; Beumer & Clevers, 2021). Injury recruits additional plasticity, allowing surviving stem cells and their progeny to rebuild the crypt compartment (Murata *et al*, 2020; Tetteh *et al*, 2016). This response includes developmental reversion, in which adult epithelial cells reactivate fetal-like programs after helminth infection, colitis or irradiation (Ayyaz *et al*, 2019; Nusse *et al*, 2018; Yui *et al*, 2018). Signals from the surrounding niche, including TGF-β1 and mesenchymal asporin, can induce this regenerative state (Chen *et al*, 2023; Iqbal *et al*, 2025).

YAP connects changes in the tissue environment to epithelial plasticity and the reactivation of developmental programs during repair (Cheung *et al*, 2020; Gregorieff *et al*, 2015; Moya & Halder, 2019; Yui *et al*, 2018). These states share properties with fetal epithelium without reproducing development in full (Viragova *et al*, 2024). Their emergence raises the possibility that repair also reverses some age-associated molecular changes. Epigenetic age declines during early mouse embryogenesis (Kerepesi *et al*, 2021), and experimental partial reprogramming can reduce molecular features of aging in adult tissues (Browder *et al*, 2022; Ocampo *et al*, 2016), with functional benefits demonstrated by restored vision in aged mice (Lu *et al*, 2020). Whether the endogenous plasticity of intestinal repair includes a similar reduction in age-associated features, and whether any reduction persists as epithelial cells regain their differentiated functions, remains unresolved.

The course of repair is especially important in aged tissue. Changes in Wnt signaling, Paneth-cell *Notum*, metabolism and inflammatory signaling impair intestinal regeneration (Mihaylova *et al*, 2018; Nalapareddy *et al*, 2017; Omrani *et al*, 2023; Pentinmikko *et al*, 2019), and sustained fetal-like states have been linked to defective repair in old mice (Iqbal *et al*, 2025; Korcari *et al*, 2026). Related plasticity also supports intestinal tumorigenesis (Bala *et al*, 2023; Gregorieff *et al*, 2015; Mzoughi *et al*, 2025). A younger expression state must be interpreted in relation to the cells that acquire it and their subsequent fate. Following regeneration through recovery can distinguish a temporary response to rebuilding from a lasting change in the repaired tissue.

Measuring this response requires separating changes in epithelial state from changes in the surrounding tissue. Whole-tissue transcriptomic clocks combine expression from epithelial, immune and stromal populations (Schaum *et al*, 2020; Tyshkovskiy *et al*, 2026), whose abundance and activity change after injury. Their predictions can consequently reflect both cellular aging and shifts in tissue composition (Buckley *et al*, 2023). Clocks trained on a defined compartment can reduce contributions from other populations, but still respond to differentiation and composition within that compartment. Validation in independent studies, comparison within epithelial populations and agreement with aging signatures measured separately from the clocks are needed to interpret regenerative age trajectories (Buckley *et al*, 2023; Moqri *et al*, 2024; Rutledge *et al*, 2022).

Here, we developed transcriptomic clocks for intestinal tissue and its epithelial compartment and applied them across developmental, injury and recovery datasets. We compared regenerative expression changes with separately constructed aging signatures and identified the cellular programs associated with younger predictions. Time courses, spatial analyses and tumor models then examined how these responses depend on the stage and context of repair. Finally, we analyzed liver regeneration after partial hepatectomy (Yang *et al*, 2023) to test whether the association extends to a normally quiescent organ. Together, these analyses identify reversible changes in age-associated expression during intestinal regeneration and a younger transcriptomic state maintained through the last sampled stage of old-liver regeneration.

## Results

### Intestinal transcriptomic clocks measure age-associated expression across studies

To measure age-associated expression during regeneration, we developed relative transcriptomic clocks for small-intestinal tissue, colon tissue and the epithelial compartment. Each model estimates transcriptomic age relative to a reference group from the same dataset, following the framework of Tyshkovskiy *et al* (2026). The small-intestine and colon clocks were trained on 465 and 252 profiles, respectively, and a pan-intestinal model combined these datasets after YuGene scaling (Lê Cao *et al*, 2014). With feature selection and model tuning repeated within each training fold, all three clocks predicted age in held-out datasets (**Fig. 1A-C**). Their lower accuracy when datasets were withheld than under pooled cross-validation emphasized the importance of testing generalization across experiments (**Extended Data Fig. 1A-D**). We then restricted training to sorted or purified epithelial populations, including animal-level profiles from Tabula Muris Senis (Tabula Muris Consortium, 2020). This epithelial clock used 178 animals across 14 studies and retained 2,000 genes, predicting age under nested leave-one-study-out validation with r = 0.93 and a mean absolute error of 3.0 months (**Fig. 1D**). Validation was similar when a random half of the youngest animals defined the reference, and the model correctly ordered young and old animals in four additional epithelial studies excluded from model development (**Supplementary Table 6**). These analyses establish a measure of relative epithelial transcriptomic age that can be compared with clocks trained on broader intestinal tissue profiles.

**Figure 1.**
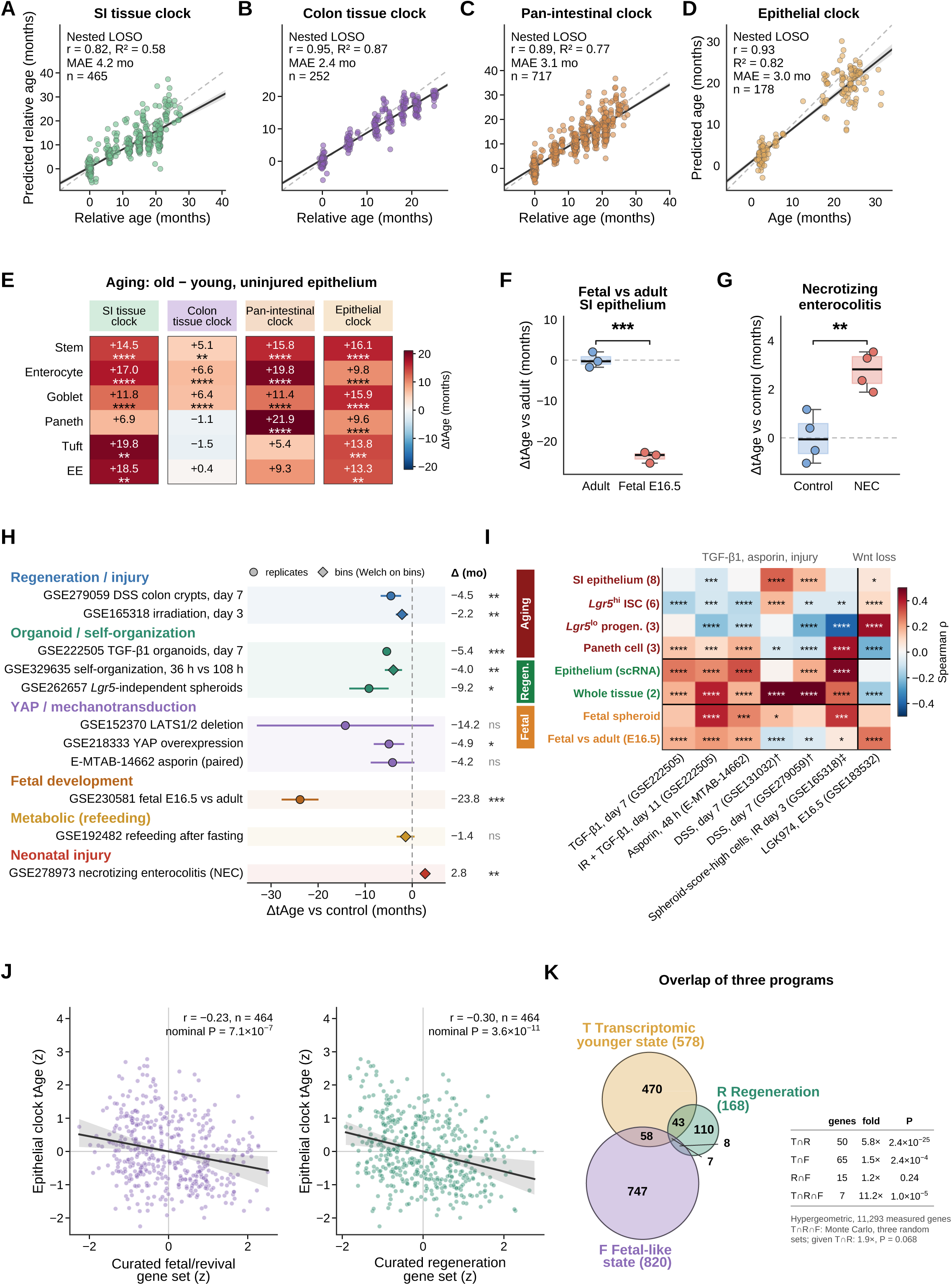
Intestinal transcriptomic clocks applied to aging, development and regeneration. **(A-D)** Nested leave-one-study-out validation of the SI tissue clock (A, 465 samples from eight studies), colon tissue clock (B, 252 samples from seven studies), pan-intestinal clock (C, 717 samples) and epithelial clock (D, 178 animals from 14 studies). Gene selection and model tuning were repeated within each training fold. A-C show ages relative to the youngest group of each series. D shows chronological ages after referencing each held-out study to its youngest group. Points represent samples, the solid line and band show a least-squares fit and 95% bootstrap confidence interval, and the dashed line indicates identity. Pearson r, RZ, mean absolute error (MAE) and n are shown. Alternative validation schemes and estimators are presented in Extended Data Fig. 1A-D. **(E)** Old-minus-young predicted age in uninjured SI epithelial populations on four clocks. Day-0 cells of each type were pooled across two mice per age and randomly divided into bins of about 30 cells, yielding 389 bins. Stars indicate Welch tests on bins with Benjamini-Hochberg (BH) adjustment across cell types within each clock. EE, enteroendocrine. **(F)** Fetal E16.5 versus adult SI epithelium (GSE230581; three freshly sorted epithelial samples per group, GSM7227026–GSM7227028 and GSM7227023–GSM7227025), referenced to adult epithelium. Welch P = 1.2 × 10 ⁴. Fetal samples are outside the adult training range, so only their relative ordering is interpreted. **(G)** Neonatal necrotizing enterocolitis versus control epithelium (GSE278973, four random bins of pooled cells from two mice per group). Stars indicate a Welch test on bins. In F and G, boxes show medians and interquartile ranges, whiskers extend to 1.5 times the interquartile range, and points represent samples or bins. **(H)** Epithelial-clock differences between regenerative states and their study-specific controls. Circles represent comparisons based on replicate animals, cultures or donor-line means. Diamonds represent comparisons based on random bins of cells from one library per condition (one sample, or the pooled cells of several mice for irradiation and necrotizing enterocolitis); their intervals and tests are nominal and computed across bins of that library. Bars show 95% confidence intervals calculated over the corresponding units. The irradiation bins contain pooled cells from the mice sampled on each day and are referenced by cell type to day 0. LATS1/2 profiles were averaged within mouse, with two mice per group. Spheroid and organoid profiles were averaged within donor, with three and two donors, respectively. Asporin was tested across four donor-matched pairs. Other comparisons used Welch tests, with BH adjustment across all 11 displayed contrasts. The NEC comparison has no interval because only its bin-level P value was available. Time courses show the time point with the highest regeneration score. All time points are listed in Supplementary Table 2. Individual observations appear in Extended Data Fig. 2 and Fig. 2. Tests on bins in E, G and H describe the sampled pooled cells and do not provide animal-level replication. **(I)** Spearman correlations between regenerative-contrast log₂ fold changes and intestinal aging, regeneration and fetal reference signatures, calculated over the union of the 1,000 most significant genes in each pair. Six regenerative contrasts and one fetal Wnt-secretion-inhibition contrast are shown. Color indicates ρ. Stars denote BH-adjusted correlations across 56 pairs, treating genes as independent. These significance estimates do not account for gene dependence. † indicates DSS contrasts used to construct the whole-tissue regeneration reference. ‡ indicates contrasts with one pooled profile per group and genes ranked by absolute log₂ fold change. Reference construction and shared source datasets are detailed in Methods. **(J)** Epithelial age versus scores of the 22-gene composite fetal/revival marker panel and the 26-gene curated repair/YAP-target set (both listed in Supplementary Table 5; the composite panel differs from the curated revival set of K), across 464 profiles standardized within dataset and cell type. Lines show least-squares fits and bands show 95% confidence intervals from 2,000 resamples of dataset-cell-type clusters. Pearson r and nominal P are shown. Dataset-level mixed models are reported in Supplementary Table 5. **(K)** Overlap of younger-state upregulated genes (T, 578), the measured regeneration reference (R, 168) and the measured top 1,000 fetal-higher genes of the GSE115541 fetal-versus-adult intestine reference (F, 820; Methods), within 11,293 measured genes. Pairwise enrichment uses hypergeometric tests. The three-way overlap of seven genes is compared with three independent random sets by Monte Carlo simulation. Conditional on the 50 genes shared by T and R, fetal overlap is 1.9-fold enriched, P = 0.068. tAge, transcriptomic age. ΔtAge, difference from the indicated reference. SI, small intestine. All clocks use BayesianRidge unless stated otherwise. Stars indicate P or q < 0.05 (*), < 0.01 (**), < 0.001 (***) or < 10 ⁴ (****). ns, not significant.

The clocks also distinguished young and old epithelium in the uninjured controls of our 5-FU experiment and in a published single-cell aging study (GSE210669) (Choi *et al*, 2023), with differences among clocks in the size of the age separation (**Fig. 1E, Extended Data Fig. 1E**). In a *Citrobacter rodentium* infection study, tissue-clock predictions were higher in old than in young colon, although the infection response depended on the clock and the age-by-infection interaction was not significant (GSE288285, **Extended Data Fig. 1F**) (Korcari *et al*, 2026). Related data from the same publication contributed to training the epithelial clock, so this analysis does not test its performance in an independent study. Across additional cohorts, including a study of GLP-1 receptor agonism (Huang *et al*, 2025), predicted age consistently increased with chronological age, whereas responses to metformin and other interventions were small or inconsistent across models (**Extended Data Fig. 1G,H, S**upplementary Table 3). Tissue-clock predictions also increased with age in non-intestinal tissues, indicating that they retain some shared mammalian aging signals (**Supplementary Table 6**). Thus, the models reliably ordered age, while intervention sensitivity depended on the model and experimental setting.

### Regenerative epithelial states are predicted to be younger and partly reverse age-associated expression

Fetal epithelium provides a developmental reference for the programs reactivated during adult repair and gives rise to adult intestinal stem cells (Guiu *et al*, 2019). In small-intestinal expression profiles, fetal epithelium was predicted to be younger than adult epithelium (GSE230581, **Fig. 1F)** (Pikkupeura *et al*, 2023). Injury did not invariably produce a younger prediction. In neonatal necrotizing enterocolitis, which combines epithelial injury with impaired regeneration (Xiong *et al*, 2026), affected epithelium was predicted to be older than neonatal controls (GSE278973, **Fig. 1G)**. These comparisons establish the direction of the clock response in developmental and disease contexts, although fetal and neonatal tissues lie outside its adult training range.

We next examined adult regenerative states, comparing each with its control from the same study. These encompassed DSS colitis and irradiation, organoid culture and self-organization, YAP activation or altered mechanotransduction, and refeeding after fasting (Chen *et al*, 2023; Cheung *et al*, 2020; Imada *et al*, 2024; Iqbal *et al*, 2025; Lu *et al*, 2023; Marefati *et al*, 2025; Pikkupeura *et al*, 2023; Schwayer *et al*, 2026; Xiao *et al*, 2025). For time courses, the comparison used the sampled time point with the highest regeneration score, selected without reference to predicted age. At these time points, all nine adult regenerative states were predicted to be younger than their controls (**Fig. 1H**). Four of the six comparisons with replicate animals or cultures were significant after adjustment for multiple testing. The remaining comparisons used pooled-cell bins, with statistical estimates based on variation among bins within each condition. Asporin-treated organoids were younger in all donor-matched pairs, although the paired comparison was not significant. YAP activation and TGF-β1 exposure produced significant decreases, while the LATS1/2-deficient cohort showed the same direction with greater uncertainty (**Extended Data Fig. 2A-D**). Other time points were not uniformly younger, indicating that the response depends on the stage of regeneration (Supplementary Table 2).

Organoid comparisons helped separate regenerative state from differences between experimental systems. Fetal epithelium forms poorly differentiated spheroids in culture (Mustata *et al*, 2013), and an *Lgr5*-independent adult lineage can also form spheroids and contribute to repair (Marefati *et al*, 2025). These spheroids and progenitor-like states during organoid self-organization were predicted to be younger than their more differentiated counterparts (**Extended Data Fig. 2E,F)** (Schwayer *et al*, 2026). Because stem and differentiated epithelial cells also differ in predicted age before injury, we examined the contribution of enterocyte identity. Adjustment for an enterocyte-expression score retained the younger response in DSS colitis, re-plated organoids and irradiated cell types, although identity and age predictions could not be separated during organoid self-organization. In a descriptive linear approximation, in which each clock gene’s log2 fold change was multiplied by its coefficient scaled to the training standard deviation of its clock input, genes differing between enterocytes and stem cells accounted for a median of 61% of the summed approximation (20% to 94% across 13 contrasts), and the remaining clock genes summed to a decrease in every contrast (**Supplementary Table 4**). These results connect the younger prediction to epithelial plasticity while showing that reduced enterocyte identity is not its only source. During organoid self-organization and DSS colitis, an initial increase in predicted age preceded the regenerative decrease, followed by a later return toward or above the control level (**Extended Data Fig. 2G,H**). The younger state represents a phase of these responses, not the complete course of injury and repair.

We then compared regeneration with aging expression changes without using the clock predictions. We assembled aging meta-signatures for *Lgr5*-high stem cells, progenitors, Paneth cells, small-intestinal tissue and purified epithelium from multiple studies (Mihaylova *et al*, 2018; Nefzger *et al*, 2022; Pentinmikko *et al*, 2019; Tyshkovskiy *et al*, 2026). Regeneration references combined published injury datasets, and developmental references described fetal spheroids and fetal versus adult epithelium (Czarnewski *et al*, 2019; Kolde *et al*, 2012; Mustata *et al*, 2013; Pikkupeura *et al*, 2023; Xiao *et al*, 2025). The corresponding scores measured regenerative and fetal-like expression separately from the clocks. Comparisons within annotated cell types showed that predicted age and these scores did not always change together (**Extended Data Fig. 3A-D**). During organoid self-organization, stem and progenitor populations became younger without uniformly increasing the regenerative or fetal-like scores. In irradiation-derived organoids, TGF-β1 reduced predicted age even when irradiation alone increased it (Chen *et al*, 2023). These differences prompted us to examine which components of aging and development were shared with each regenerative state.

At the gene level, five of six regenerative comparisons opposed the stem-cell aging signature and four opposed the progenitor signature, while most correlated positively with the Paneth-cell aging signature (**Fig. 1I**). These comparisons included TGF-β1 and asporin exposure, DSS injury and regenerating epithelium after irradiation (Chen *et al*, 2023; Czarnewski *et al*, 2019; Iqbal *et al*, 2025; Lu *et al*, 2023; Xiao *et al*, 2025). By contrast, inhibiting Wnt secretion in fetal intestine shifted expression toward the stem-cell and progenitor aging signatures (Maimets *et al*, 2022), consistent with the contribution of impaired Wnt signaling to aged stem-cell dysfunction (Nalapareddy *et al*, 2017). All six regenerative comparisons correlated positively with the regeneration meta-signatures. Regeneration opposed selected stem-cell and progenitor aging changes, with the relationship depending on the cellular aging reference.

The same comparisons distinguished fetal progenitor programs from the broader fetal transcriptome. TGF-β1-treated organoids, asporin-treated organoids and irradiated epithelium resembled the fetal spheroid program more closely than the fetal-versus-adult tissue comparison, while both DSS responses opposed the latter (**Fig. 1I**) (Chen *et al*, 2023; Czarnewski *et al*, 2019; Iqbal *et al*, 2025; Lu *et al*, 2023; Mustata *et al*, 2013; Pikkupeura *et al*, 2023; Xiao *et al*, 2025). Several injury and niche-signal responses also increased inflammatory and matrix-associated programs and reduced oxidative and fatty-acid metabolism, while their cell-cycle responses differed (**Extended Data Fig. 4A,B**). Thus, the regenerative states shared features of fetal progenitors without reproducing fetal epithelial development as a whole.

### Younger epithelial states share regenerative programs and reduced enterocyte identity

To identify expression changes recurring across younger states, we combined the nine adult regenerative comparisons with the fetal-versus-adult comparison. For this descriptive signature, time courses contributed the state with the lowest epithelial age. The resulting signature contained 578 upregulated and 550 downregulated genes that changed consistently across the measured comparisons (**Extended Data Fig. 5A,B**). Upregulated genes included the revival stem-cell markers *Clu* and *Anxa1,* the chaperones *Hspa1a* and *Hspa1b*, *Msln* and several S100 genes (Ayyaz *et al*, 2019). They also included the Hippo-YAP regulators *Ajuba* and *Wwc2* and the YAP transcriptional partner *Tead4* (Das Thakur *et al*, 2010; Wennmann *et al*, 2014; Zhao *et al*, 2008). Downregulated genes included nutrient transporters and metabolic enzymes characteristic of mature enterocytes, including *Slc5a1*, *Slc13a1*, *Fbp1*, *Otc*, *Ces1d* and *Ces1f* (Haber *et al*, 2017; Moor *et al*, 2018). The recurrent changes combined regenerative and stress responses with reduced mature epithelial differentiation.

Regulatory analyses supported this change in epithelial state. Activities inferred from target-gene expression were generally lower for *Cdx2* and *Hnf1a* and higher for AP-1, NF-κB, STAT3, SMAD and HSF1 (**Extended Data Fig. 6A**) (Badia-i-Mompel *et al*, 2022; Müller-Dott *et al*, 2023). PROGENy analysis similarly inferred increased hypoxia, p53, TGF-β and NF-κB signaling in most comparisons (**Extended Data Fig. 6B**) (Schubert *et al*, 2018). These expression patterns associate the younger state with altered differentiation and repair-associated signaling, including pathways also activated by cellular stress.

We next asked whether proliferation alone could account for the association between regeneration and predicted age. Across the epithelial profiles, higher fetal/revival and regeneration scores accompanied lower epithelial age (**Fig. 1J**). The regeneration association remained after adjustment for a proliferation score and in models accounting for differences among datasets (**Extended Data Fig. 5C, Supplementary Table 5**). Thus, cell-cycle expression did not fully explain the younger predictions, although proliferation and epithelial reprogramming remain closely related during repair (Iqbal *et al*, 2025; Metcalfe *et al*, 2014; Yui *et al*, 2018).

The younger-state signature was strongly enriched for regeneration genes **and** included all 22 measured genes of the curated 25-gene revival stem-cell set and all 15 measured genes of the curated YAP-target set (**Fig. 1K, Extended Data Fig. 5D**) (Ayyaz *et al*, 2019; Cheung *et al*, 2020; Gregorieff *et al*, 2015; Kolde *et al*, 2012; Yui *et al*, 2018). Enrichment for fetal genes was much weaker and depended on the size of the fetal reference set. This difference, together with the limited overlap between the regeneration and fetal references, places the younger-state signature closer to a repair program than to a complete return to the fetal transcriptome (Mustata *et al*, 2013; Yui *et al*, 2018).

YAP perturbations further connected this signature to regenerative regulation. The YAP-overexpression and *LATS1/2*-deletion datasets used to construct the signature increased its score, as expected (**Extended Data Fig. 5E**). In a separate dataset of *Apc*-mutant organoids not used to construct the signature, inducible YAP activation increased the score over time, while constitutively active YAP in *Apc/Trp53*-mutant organoids produced the same direction without statistical significance (**Extended Data Fig. 5F**) (Cheung *et al*, 2020; Gregorieff *et al*, 2015; Pikkupeura *et al*, 2023). Repairing Sca1-positive colonic epithelium also expressed the signature more strongly than homeostatic epithelium in a separate microarray dataset (**Extended Data Fig. 5G**) (Yui *et al*, 2018).

Chromatin accessibility provided a complementary view of epithelial regulation. Intestinal progenitors can reorganize lineage-associated chromatin as they regain stem-cell function after injury (Jadhav *et al*, 2017). In organoids derived after irradiation and exposed to TGF-β1, chromVAR deviations were higher at TEAD, AP-1, SMAD and NF-κB motifs and lower at HNF4/CDX2 motifs than in non-irradiated vehicle controls (GSE222505, **Extended Data Fig. 5H**) (Chen *et al*, 2023). STAT3 motif accessibility did not change, despite higher inferred activity in the expression analysis. Increased TEAD accessibility is consistent with YAP-associated transcription, while lower HNF4/CDX2 accessibility fits the loss of mature enterocyte features (Verzi *et al*, 2013; Zhao *et al*, 2008). Because this comparison combines irradiation history with TGF-β1 exposure, it does not isolate either effect. Together, the expression and accessibility analyses associate the younger epithelial state with reorganization of repair and differentiation programs.

### Epithelial transcriptomic age changes during injury and recovery

To resolve the timing of the younger epithelial state, we analyzed a single-cell irradiation time course (GSE165318, **Fig. 2**) (Lu *et al*, 2023). Epithelial expression diverged from baseline after injury and moved back toward it during recovery, both in pooled epithelium and within stem cells and enterocytes (**Fig. 2A**). The epithelial clock predicted an older pooled epithelium during the early injury response and a younger state at day 3, when regeneration scores peaked. Stem cells did not show the early age increase observed in enterocytes, and their regeneration score was already elevated at day 1. During recovery, predictions in stem cells and enterocytes approached baseline, while pooled epithelial predictions increased above it (**Fig. 2B**). Fetal-like scores changed less than regeneration scores and were below baseline by day 14. The age trajectory was thus associated with regenerative activity but also depended on the epithelial population being measured.

**Figure 2.**
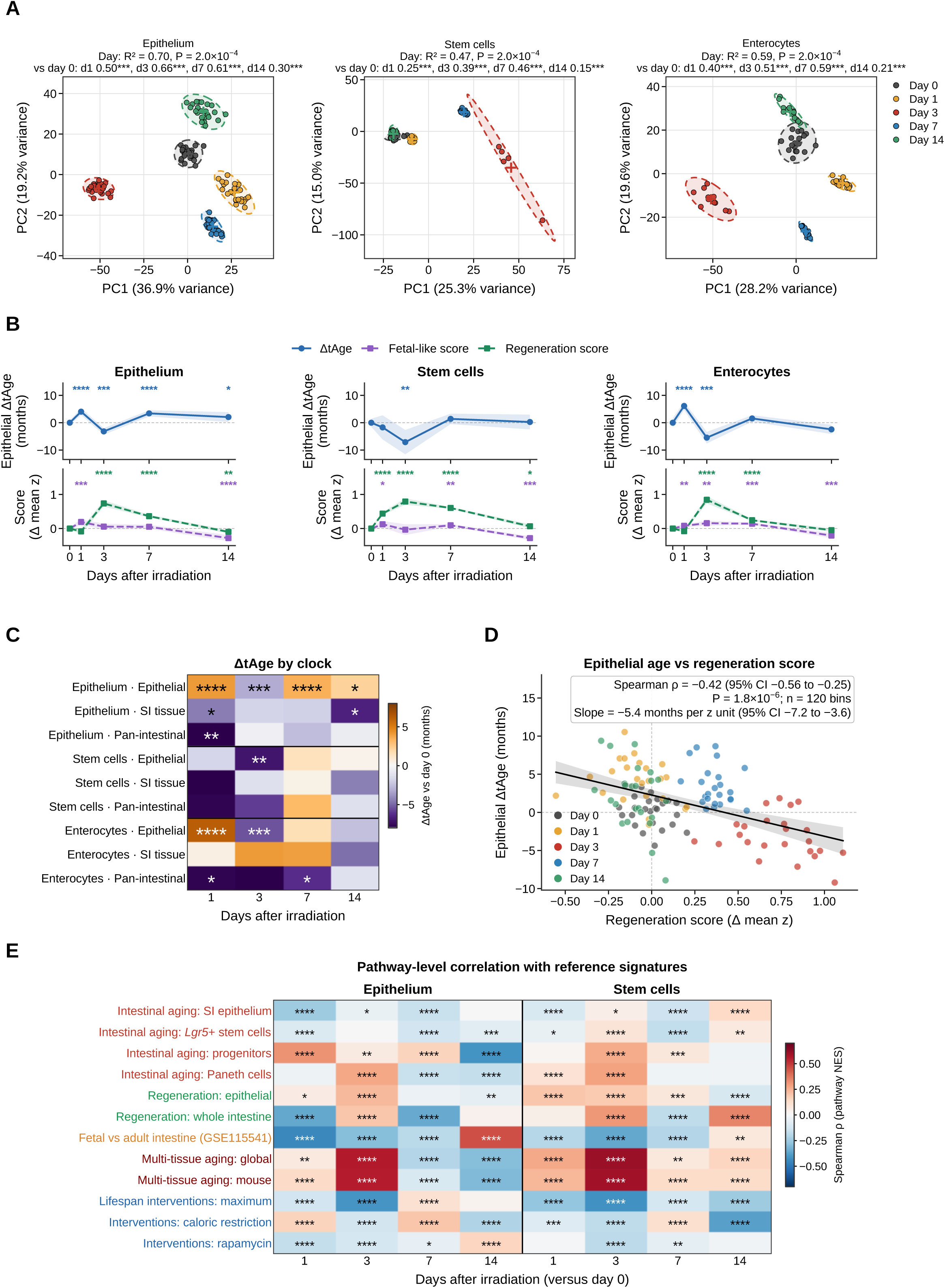
Epithelial transcriptomic age during regeneration after irradiation. Small-intestinal single-cell profiles from male C57BL/6J mice before and 1, 3, 7 and 14 days after 15 Gy abdominal irradiation (GSE165318). The series contains 17 mice, with four, four, three, three and three mice at the respective days. Cells were pooled within each day before forming 24 random bins of pooled epithelium and bins of about 30 stem cells or enterocytes, with 5-18 bins per population and day. All tests in A-D therefore describe differences among bins of pooled cells, not differences among animals. **(A)** Principal component analysis of pooled epithelium, stem cells and enterocytes using 2,000 variable genes from standardized log₂ counts per million. Crosses indicate group centroids and ellipses show 95% normal-theory regions. Labels give the variance explained by day (PERMANOVA RZ), with an overall permutation P from 4,999 permutations and BH-adjusted pairwise comparisons against day 0. **(B)** Epithelial ΔtAge and fetal-like and regeneration scores relative to day 0. Lines and bands show means and 95% confidence intervals over bins, with a common y range for each row. Stars show Mann-Whitney comparisons with day 0, BH-adjusted within panel and colored by measure. The fetal-like and regeneration scores use the 300 leading genes of the E16.5-versus-adult and epithelial regeneration references, respectively. **(C)** Mean ΔtAge on the epithelial, SI tissue and pan-intestinal clocks for each population and day. Color indicates the mean difference from day 0. Stars denote bin-level day-versus-baseline tests as in B. **(D)** Epithelial ΔtAge versus regeneration score across 120 pooled-epithelium bins, colored by day. The line and band show a least-squares fit and its 95% confidence interval. Spearman ρ, its Fisher-z confidence interval and nominal P, and the fitted slope with its confidence interval are shown. **(E)** Spearman correlations of pathway normalized enrichment scores (NES) between each day-versus-day-0 response and intestinal aging, regeneration, fetal, multi-tissue aging and lifespan-intervention references. Responses were ranked by log₂ fold change in pooled day pseudobulks and analyzed by pre-ranked GSEA, with 20,000 random gene sets and 1,519-2,904 pathways shared by each pair. Color indicates ρ. Stars show BH-adjusted correlation P values across all 468 contrast-reference pairs of the two time courses (26 contrasts, irradiation and 5-FU, epithelium and stem cells, × 18 references, including references not displayed). These calculations treat overlapping pathways as independent and are descriptive. Extended Data Fig. 10A shows the full matrices for days 3, 7 and 14. *Star* thresholds are as in Fig. 1.

Tissue-clock responses differed from the epithelial-clock trajectory and did not consistently identify the day-3 minimum (**Fig. 2C**). Across pooled epithelial bins from the five sampled days, higher regeneration scores accompanied younger epithelial predictions, largely reflecting differences between time points (**Fig. 2D)**. Pathway changes followed a different course. At peak regeneration, pooled epithelium and stem cells were positively associated with multi-tissue aging signatures, while pooled epithelium during later recovery opposed progenitor and multi-tissue aging signatures and resembled the fetal reference (**Fig. 2E**). The clock minimum and the reversal of broader aging-associated pathways were not synchronous. These comparisons distinguish a younger regenerative expression state from a uniform reversal of the molecular responses associated with aging.

We examined candidate niche signals associated with this state using ligand-receptor inference and target-gene analysis (Dimitrov *et al*, 2024; Dimitrov *et al*, 2022; Türei *et al*, 2021). Revival-marker-high cells expressed inflammatory and matrix-associated genes, including the TGF-β-responsive genes *Ctgf, Serpine1* and *Jun* (**Extended Data Fig. 4C,F**) (Dennler *et al*, 1998; Igarashi *et al*, 1993; Pertovaara *et al*, 1989). Increased *Ctgf* also fits YAP-TEAD-dependent transcription (Zhao *et al*, 2008). Repeating the interaction analysis in *Ly6a*-expressing epithelial cells identified candidate *Cd44*-associated interactions in both the irradiation and 5-FU datasets, with different predicted partners in the sampled cell populations (**Extended Data Fig. 4D,E, Supplementary Table 5**). These findings connect the regenerative state to candidate inflammatory and matrix signals without establishing direct signaling activity. Experimental studies showing that TGF-β1 and mesenchymal asporin induce epithelial reprogramming provide a biological context for these associations (Chen *et al*, 2023; Iqbal *et al*, 2025).

We next compared repair in young and old mice after 5-FU injury, which targets proliferating cells (Iqbal *et al*, 2025; Longley *et al*, 2003; Pentinmikko *et al*, 2019). Small-intestinal epithelial cells were profiled from young and old females before treatment and through six days of recovery, with two mice per age and collection day before quality-control exclusions. Day-2 samples and young day-3 samples yielded too few cells for analysis. *Ly6a* expression increased in both age groups, with a larger and more sustained induction in old epithelium, while *Clu* induction was stronger in young epithelium. Both metacell measurements and per-mouse pseudobulks supported these differences (**Extended Data Fig. 7A-D, Supplementary Table 4**). The distinct marker responses are consistent with previous evidence that fetal-like epithelial states persist longer in aged intestine after injury (Iqbal *et al*, 2025; Korcari *et al*, 2026).

Time after injury accounted for most of the separation among epithelial expression profiles (**Fig. 3A**). At the mouse level, epithelial age was lower at day 4 in both young and old animals and no longer differed from baseline by day 6 (**Fig. 3B, Extended Data Fig. 8C**). Regeneration scores increased in both age groups, while fetal-like expression appeared earlier in young epithelium. Higher regeneration scores accompanied younger age predictions across mice and within the metacell profiles (**Fig. 3C, Supplementary Table 5**). To test how much the clock response depended on known developmental and repair programs, we retrained the model after excluding fetal-versus-adult differentially expressed genes and the regeneration, repair, revival and younger-state gene sets. This reduced age-prediction accuracy but retained a younger response after 5-FU in both metacell and mouse-level analyses (**Extended Data Fig. 9A-D, Supplementary Table 6**). The response was not confined to genes used to define these programs. Capping clock inputs at their observed training ranges retained the day-4 decrease while attenuating the metacell differences (**Extended Data Fig. 9E**). These sensitivity analyses support a reproducible direction of change within the principal modeling framework.

**Figure 3.**
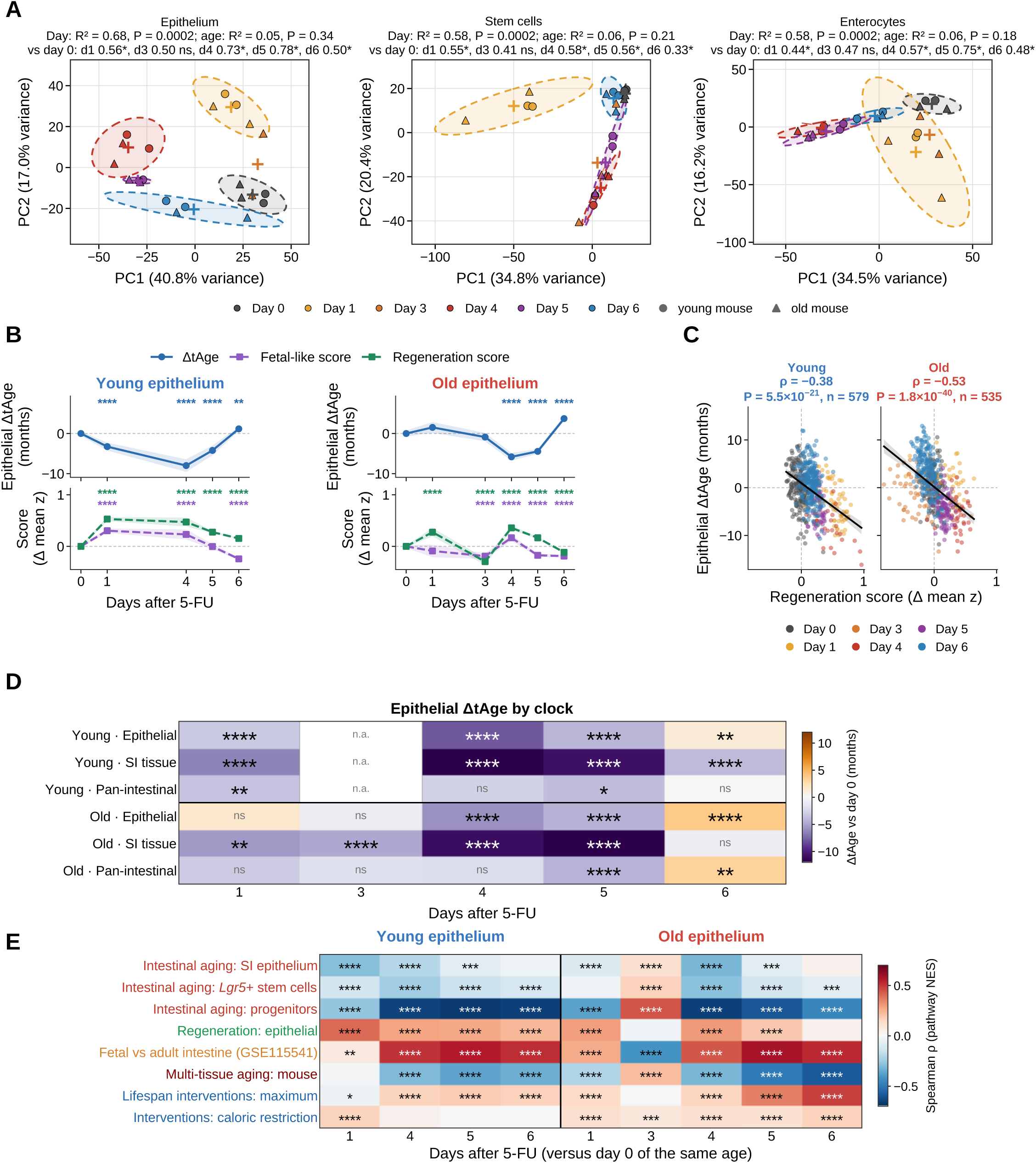
Epithelial transcriptomic age after 5-FU injury in young and old mice. Single-cell RNA sequencing of small-intestinal epithelium after 5-FU treatment, with two young (4-month-old) and two old (21-month-old) mice collected at each sampled day. Day 2 in both ages and day 3 in young mice yielded insufficient cells and were not analyzed. A total of 44,815 cells passed the cell filters, 44,792 of them in the 22 samples analyzed; the low-yield old day-1 sample is included in all panels. **(A)** Principal component analysis of per-mouse pseudobulk profiles of pooled epithelium, stem cells and enterocytes. Circles represent young mice and triangles old mice. Crosses indicate group centroids and ellipses show 95% normal-theory regions. PERMANOVA assesses day and age as described in Methods. With four mice at a day, the smallest attainable pairwise permutation P is approximately 0.03 when age groups are pooled. **(B)** Epithelial ΔtAge on the YuGene-BayesianRidge clock, and fetal-like and regeneration scores, relative to day 0 of the same age. Lines and bands show means and 95% confidence intervals over metacells, with a common y range within each row. Stars denote Mann-Whitney comparisons with day 0, BH-adjusted within age and measure and colored by measure. **(C)** Epithelial ΔtAge versus regeneration score across young and old metacells. Lines and bands show least-squares fits and 95% confidence intervals. Spearman ρ, nominal P and metacell counts are given. Day colors match A. **(D)** Mean ΔtAge on the epithelial, SI tissue and pan-intestinal clocks for pooled epithelium. Colors show the mean difference from day 0 and stars follow B. n.a., not analyzed. Panels B-D use metacells from the sampled mice. Their tests describe differences among metacells and do not provide animal-level inference. Per-mouse estimates are shown in Extended Data Fig. 8C and modeled in Supplementary Table 4. **(E)** Pathway-level correlations of each day-versus-baseline epithelial response with intestinal aging, epithelial regeneration, fetal, multi-tissue aging and lifespan-intervention references. Responses were calculated from per-mouse pseudobulk profiles and ranked by moderated t statistics. Fig. 2E instead uses log₂ fold changes from pooled mice, so correlation magnitudes are not directly interchangeable between the two figures. Correlation and star definitions follow Fig. 2E. The full matrix appears in Extended Data Fig. 10B. *Star* thresholds are as in Fig. 1.

The response nevertheless depended on how transcriptomic age was measured. The tissue clocks also produced younger predictions during the 5-FU time course, but their timing differed from that of the epithelial clock (**Fig. 3D, Extended Data Fig. 8A,B**). Changing the final estimator to ElasticNet did not reproduce the primary model’s day-4 decrease, and quantile-normalized input altered the trajectory (**Extended Data Fig. 7E-H, Supplementary Table 4**). Associations with regenerative expression also varied by population, being present in stem cells of both age groups but in enterocytes only in young mice (**Extended Data Fig. 8D**). The day-4 decrease persisted in several sensitivity analyses but was not reproduced by every estimator.

Pathway analysis placed these age responses within the progression from injury to repair (**Fig. 3E, Extended Data Fig. 10A-D**). From day 4, expression changes in young and old epithelium generally opposed progenitor and multi-tissue aging signatures and are positively associated with fetal and lifespan-intervention signatures. Old epithelium at day 3 retained the earlier pattern, with associations in the opposite direction. By day 6, the pathway response in old epithelium still opposed several aging signatures, even though the younger mouse-level clock estimate was no longer evident. The two measurements captured overlapping but temporally distinct features of regeneration: a transient reduction in epithelial transcriptomic age and a broader reorganization of age-associated pathways.

### Spatial transcriptomics identifies a younger epithelial state during intestinal repair

The irradiation and 5-FU time courses linked regeneration to a younger epithelial transcriptomic state. To examine its spatial distribution, we analyzed small-intestinal sections before and three days after irradiation (GSE227742) (Das *et al*, 2025). Regeneration and fetal-like scores increased after injury, and epithelial spots were predicted to be younger than those in the uninjured section (**Fig. 4A,B**). After neighboring epithelial spots were pooled into metaspots, the expression scores remained elevated and mean predicted age remained lower, although the age comparison was not significant (**Extended Data Fig. 11A,B**). Lower spot-level predictions occurred across all eight dominant epithelial labels, including stem, secretory and absorptive populations (**Fig. 4C**).

**Figure 4.**
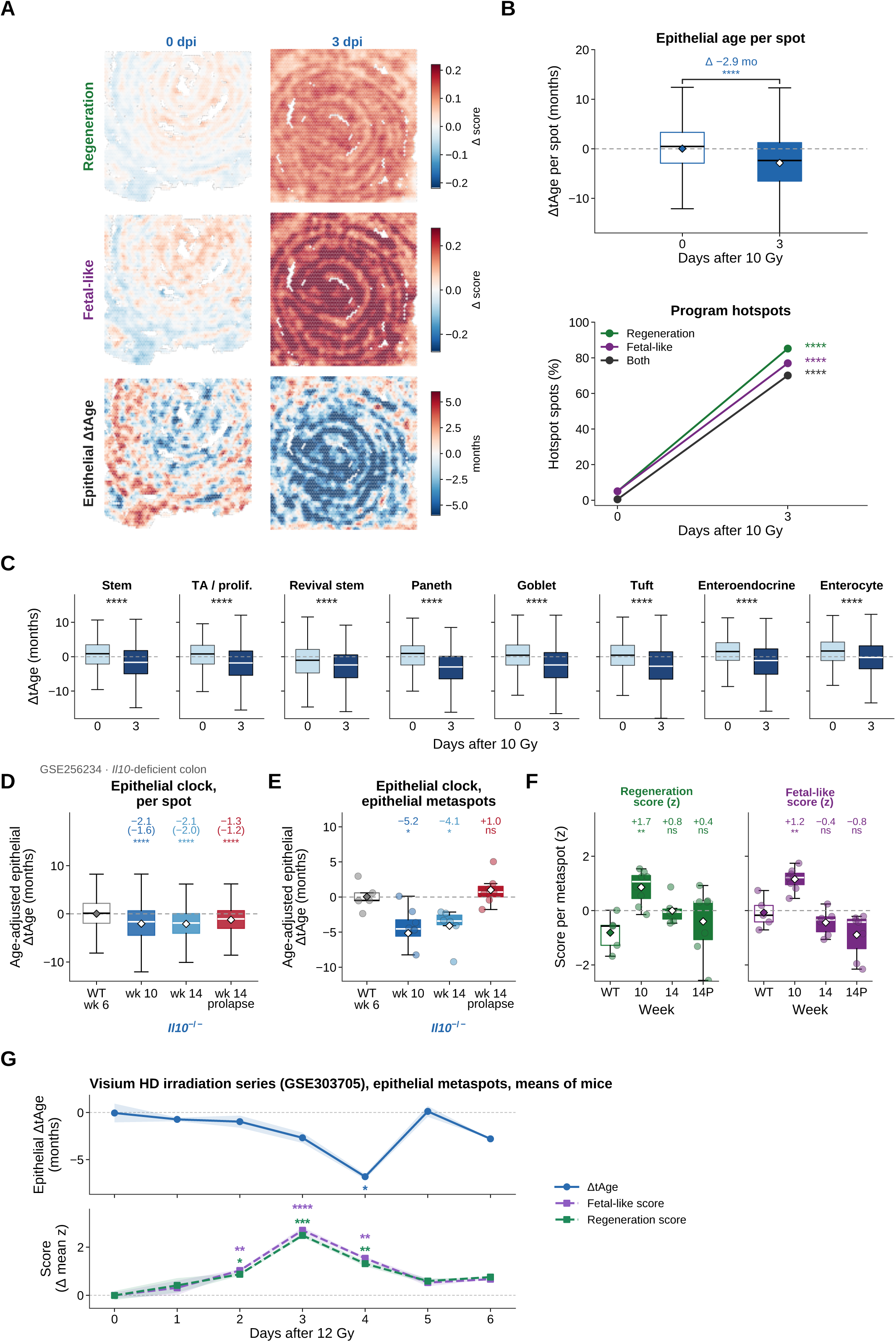
Spatial epithelial age after irradiation and in chronic colitis. **(A)** Per-spot maps of irradiated SI (GSE227742) before and 3 days after 10 Gy irradiation (dpi) on standard diet, showing regeneration and fetal-like scores minus the mean of the 0-dpi section, and epithelial ΔtAge relative to that section (seven-nearest-neighbor smoothing for display only). **(B)** Quantification of A. Top, per-spot epithelial ΔtAge (boxes, medians and interquartile ranges; diamonds, means; Δ, reference-relative mean prediction; Mann–Whitney P). Bottom, proportions of spots above the 95th percentile of the 0-dpi section (Both, spots above the thresholds for both the regeneration and fetal-like scores); stars compare 3 dpi with 0 dpi (Fisher’s exact test, BH; all q < 10 ⁵). **(C)** Epithelial ΔtAge per spot, grouped by the cell type with the highest marker-gene score, at 3 dpi versus 0 dpi (pale blue, 0 dpi; dark blue, 3 dpi; Mann–Whitney, BH). **(D–F)** *Il10*-deficient colon (GSE256234) at weeks 10 and 14 and at week 14 with rectal prolapse (14P), relative to the wild-type week-6 section. In D and E, the chronological age elapsed since week 6 (week 10, 4 weeks = 0.93 months; week 14, 8 weeks = 1.87 months) is subtracted from each estimate (age-adjusted ΔtAge). **(D)** Age-adjusted epithelial ΔtAge of every spot (numbers, mean difference with the depth-adjusted estimate in parentheses). **(E, F)** Age-adjusted epithelial ΔtAge (E) and regeneration and fetal-like scores (**F**; z-scored across metaspots) of epithelial metaspots (spots of the crypt-base, transit-amplifying, differentiated and revival compartments pooled by k-means on their coordinates; at least 200,000 counts; five to eight metaspots per section, because spots of these colon sections carry a median of 270 to 880 counts). Boxes, median and interquartile range; whiskers, 1.5 × interquartile range; dots, metaspots; diamonds, means; numbers, mean difference from the wild-type section; stars, Mann–Whitney test across spots (D) or metaspots (E, F). In A–F, spots and metaspots are spatially dependent subsamples of one section per condition, so the P values are nominal and compare spots or metaspots within single sections and do not test differences between sections or animals. **(G)** Means across mice of epithelial ΔtAge (top) and of the regeneration and fetal-like scores (bottom; z-scored across metaspots, minus day 0) of epithelial metaspots (320 µm, at least 200,000 counts) in Visium HD jejunum before (day 0) and 1 to 6 days after 12 Gy whole-body irradiation (GSE303705; 23 mice in the deposited experiment, of which 20 could be assigned from the authors’ bin-level labels and analyzed, and 17 contributed metaspots passing the count threshold; four capture areas in the source experiment, each containing a day-0 mouse; retained metaspot and mouse numbers in Extended Data Fig. 13); shading, s.e.m. across mice. The numbers of mice contributing to the metaspot analysis at days 0 to 6 were 3, 2, 3, 3, 2, 3 and 1, respectively. Each mouse is summarized by the mean of its metaspots, and each day is compared with day 0 by ordinary least squares with capture area as a covariate (y ∼ day + capture area; Benjamini–Hochberg adjustment across days within each readout); the day-6 mouse is not tested.

We then examined chronic colitis, in which repeated injury and repair occur during persistent inflammation. Sections from *Il10*-deficient mice at 10 and 14 weeks, including a 14-week mouse with prolapse, were compared with a wild-type section at six weeks (GSE256234) (Fransson *et al*, 2026). We adjusted for the elapsed chronological time from the colitis predictions to align them with the reference age. All three colitis sections had lower mean spot-level predictions (**Fig. 4D**). Metaspot predictions were also lower at weeks 10 and 14, but not in the prolapse section (**Fig. 4E**). Mean regeneration and fetal-like scores were elevated only at week 10 (**Fig. 4F**), although spots with high regeneration scores occupied a larger fraction of the later sections (**Extended Data Fig. 12A,B**). The relationship between regenerative expression and predicted age varied across these colitis samples and with spatial aggregation.

We next followed acute repair in a Visium HD jejunal time course (GSE303705) (Ofer *et al*, 2026). After filtering, 17 mice contributed epithelial metaspots to the age analysis. Post-injury time points were compared with baseline using mouse means and adjusting for capture area. Regeneration and fetal-like scores peaked at day 3 and remained elevated at day 4 (**Fig. 4G, Extended Data Fig. 13A**). Epithelial predictions were youngest at day 4 and returned close to baseline by day 5, while the smaller day-3 decrease was not significant (**Fig. 4G, Extended Data Fig. 13B**). The small-intestinal tissue clock showed the same direction on days 2 to 4 without significant differences from baseline (**Extended Data Fig. 13C**). Both day-4 mice were profiled with an uninjured control on the same capture area. These spatial analyses support a transient younger epithelial state during acute repair, with a less uniform response during chronic inflammation.

### A younger epithelial state accompanies regenerative activity during tumor development

Regenerative and fetal-like programs also support intestinal tumor development (Bala *et al*, 2023; Gregorieff *et al*, 2015; Mzoughi *et al*, 2025). We then asked whether the epithelial age response during repair extends to colitis-associated tumorigenesis. Spatial analysis of colon after azoxymethane and DSS treatment showed progressive expansion of regeneration and fetal-like programs (AOM/DSS, GSE227598, **Fig. 5A,B**) (Das *et al*, 2025). By day 43, most spots had high regeneration scores and more than half had high fetal-like scores.

**Figure 5.**
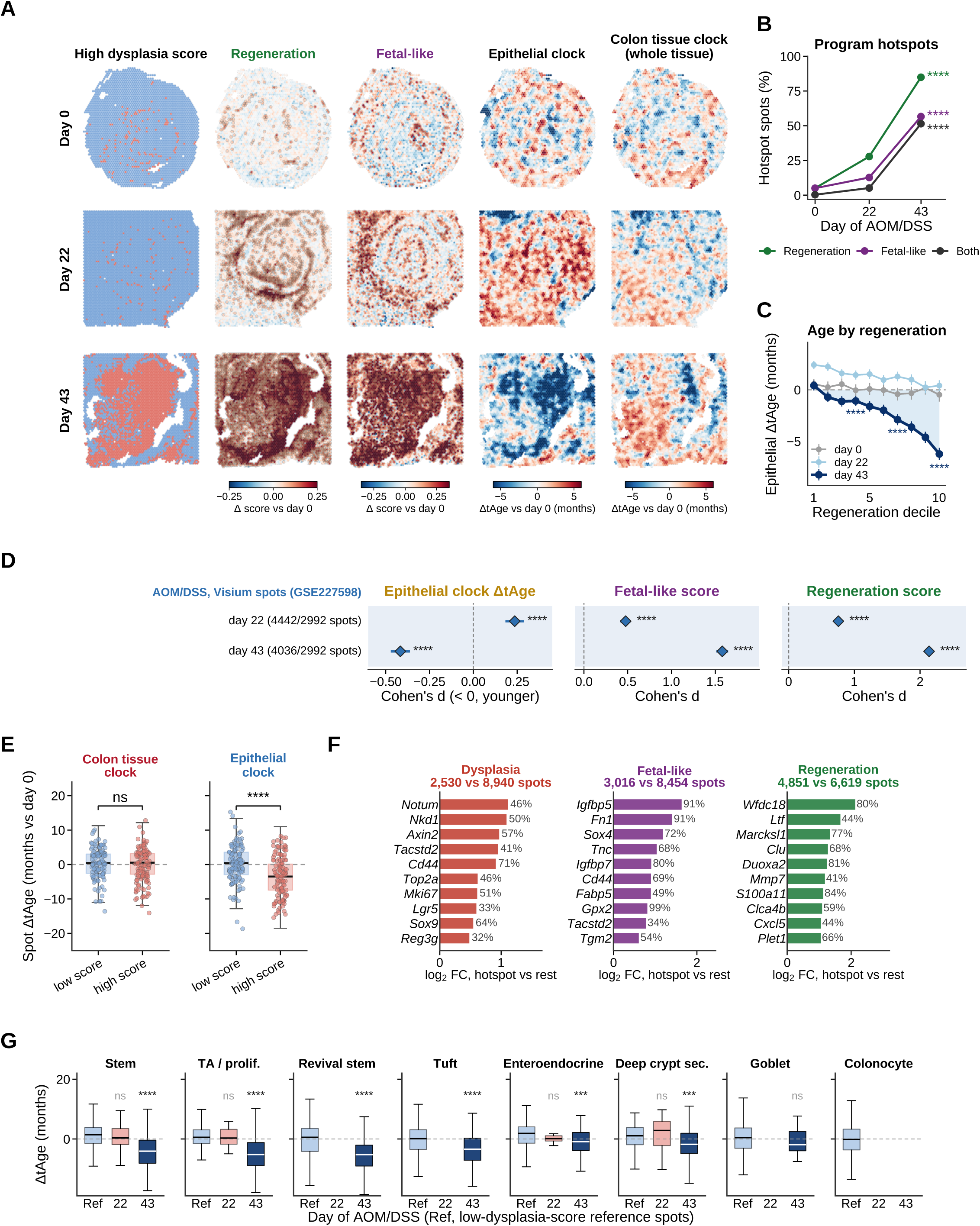
Spatial epithelial age and regenerative expression during colitis-associated tumorigenesis. **(A)** Azoxymethane and dextran sulfate sodium (AOM/DSS) colon Visium sections (GSE227598) at days 0, 22 and 43 on standard diet. Columns show high-dysplasia-score spots, regeneration and fetal-like scores, epithelial-clock age and colon tissue clock age, each referenced to day 0 (seven-nearest-neighbor smoothing for display only). The dysplasia score is expression-based and does not identify histopathologically confirmed lesions. **(B)** Proportions of spots above the day-0 95th-percentile threshold for each program (Both, spots above the thresholds for both the regeneration and fetal-like scores) on standard diet. Stars compare day 43 with day 0 (Fisher’s exact test, BH); every day-22 proportion also exceeds day 0 (q < 10 ⁵). **(C)** Mean epithelial ΔtAge by regeneration-score decile at days 0, 22 and 43 on standard diet (intervals, 1.96 times the spot-level standard error). Stars compare each day-43 decile with all day-0 spots (Mann–Whitney, BH across deciles); shading marks day-43 deciles predicted to be younger than day 0. Spearman correlations across spots were −0.05 at day 0 (2,992 spots), −0.11 at day 22 (4,442 spots) and −0.32 at day 43 (4,036 spots). **(D)** Cohen’s d with 95% confidence intervals for epithelial ΔtAge, fetal-like and regeneration scores in AOM/DSS Visium sections at days 22 and 43 versus day 0 (GSE227598; spot numbers in parentheses). Welch t test across spots. Confidence intervals are d ± 1.96 × SE with SE = √[(n₁ + n₂)/(n₁n₂) + dZ/(2(n₁ + n₂))], with spots as units, as computed in the original analysis. **(E)** Spot ΔtAge on the colon tissue and epithelial clocks in the lower and upper tertiles of the expression-based dysplasia score among epithelial-cell-type spots of the day-22 and day-43 sections on standard diet (low and high score; not a histological classification). Boxes, medians and interquartile ranges; plotted points are a display subset; Mann–Whitney comparisons use all spots. **(F)** Top marker genes of dysplasia, fetal-like and regeneration hotspots (spots at or above the 95th percentile of the day-0 section for each score; hotspot and background spot numbers above each panel, of 11,470 spots) ranked by log₂ fold change against all other spots; percentages give the fraction of hotspot spots expressing each gene. **(G)** Epithelial ΔtAge per spot, grouped by the cell type with the highest marker-gene score, in high-dysplasia-score spots of each section (days 22 and 43 on standard diet) versus low-dysplasia-score reference spots of the same dominant cell type (the lower tertile of the dysplasia score among epithelial spots of the treated day-22 and day-43 sections, pooled; Ref in the artwork). Groups with fewer than 10 spots are not drawn. Dark blue, day 43. Stars, Mann–Whitney with BH adjustment. Spots are dependent subsamples of one section per condition, so the P values compare spots within single sections and do not test differences between sections or animals.

As lesions developed, epithelial age became more inversely associated with regeneration score, with the strongest association at day 43 (**Fig. 5C**). Mean epithelial age increased slightly at day 22 before falling below baseline at day 43 (**Fig. 5D**), while the colon tissue clock showed little overall change (**Fig. 5A**). High-dysplasia-score epithelial spots were predicted to be younger by the epithelial clock, but the colon tissue clock did not distinguish them from low-scoring spots (**Fig. 5E**). The different training compartments provide one possible explanation for this divergence. The dysplasia score included regeneration, proliferation and Wnt-response genes, whose expression characterized the high-scoring regions. Fetal-like hotspots were enriched for *Igfbp5, Fn1* and *Sox4,* while regeneration hotspots included *Clu* and *Mmp7* (**Fig. 5F**). At day 43, within each of the seven epithelial types with sufficient spots for comparison, high-dysplasia spots were younger than low-scoring spots dominated by the same cell type (**Fig. 5G**).

Single-cell data provided a comparison within defined tumor-associated populations (GSE224679) (Bala *et al*, 2023). In *Apc*-mutant tumors, stem, transit-amplifying, tuft, enteroendocrine and goblet cells were predicted to be younger than enterocytes, showing that cell identity contributed to the predictions (**Extended Data Fig. 12C, Supplementary Table 4**). In the separate AOM/DSS comparison, revival stem cells from lesions were predicted to be older than those from normal tissue, while the goblet-cell difference was not significant ( **Extended Data Fig. 12D**). These analyses used bins of pooled cells from one sample per group. Together with the spatial results, they associate younger epithelial predictions with regenerative states in developing lesions, while showing that the age response is not shared by every tumor-associated population.

### Regenerative states share repair programs but differ in their relationship to aging

We compared the regenerative datasets directly to determine which expression changes they shared (**Fig. 6A**). TGF-β1-treated organoids, asporin-treated organoids and *Lgr5*-independent spheroids showed closely related responses, which overlapped partly with irradiated stem cells and enterocytes. The day-4 response to 5-FU was similar in young and old epithelium and positively associated with DSS injury and *LATS1/2* deletion, but opposed the response to *YAP* overexpression in organoids. Fetal epithelium resembled the adult regenerative states only weakly. A younger epithelial prediction thus accompanied several related repair states without requiring an identical expression program or a complete return to fetal development.

**Figure 6.**
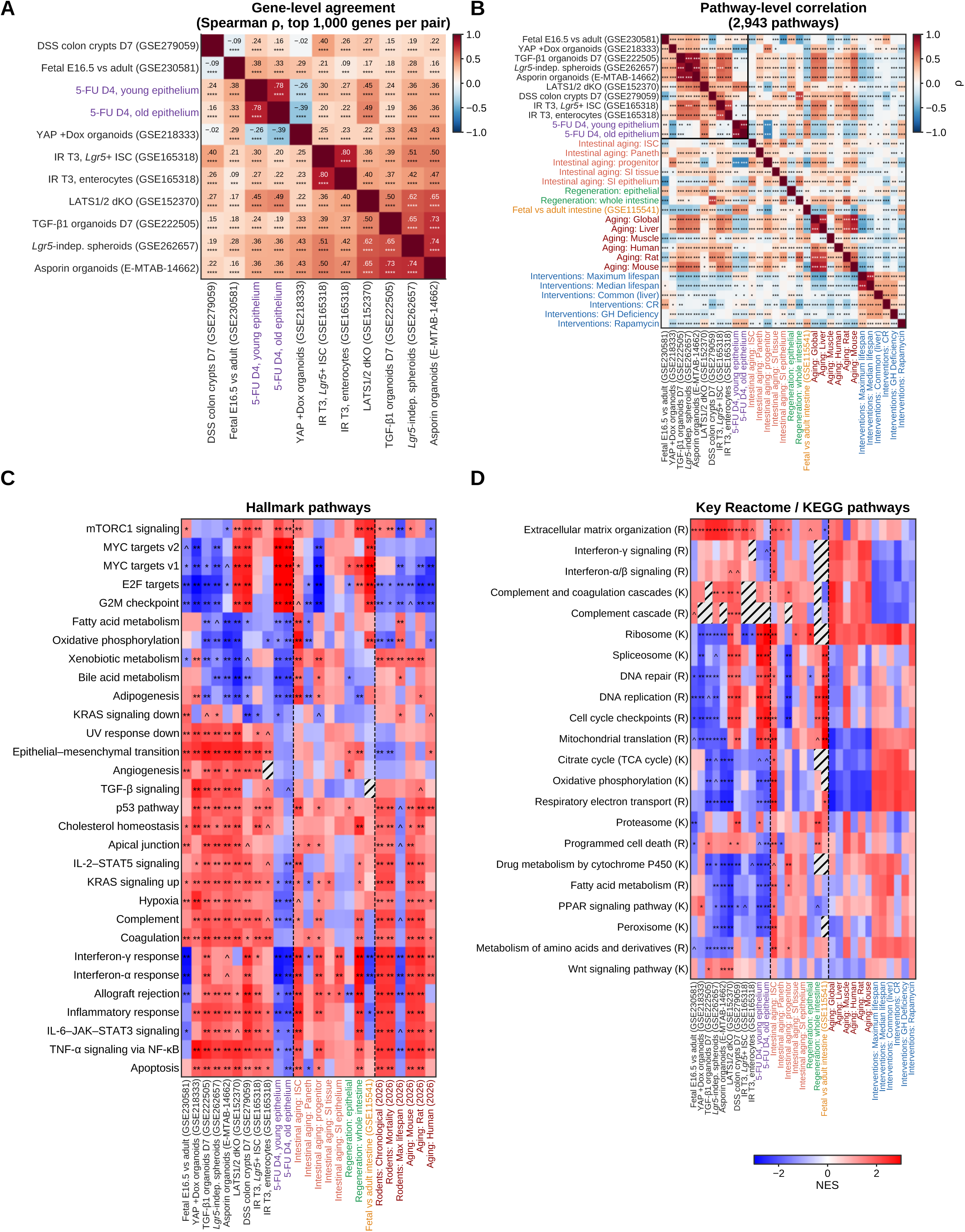
Gene- and pathway-level comparison of intestinal regenerative states with aging and longevity signatures. **(A)** Spearman correlation of gene-level log₂ fold changes among eleven contrasts over the union of the 1,000 genes with the largest absolute log₂ fold change in each pair, ordered by hierarchical clustering. The contrasts are fetal E16.5 versus adult epithelium (GSE230581), YAP overexpression (+Dox) organoids (GSE218333), TGF-β1-treated organoids at day 7 (GSE222505), *Lgr5*-independent spheroids (GSE262657), asporin-treated organoids (E-MTAB-14662), LATS1/2 double-knockout epithelium (GSE152370), DSS colonic crypts at day 7 (GSE279059), *Lgr5* stem cells and enterocytes 3 days after irradiation (GSE165318; cells of the three day-3 and four day-0 mice pooled) and pooled epithelium 4 days after 5-FU in young and old mice (two mice per group). Values, ρ; stars, BH q across pairs (the tests treat genes as independent, so the q values are descriptive and do not account for gene dependence). **(B)** Spearman correlation of pathway NES among the regenerative contrasts (boxed), intestinal aging (SI tissue and SI epithelium signatures (Methods); *Lgr5* stem cells, progenitors, Paneth cells), regeneration meta-signatures, the GSE115541 fetal-versus-adult intestine reference (Methods), and published aging and lifespan-intervention signatures, over up to 2,943 GO, Reactome and KEGG pathways. Cell values of A–D are given in Supplementary Table 5. Stars, BH q across pairs (the tests treat pathways as independent, so the q values are descriptive and do not account for genes shared between pathways). **(C)** Hallmark NES for the regenerative contrasts, intestinal meta-signatures and published rodent and species aging, mortality and maximum-lifespan signatures; the 30 sets with the largest mean absolute NES across the regenerative contrasts are shown, after omitting eight tissue-associated sets (HALLMARK_PANCREAS_BETA_CELLS, HALLMARK_SPERMATOGENESIS, HALLMARK_HEDGEHOG_SIGNALING, HALLMARK_ESTROGEN_RESPONSE_EARLY, HALLMARK_ESTROGEN_RESPONSE_LATE, HALLMARK_ANDROGEN_RESPONSE, HALLMARK_MYOGENESIS and HALLMARK_HEME_METABOLISM). **(D)** NES for selected Reactome (R) and KEGG (K) pathways across regenerative contrasts, intestinal references and published aging and intervention signatures, as labeled. Pre-ranked GSEA with 20,000 random gene sets as the null. In C and D, ** FDR < 0.01, * FDR < 0.05, ^ FDR < 0.1; hatched, pathway not evaluated; reference NES in D have no FDR. Published aging, mortality, lifespan and intervention signatures, Tyshkovskiy et al., 2019, 2023, 2026.

Their relationships to aging also differed at the pathway level (**Fig. 6B**). Most organoid responses, *LATS1/2* deletion and the day-4 5-FU response opposed the small-intestinal epithelial aging signature, while DSS injury and irradiated stem cells resembled it. Organoid and irradiation responses also shared features with multi-tissue aging, including increased inflammatory, p53 and matrix programs and reduced oxidative and fatty-acid metabolism ( **Fig. 6C,D**). The 5-FU response instead opposed several multi-tissue aging signatures, increasing cell-cycle programs while reducing interferon and TNF-α/NF-κB responses. The 5-FU response was also positively associated with rapamycin and maximum-lifespan-intervention signatures more closely than the organoid and irradiation responses (**Fig. 6B**). These findings indicate that a younger epithelial state can coexist with stress-associated features of aging, and the balance between these responses depends on the injury and experimental system.

Cell-type-resolved analysis of the 5-FU series further separated early damage from regenerative growth (**Extended Data Fig. 14A,B**). Stem cells, enterocytes and goblet cells initially increased p53 and TNF-α/NF-κB programs. Later, they increased cell-cycle and ribosomal programs and reduced interferon responses. Old goblet cells retained the early damage pattern at day 3 before making this transition. Paneth cells followed a different course, reducing several growth and metabolic programs, although their pseudobulks were small and uneven, particularly after injury. These differences place the age response within a coordinated but population-dependent progression from damage to repair.

### Liver regeneration is accompanied by younger transcriptomic predictions

To examine whether the association extends to a normally quiescent organ, we analyzed liver expression before and after partial hepatectomy in young and old mice (GSE185705, **Fig. 7A**) (Yang *et al*, 2023). We applied 24 published clocks predicting chronological age, normalized age or mortality (Tyshkovskiy *et al*, 2026). The multi-species chronological clock showed the most consistent decrease and predicted old liver to be younger at every sampled post-surgical time point, through the final observation at 240 h (**Fig. 7B**). Across the full set, most clocks shifted downward in old liver, although support varied by model (**Fig. 7C**). The average post-surgical response was significant for 13 clocks in old liver and none in young liver after correction. The direct age-by-time comparison was not significant after correction, so the stronger response in old liver does not establish an old-specific effect (**Supplementary Table 7**).

**Figure 7.**
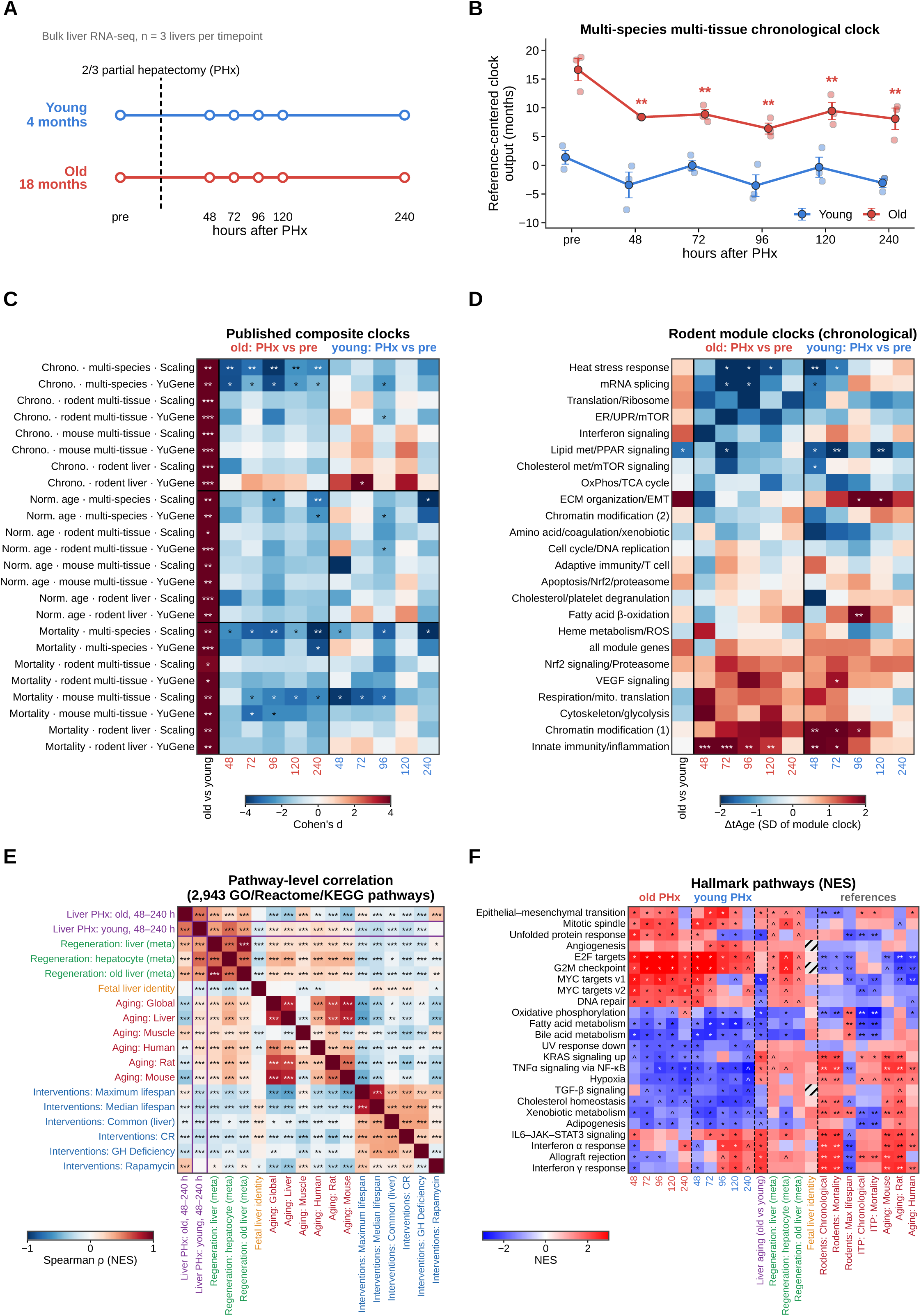
Transcriptomic age of young and old liver during regeneration after partial hepatectomy. **(A)** Design of the liver-regeneration study (GSE185705), in which livers of young (4 months) and old (18 months) mice were profiled by bulk RNA-seq before and 48, 72, 96, 120 and 240 h after two-thirds partial hepatectomy (PHx), with three livers per age group and timepoint. **(B)** Reference-centered output of the multi-species multi-tissue chronological clock (scaled input; months) for young (blue) and old (red) liver: expression is centered on young pre-PHx liver, but the clock intercept and the constant contribution of clock genes absent from the data are retained, so the output is not ΔtAge from young pre-PHx liver and the young pre-PHx mean is not zero. Post-PHx-minus-baseline differences within each age are interpretable without recalibration. Mean ± s.e.m. with individual livers (three per group). Stars, pooled-variance planned contrast versus pre-PHx liver of the same age, Benjamini–Hochberg q across the ten post-PHx contrasts of the clock (* < 0.05, ** < 0.01, *** < 0.001). This clock gave the most consistent decrease among the 24 clocks of C; every clock is shown in C and in Supplementary Table 7. **(C)** Cohen’s d for old versus young pre-PHx liver and for each post-PHx timepoint versus pre-PHx liver of the same age, for 24 published chronological, normalized-age and mortality clocks (multi-species, rodent multi-tissue, mouse multi-tissue and rodent liver; scaled and YuGene input). Stars, pooled-variance planned contrast versus pre-PHx liver of the same age, Benjamini–Hochberg q across the ten post-PHx contrasts per clock (* < 0.05, ** < 0.01, *** < 0.001); old-versus-young column, Welch P. **(D)** Change from pre-PHx liver on rodent chronological module clocks (labeled by their dominant pathways), in units of the standard deviation of each module estimate across livers. Stars, pooled-variance contrast with Benjamini–Hochberg q across the 24 rows (23 modules and the all-module-gene estimate) and ten post-PHx contrasts (* < 0.05, ** < 0.01, *** < 0.001); old versus young as in C. Module-level significance depends on the pooled-variance assumption, with Welch-test sensitivity results provided in Supplementary Table 7. **(E)** Spearman correlation of pathway NES among the PHx responses of old and young liver (48 to 240 h pooled), liver regeneration meta-signatures (aggregated lists in Supplementary Table 8; per-dataset contributors not retained), a fetal-liver identity signature (fetal liver versus other fetal tissues, labeled Fetal liver identity), and published aging and lifespan-intervention signatures over the pathways with NES in both profiles (up to 2,943 GO, Reactome and KEGG pathways; stars, BH q across pairs; the tests treat pathways as independent, so the q values are descriptive and do not account for genes shared between pathways). The old-versus-young difference of pre-surgical liver is not included because it shares reference livers with both responses (Methods). **(F)** Hallmark NES (the 24 sets with the largest absolute NES in any PHx contrast, after omitting the eight sets listed for Fig. 6C) for each PHx timepoint (hours), pre-surgical liver aging, liver meta-signatures and published rodent, Interventions Testing Program (ITP; Tyshkovskiy et al., 2026) and species signatures (** FDR < 0.01, * < 0.05, ^ < 0.1; hatched, not available). Published clocks and signatures, Tyshkovskiy et al., 2019, 2023, 2026.

Module clocks resolved different components of the hepatic response. In old liver, heat-stress, RNA-splicing and lipid-metabolism modules shifted toward younger predictions, while the innate-immunity module and one chromatin-modification module shifted toward older predictions at several time points (**Fig. 7D**). Some changes also occurred in young liver. The overall younger prediction thus combined opposing changes in molecular programs, with module-level statistical support sensitive to the variance model used for testing (**Supplementary Table 7**).

Pathway comparisons with aging signatures from independent cohorts supported a reversal of selected age-associated changes in old liver (**Fig. 7E**) (Tyshkovskiy *et al*, 2019; Tyshkovskiy *et al*, 2023). Old-liver regeneration opposed liver, mouse and multi-tissue aging signatures, while young-liver associations were weakly positive when pooled across time. Comparisons with separate gene-level aging signatures showed the same pattern (Tyshkovskiy *et al*, 2026). Both ages increased cell-cycle and DNA-repair programs and reduced hepatic metabolic programs, but inflammatory responses differed (**Fig. 7F, Extended Data Fig. 15A**). Old liver initially reduced interferon and allograft-rejection programs that were elevated in aging, whereas young liver increased these programs later in regeneration. The old-liver response also resembled the rapamycin signature, without comparable agreement with the broader lifespan-intervention signatures (**Extended Data Fig. 15B**). Mortality-module predictions likewise differed among growth, stress, immunity and matrix programs (**Extended Data Fig. 15C**), and do not directly measure survival. These findings extend the association between regeneration and younger transcriptomic states beyond the intestine, with the old-liver response persisting to the last sampled time point.

## Discussion

Restoring injured tissue requires cells to change state, and our findings indicate that some of these transitions also reverse age-associated gene expression. The younger epithelial state observed during intestinal regeneration combines repair-associated plasticity with a partial reversal of expression changes found in aging stem cells and progenitors. Its transient nature suggests that transcriptomic rejuvenation can be part of the process of rebuilding a tissue. The accompanying response in old liver extends this association beyond a continuously renewing epithelium, although its duration remains unresolved. The central biological question is which of these molecular changes enable repair and which could be retained after normal tissue organization and function are restored.

This interpretation places regeneration among physiological processes in which molecular age can change over short periods. Biological age estimates can rise during severe stress and fall during recovery (Poganik *et al*, 2023), while partial reprogramming can reduce age-associated features as cells temporarily suppress parts of their differentiated identity (Gill *et al*, 2022; Roux *et al*, 2022). Intestinal repair links these processes, where, an injury response is followed by a plastic state in which epithelial transcriptomic age decreases. The return toward baseline during recovery shows that the younger state does not necessarily become a lasting feature of the rebuilt tissue.

The pathway comparisons also show why regeneration cannot be placed on a single axis from old to young. Repair recruits stress, inflammatory and metabolic responses that overlap with aging, while reversing other age-associated changes. Reduced expression of mature metabolic genes can accompany plasticity during repair, while persistence after recovery could signal incomplete differentiation. The younger clock predictions and the variable relationships to aging signatures describe different aspects of the regenerative response.

Developmental reversion provides one explanation for this plasticity, but does not account for the entire younger state. The strong contribution of revival stem-cell and YAP-associated repair genes, together with the more limited overlap with fetal expression, places the response closer to epithelial repair than to a fully fetal state. YAP is a plausible link between the injured environment and this state because it couples matrix signals to epithelial reprogramming (Cheung *et al*, 2020; Gregorieff *et al*, 2015; Yui *et al*, 2018). Our perturbation and chromatin analyses support an association between YAP activity and the younger state. Whether the age response requires YAP remains to be tested. Manipulating its activity during distinct stages of repair could determine whether the processes that produce younger expression also promote epithelial differentiation and barrier recovery.

Cell turnover may also contribute to the younger epithelial state. A younger epithelial profile could arise through changes within surviving cells, production of their progeny or expansion of particular epithelial populations. The association with regenerative expression after adjustment for proliferation indicates that cell-cycle activity alone does not capture this response. Unequal inheritance of damaged material and organelles during division offers a possible mechanism for changing the state of daughter cells (Katajisto *et al*, 2015; Rujano *et al*, 2006). Lineage tracing combined with measurements of damage and age-associated expression would distinguish renewal within surviving lineages from changes produced by cellular replacement.

Successful repair also requires timely exit from a plastic state. Age-associated changes in Wnt signaling, Paneth-cell *Notum*, metabolism and inflammation can interfere with this progression (Mihaylova *et al*, 2018; Nalapareddy *et al*, 2017; Omrani *et al*, 2023; Pentinmikko *et al*, 2019). Sustained fetal-like responses in aged colon show how a program that supports regeneration can become maladaptive when it persists (Korcari *et al*, 2026). Our 5-FU series detected different regenerative gene responses in young and old epithelium, but did not resolve whether the decline in transcriptomic age occurred at different times. Entry into a younger state may be one component of repair, with recovery also requiring restoration of differentiated epithelial functions.

The niche may help govern both entry into and resolution of this state. TGF-β1 and mesenchymal asporin promote fetal-like epithelial reprogramming, while persistent asporin can delay its resolution in aged intestine (Chen *et al*, 2023; Iqbal *et al*, 2025). Molecular effects of injury may persist after predicted age returns to baseline. Colonic stem cells retain inflammatory chromatin memory after colitis (Nagaraja *et al*, 2026), and different intestinal injuries can leave either protection against a later insult or increased susceptibility to it (Biswas *et al*, 2026). Following tissue function after recovery and reinjury would reveal whether the younger regenerative state contributes to beneficial adaptation or accompanies a lasting vulnerability.

Tumor-associated repair further separates a younger expression state from tissue health. Developing intestinal lesions recruit regenerative plasticity (Bala *et al*, 2023; Gregorieff *et al*, 2015; Mzoughi *et al*, 2025), so younger predictions in regenerative lesion regions do not establish rejuvenation of transformed cells. The older predictions in lesion-associated revival stem cells make this distinction particularly clear. Spatial regions and defined cell populations can carry different age signals because they represent different mixtures and cellular states. Inflammatory memory can also promote subsequent tumor growth (Nagaraja *et al*, 2026). These observations argue for assessing transcriptomic age together with epithelial identity, differentiation and pathological growth when evaluating interventions that promote regeneration.

The liver findings suggest that regenerative reductions in age-associated expression are not confined to the intestine. Partial hepatectomy reverses selected age-associated H3K27me3 changes (Yang *et al*, 2023), and the accompanying younger transcriptomic predictions and opposition to independent aging signatures connect chromatin remodeling to a broader change in molecular state. The response persisted to the final sampled time point, with longer follow-up needed to determine its duration. Lower DNA methylation age after muscle regeneration and embryo-like methylation changes in regenerating hepatocytes show that repair can alter both age-associated and developmental epigenetic features (Chambers *et al*, 2025; Falick Michaeli *et al*, 2024). Whether these changes converge on sustained functional improvement remains to be established within each tissue.

Comparisons between successful and failed repair could clarify the relationship between transcriptomic age and tissue recovery. Transitional epithelial states persist in fibrotic lung, and failed-repair populations emerge after kidney injury (Kirita *et al*, 2020; Kobayashi *et al*, 2020; Strunz *et al*, 2020). These studies did not measure biological age, but demonstrate that persistence of an intermediate cell state can accompany dysfunction. Measuring age-associated expression as such cells either differentiate or remain arrested would test whether a younger transcriptomic state predicts successful rebuilding or simply identifies entry into repair.

The intestinal clocks provide tools for resolving these trajectories in the tissue and epithelial compartment. Validation across held-out datasets establishes that they detect chronological-age differences, while the separate aging signatures and sensitivity analyses help interpret their response during regeneration. The persistence of younger predictions after excluding developmental and repair gene sets in the 5-FU series indicates that the response extends beyond those annotated programs. These tools can guide comparisons across interventions, with barrier integrity, differentiated-cell output and resistance to reinjury providing the functional validation needed for an aging biomarker (Buckley *et al*, 2023; Moqri *et al*, 2024; Rutledge *et al*, 2022).

Regeneration provides a physiological setting in which adult mammalian tissues can acquire younger transcriptomic states. Longer follow-up could reveal which age-associated changes can remain reversed after rebuilding is complete. Linking those changes to sustained tissue function would establish whether transient molecular rejuvenation can contribute to lasting improvements in the repair of aged tissues.

### Limitations of the study

Several aspects of the design define the scope of these conclusions. The 5-FU series included two mice per age and sampled day (one usable old mouse at day 1 in the animal-level clock models, no day-2 samples and no young day-3 samples), and each day was processed in a separate capture, so time after injury and capture batch cannot be separated within this experiment. The irradiation time course, the replicated Visium HD time course and the other spatial datasets provide independent observations of the younger epithelial state during repair. Public irradiation analyses based on pooled cells and several spatial comparisons have limited animal-level replication, so we report mouse-level estimates separately from tests based on pooled-cell bins or spots. In the Visium HD analysis, 17 mice contributed epithelial metaspots, although both day-4 mice shared a capture area and day 6 included one mouse. Like other transcriptomic clocks, the epithelial clock responds to differentiation and cellular composition as well as chronological age, and its application to regenerating tissue extends beyond the uninjured states used for training. Its timing varied with clock and analysis choices, and the retrained clocks, input capping and alternative estimators show how far the response depends on these choices. Predictions are relative to a defined control and do not directly measure tissue function, lifespan or the age of individual cells. The aging signatures draw on the studies currently available, and pathway overlap reduces the independence of individual comparisons. The 5-FU experiment used female mice, with young and old animals from different sources, and the liver series was small, with module-level significance sensitive to the variance model and no significant age-by-time interaction after multiple-testing correction. Larger, sex-balanced time courses with animal-level replication and functional assays will determine whether the younger state is required for repair, whether YAP is necessary for it, and whether it predicts sustained functional benefit.

## Methods

### Study design and data sources

This study combined a new 5-fluorouracil (5-FU) injury time course in young and old mice with reanalysis of published expression, chromatin-accessibility and spatial transcriptomic datasets. The 5-FU series was profiled by single-cell RNA sequencing (**Figs. 1E, 3 and 6A-C, Extended Data Figs. 7-10 and 14**). Published data were obtained from GEO, ArrayExpress or CELLxGENE. Each injury or intervention was compared with a reference group from the same dataset.

### Animals

Animal procedures were approved by the Institutional Animal Care and Use Committee of Brigham and Women’s Hospital. Female C57BL/6J mice aged 4 months were purchased from Charles River Laboratories, and females aged 21 months were obtained from the National Institute on Aging Aged Rodent Colony. Mice were housed under specific-pathogen-free conditions with a 12-h light-dark cycle, ad libitum chow and water, and 7 days of acclimatization. Allocation to collection days was not formally randomized. Dissection, sorting and library preparation were not blinded, and computational analyses applied the same criteria to all groups. No mice were excluded before sequencing. Six samples were excluded for insufficient cell yield, and one further low-quality sample was excluded from animal-level clock models, as described below.

### 5-fluorouracil injury

5-FU (Sigma-Aldrich, F6627) was dissolved in dimethyl sulfoxide (DMSO) at 100 mg/ml and administered once intraperitoneally at 200 mg/kg before 9 a.m. DMSO-injected controls formed the day-0 group. Small intestines were collected on days 1-6 after treatment, with two young and two old mice assigned to each day and to day 0 (28 mice total).

### Isolation of small-intestinal epithelial cells

Single-cell suspensions were prepared as described previously (Pentinmikko et al, 2019). Briefly, the small intestine was flushed with cold PBS, opened longitudinally, cleared of mucus and cut into small fragments, which were incubated in 10 mM EDTA in PBS on ice. The epithelium was released by vigorous shaking, washed in cold PBS and dissociated into single cells in TrypLE Express (Gibco) containing DNase I (1,000 U/ml; Roche). Cells were stained for 30 min on ice with anti-EpCAM-APC (clone G8.8; eBioscience, 17-5791-80), anti-CD31-PE-Cy7 (clone 390; eBioscience, 25-0311-81) and anti-CD45-PE-Cy7 (clone 30-F11; BD Pharmingen, 561868), and 4′,6-diamidino-2-phenylindole (DAPI; 1 µM; Sigma-Aldrich) was added to exclude dead cells. Live (DAPI-) EpCAM+CD31-CD45-epithelial cells were sorted on a Cytek Aurora CS cell sorter (Cytek Biosciences). Cells sorted from the four mice collected on each day were processed for single-cell capture on the same day.

### Single-cell RNA sequencing

Single-cell libraries were prepared with the 10x Genomics Chromium GEM-X Universal 3′ Gene Expression v4 kit with On-Chip Multiplexing (OCM; 4-plex, 16 samples, PN-1000779) by Athenomics LLC (Framingham, MA). On each collection day, cells from the two young and two old mice were loaded into the four sample inlets of one OCM channel, so that each mouse received its own OCM barcode, with a target recovery of 5,000 cells per mouse. Libraries were sequenced on an Illumina NovaSeq X Plus with a NovaSeq X Series 25B reagent kit (300 cycles; 20104706) as paired-end 150-bp reads with 8-bp dual index reads (the run was shared with other libraries), yielding 380 to 705 million reads per daily run. Reads were processed with the Cell Ranger multi pipeline (v10.1.0, 10x Genomics) against the 10x Genomics mouse reference refdata-gex-GRCm39-2024-A, with default cell calling (no expected cell number specified), default assignment of cells to OCM barcodes (OB1 to OB4), which also removes multiplets formed between samples, and default inclusion of intronic reads. Cell Ranger called 57,880 cells across the 28 samples.

### Processing of the 5-FU single-cell data

Counts were processed in Scanpy (Wolf *et al*, 2018). Within each sample, cells were excluded for values more than five median absolute deviations from the median in log total counts, log detected genes or the fraction of counts in the 20 most expressed genes, for more than 50% mitochondrial counts, for fewer than 200 detected genes, or for a Scrublet doublet call (Wolock *et al*, 2019). The mitochondrial threshold retained mitochondria-rich enterocytes. Of 57,880 called cells in the 28 libraries, 44,815 passed these cell filters, with 1,297 doublets removed and 24,179 genes retained; 44,792 of the retained cells belonged to the 22 samples analyzed. Per-sample quality summaries give medians of detected genes, UMIs and mitochondrial fraction over all called cells before filtering, whereas cell numbers are counts after all filters, including the 50% mitochondrial filter (Supplementary Table 4). Excluding samples with fewer than 50 passing cells removed all four day-2 samples and both young day-3 samples. Before cell filtering, day-2 libraries had medians of 64-348 detected genes per cell and 47-73% of reads in cells, compared with approximately 1,900-2,800 genes and 79-84% at day 0. The remaining series contained 22 samples, with two mice per age on days 0, 1, 4, 5 and 6 and two old mice on day 3. Passing cell counts ranged from 205 to 6,613 per mouse.

Counts were normalized to 10,000 per cell and log-transformed. The 2,000 highly variable genes were selected from raw counts with seurat_v3 and sample as batch, scaled with clipping at 10, and used for PCA (50 components, arpack). Harmony integrated samples by identity (harmonypy, 30 iterations, Korsunsky *et al*, 2019). A graph of 15 neighbors over 40 Harmony components supported UMAP (minimum distance 0.3, seed 0) and Leiden clustering (resolution 1.0, igraph, two iterations, seed 0). Clusters were initially assigned by the highest marker-gene score, z-scored across clusters. Labels were transferred from GSE165318 (22,680 cells, Lu *et al*, 2023) using Scanpy ingest over 17,196 shared genes. Marker-defined endothelial, smooth-muscle, mast and immune labels were retained because these populations were poorly represented or absent in the reference. Non-epithelial cells were excluded from epithelial analyses. Stem and transit-amplifying cells were subclustered with 15 neighbors on the Harmony embedding and Leiden resolution 0.8. Subclusters were assigned as stem cells when the mean stemness score (*Lgr5*, *Ascl2*, *Olfm4*, *Smoc2*, *Slc12a2*), z-scored across subclusters, equaled or exceeded the proliferation score (*Mki67*, *Top2a*, *Ccnb1*, *Birc5*, *Ube2c*, *Cenpa*, *Cenpf*, *Cdk1*), and otherwise as transit-amplifying cells. Epithelial types were stem, transit-amplifying, enterocyte, goblet, Paneth, enteroendocrine, tuft and revival stem cells. *Ly6a*-positive cells contained at least one *Ly6a* UMI, and *Clu* detection was assessed per metacell. Integration was used for annotation only. Expression analyses used raw counts.

For pooled-epithelium analyses, raw UMI counts were summed in non-overlapping random bins formed separately within each mouse. These bins are termed metacells here. Cells were permuted with NumPy default_rng seed 0 and divided into floor(n/35) equal bins, distributing remainders and discarding bins with fewer than 25 cells (1,114 metacells of 35-43 cells). Cell-type analyses used the same procedure within each mouse and cell type, targeting 15 cells per bin with a minimum of 8 and a maximum of 120 bins (1,612 metacells of 10-28 cells). For **Fig. 1E** only, day-0 cells of each type were pooled across the two mice of each age before binning, targeting 30 cells with a minimum of 15 (389 bins of 26-35 cells). Per-mouse pseudobulks were formed by summing raw counts across all epithelial cells or across each cell type. Random partitions throughout the study used seed 0 unless specified otherwise.

### Public datasets

Deposited count, FPKM and RPKM matrices were downloaded. Ensembl and Entrez identifiers were mapped to mouse gene symbols and Entrez identifiers with NCBI gene_info, and duplicate identifiers were summed. Single-cell counts were summed to pseudobulk profiles. When animal-level profiles were unavailable, cells were partitioned into random bins, generally four per condition for pooled epithelium or up to three bins containing at least 15 cells per cell type and sample. Dataset-specific partitions and biological replication are described below and in Supplementary Table 2. These bins were treated as subsamples of the source material. Microarray data (E-MTAB-5249, Affymetrix Mouse Gene 2.0 ST) were RMA-normalized with oligo (Carvalho & Irizarry, 2010) and analyzed using signature scores.

### Intestinal transcriptomic clocks

#### Training data

The small-intestine (SI) tissue clock used 465 profiles from eight studies: 421 whole-tissue or segment profiles, 24 crypt profiles (GSE129713), 10 villus-fraction profiles (GSE182252) and 10 per-animal ileal single-cell pseudobulks (GSE288291). The eight accessions were GSE314817, GSE132040 (Schaum *et al*, 2020), GSE213722, GSE129713, GSE182252, GSE234563, GSE288291 and GSE274687. The colon tissue clock used 252 profiles from seven studies: GSE314817, GSE213722, the GSE272841 control arm, young and aged controls from GSE280382, GSE248002, GSE274687 and Tabula Muris Senis large-intestine single-cell data (Tabula Muris, 2020). Only control genotypes and untreated animals were included, with single-cell profiles summed per animal and compartment. The pan-intestinal clock used the union of these sets (717 profiles across 22 series). The model validated in **Fig. 1C** was used in every pan-intestinal application. Tissue-clock leave-one-study-out validation used 12 folds because Tabula Muris Senis bulk and single-cell data were kept separate, although they originate from one study. Crypts, GSE288291 and Tabula Muris Senis large-intestine data also contributed to epithelial-clock training. Training series and overlap are listed in Supplementary Table 6.

The epithelial clock used one profile per animal from 178 animals in 14 studies. Sorted or purified profiles included *Lgr5*-high stem cells, *Lgr5*-low progenitors, Paneth cells, enterocytes, crypts and epithelium from GSE129713/GSE129510, GSE210240, GSE122441, GSE84061, GSE190082, GSE198137, GSE122230 (ad libitum arm), GSE262228 (young and old arm), GSE261216, GSE285985 and E-MTAB-7916 (Pentinmikko *et al*, 2019). Single-cell data from GSE200536, GSE210241, GSE288290, GSE288291 and Tabula Muris Senis large intestine were summed per animal. Libraries from the same animal were averaged, producing profiles that could combine epithelial populations. Subseries from one publication were assigned to the same study, including GSE129713/GSE129510, GSE210240/GSE210241 and GSE288290/GSE288291. GSE122230 and GSE200536 reported young and old groups without exact ages and were assigned 3 and 22 months. E-MTAB-7916 reported ages of 3-9 months and over 24 months and was assigned 6 and 24 months. Applications to individual cell types use a composition different from the training profiles and were interpreted relative to a matched cell-type reference.

#### Model fitting

Clocks predicted age relative to a reference group within each dataset and tissue, following Tyshkovskiy *et al* (2026). The youngest group of each training series supplied the reference expression profile, and the target was (chronological age minus reference age)/48 months, using a maximum recorded mouse lifespan of 4 years. Predictions were multiplied by 48 to express differences on a month scale. For tissue clocks, counts were normalized to counts per million, YuGene-transformed (Lê Cao et al, 2014) on these unlogged values over all genes of each profile (in training, the 13,015 genes measured in every series), and re-expressed as log2(1 + 10⁴ × YuGene). The 4,000 most variable genes were standardized. ElasticNet selected features by five-fold cross-validation over l1 ratios of 0.3, 0.5, 0.7 and 0.9 and 25 penalty values, followed by BayesianRidge fitting on genes with non-zero coefficients (Zou & Hastie, 2005). The selected l1 ratio was 0.3 for all tissue clocks. Penalties and selected-gene counts were 0.0023 and 205 for SI, 0.0015 and 146 for colon, and 0.00037 and 550 for the pan-intestinal model.

The implementation differed from Tyshkovskiy *et al* (2026) in using counts-per-million normalization, variance-based gene preselection, the stated ElasticNet grid and a log2(1 + 10⁴ × YuGene) transformation before standardization. Training and query profiles underwent the same sequence of transformations. In the deposited code (build_si_colon_v2.py for the tissue clocks; build_epithelial_v3e_yugene.py and apply_epithelial_v3ey.py for the epithelial clock), log2(counts per million + 1) values are converted back to counts per million before YuGene, so YuGene is computed on unlogged values and the logarithm is applied only afterwards. Because YuGene divides each value by the sample total before ranking, a per-sample scale factor cancels: for 12 training profiles (GSE274687 pseudobulks), inputs computed with relative-log-expression size factors (0.18 to 3.9) instead of counts per million differed by at most 3 × 10⁻¹Z, whereas applying YuGene after the logarithm would have made them differ by up to 0.06 (c66_yugene_scale_invariance.py).

For the epithelial clock, 12,762 genes measured across all training studies were merged. The 9,670 detected in at least 80% of profiles formed the normalization panel, and profiles detecting fewer than 60% of merged genes were removed before YuGene transformation. BayesianRidge was fitted without ElasticNet preselection. Grouped cross-validation, with animals as groups, selected 1,000, 2,000 or 4,000 variable genes and gamma hyperparameters α₁ = α₂ and λ₁ = λ₂, each set to 10⁻⁶ or 10⁻Z. The final model retained 2,000 genes. Sensitivity analyses used an ElasticNet estimator and a quantile-normalized input, mapping each profile to the mean sorted training expression profile.

#### Validation

Performance was assessed by pooled five-fold cross-validation and leave-one-study-out validation. Tissue clocks were also evaluated with ElasticNet as the final estimator (**Extended Data Fig. 1C,D**). Within each tissue-clock training fold, variable-gene selection, standardization, ElasticNet tuning with study-grouped inner folds, feature selection and BayesianRidge fitting were repeated. Epithelial-clock gene-panel size, priors, standardization and coefficients were selected within each training fold. Gene and profile filters that use detection but not age were applied once to all training profiles before the outer folds and were not recomputed within folds: for the epithelial clock, the 12,762 genes measured in all training studies, the 9,670 detected in at least 80% of profiles (the YuGene panel) and the removal of profiles detecting fewer than 60% of genes; for the tissue clocks, the 13,015 genes measured in every series. Held-out profiles therefore contributed to these detection filters but not to variable-gene selection, standardization or model fitting. Each held-out study was centered on its own youngest group, so validation measures how well age differences within a study are recovered relative to a reference group of that study, not the absolute age of a sample without a reference. Reference sensitivity was assessed in ten runs using a randomly selected half of each series’ youngest group (seeds 0-9), with the remaining young animals retained as ordinary training or test observations. This procedure was applied to the epithelial and pan-intestinal clocks within their nested validation schemes. Pearson r, RZ and mean absolute error were reported on the month scale, alongside the error of a null model predicting the training-fold mean (Tyshkovskiy *et al*, 2026). For studies with two ages, old-minus-young differences, ROC area and error relative to the null were also reported because correlation primarily measures group separation. Validation and reference-sensitivity results are provided in Supplementary Table 6.

To test dependence on developmental and regenerative genes, the epithelial clock was retrained after excluding genes differentially expressed between E16.5 and adult SI epithelium (GSE230581, DESeq2 q < 0.05 in either direction, 4,676 candidate genes), the top 300 epithelial regeneration genes, the curated repair and fetal/revival sets, and both younger-state signatures. In total, 4,922 of 9,670 candidate genes were excluded, leaving 4,748. Normalization, training design and nested validation matched the original procedure. A control model retrained on all candidate genes within the same pipeline selected 4,000 variable genes, compared with 2,000 in the original model. Two further exclusion analyses removed only fetal-upregulated genes or all fetal-versus-adult genes. Models were applied to 5-FU per-mouse pseudobulks and pooled-epithelium, stem-cell and enterocyte metacells, referenced to day 0 within age and population, and to irradiation pooled-epithelium bins (24 per day) and cell-type bins (approximately 30 cells). Validation and per-mouse analyses are in Supplementary Table 6. To assess extrapolation by the original clock, normalized, standardized and day-0-centered 5-FU inputs were compared with the 178 training profiles. We calculated Mahalanobis distances in the first 20 training principal components, expressed these as percentiles of training distances, and counted clock genes outside their training range. Predictions were then repeated after capping each gene at its training minimum and maximum. Day contrasts used Mann–Whitney tests on metacells, BH-adjusted within age and readout, and linear models adjusted for log10 UMIs on per-mouse profiles.

Because leave-one-study-out performance informed model development, the final epithelial clock was also applied without modification to four aging studies not used to train, tune or choose any epithelial-clock version: GSE210669 (sorted SI epithelium, 5 and 24 months, three mice per age), GSE274687 (three young and three old mice), GSE182252 (villous epithelium, four young and six aged mice) and GSE213722 (upper and lower SI epithelium at 4, 12 and 24 months). Each was referenced to its own youngest group, with mice as units. GSE274687, GSE182252 and GSE213722 contributed to tissue-clock training and provide independent evaluation only of the epithelial clock. GSE210669 originated from the Choi *et al* (2023) study, independently of the similarly numbered GSE210240/GSE210241 training subseries from SuperSeries GSE210242. Results are reported in Supplementary Table 6.

### Application of the clocks

Query counts were mapped to Entrez identifiers, normalized to counts per million and transformed as in training: YuGene on the unlogged values, over the 9,670-gene training panel for the epithelial clock and over all genes of the query profile for the tissue clocks, followed by log2(1 + 10⁴ × YuGene). Each gene was standardized with the training mean and standard deviation, the mean of the dataset’s reference group was subtracted, clock genes absent from the query were set to zero (equivalent to imputing the reference-group mean), and the model prediction was multiplied by 48. Estimates are differences in months from a reference group within the same dataset (ΔtAge). They are interpreted as relative transcriptomic age, not as absolute age or as a gain or loss of lifespan, and they require a defined reference cohort. Fetal and neonatal samples lie outside the adult training range and were interpreted only as a relative ordering. The epithelial clock (YuGene input, BayesianRidge estimator) was used as the primary clock for epithelial profiles. The SI tissue, colon tissue and pan-intestinal clocks, the ElasticNet estimator and quantile-normalized input were secondary analyses, and disagreements with the primary clock are reported.

### Evaluation in additional aging cohorts

Additional aging and intervention cohorts were scored relative to their own young or control group (**Extended Data Fig. 1E-H**). Cohorts included in a clock’s training data (†), or from the same publication (‡), were marked and excluded from its independent validation. Cross-compartment applications were interpreted as reference-relative differences within that dataset. The seven cohorts in **Extended Data Fig. 1H** were GSE280382 (young and aged colon controls), GSE288285 (colon), GSE100701 (colon, ad libitum arm), GSE210669 (SI epithelium), GSE301362 (SI organoids, two per group), GSE193093 (large intestine, two per group) and GSE225576 (eight non-intestinal tissues). GSE210669 cells were summed per mouse and epithelial type (three mice per group). Planned age and treatment contrasts used pooled-variance t statistics with two-sided label-permutation P values and BH adjustment, with age-by-treatment effects estimated as differences in differences. The aging summary in **Extended Data Fig. 1H** used Welch tests adjusted within clock, displaying significance only for groups with at least three samples. All intervention contrasts and BH adjustments within study and clock are listed in Supplementary Table 3. GSE288285 shares a publication with epithelial training datasets GSE288290 and GSE288291 and is not an independent epithelial-clock test (Korcari *et al*, 2026).

### Regenerative and developmental comparisons

The epithelial clock was applied to freshly sorted fetal E16.5 versus adult proximal SI epithelium (GSE230581, Pikkupeura *et al*, 2023), neonatal necrotizing enterocolitis versus control epithelium (GSE278973, Xiong *et al*, 2026), and nine regenerative comparisons: TGF-β1-treated versus vehicle organoids at day 7 (GSE222505, Chen *et al*, 2023), YAP overexpression (GSE218333), LATS1/2 deletion versus control colon epithelium (GSE152370), DSS colitis (GSE279059, Xiao *et al*, 2025), *Lgr5*-independent spheroids versus organoids (GSE262657, Marefati *et al*, 2025), organoid self-organization (GSE329635, Schwayer *et al*, 2026), refeeding after fasting (GSE192482, Imada *et al*, 2024), irradiation day 3 versus day 0 within epithelial cell type (GSE165318, Lu *et al*, 2023), and asporin versus vehicle organoids (E-MTAB-14662, Iqbal *et al*, 2025). The GSE230581 comparison used the sorted epithelial samples GSM7227026–GSM7227028 (E16.5, EpCAM⁺CD31⁻CD45⁻) and GSM7227023–GSM7227025 (adult, Lin⁻EpCAM⁺CD24-mid), not the fetal organoid samples of the same series. For time courses, **Fig. 1H** shows the time point with the highest regeneration score, selected without using clock predictions. Complete trajectories are in Supplementary Table 2 and **Extended Data Figs. 2 and 3**. Differences in **Fig. 1H** are mean epithelial-clock predictions relative to controls. Intervals are 95% Welch–Satterthwaite intervals except for the paired asporin comparison, and no interval is shown for NEC. GSE152370 bins were averaged within mouse (two per group), GSE262657 profiles within donor (three spheroid donors in three media and two organoid donors), and asporin was analyzed as four donor-matched differences using a paired t interval and test. Other contrasts used Welch tests, with BH adjustment across the 11 comparisons. Where all bins of a condition came from one library (GSE329635 and GSE192482, one sample per time point or condition; GSE165318, the pooled cells of three or four mice per day; GSE278973, the pooled cells of two mice per group), the Welch intervals and tests are nominal, are retained as calculated and quantify variation among bins of that library, not variation between animals or cultures.

The GSE222505 comparison of 24 versus 0 h after re-plating was excluded from **Fig. 1H** because TGF-β1 exposure was confounded with culture time. Other figures report Cohen’s d using the square root of the mean group variances, or a degrees-of-freedom-weighted pooled standard deviation. Confidence intervals were d ± 1.96 × SE, with SE = √[(n₁ + n₂)/(n₁n₂) + dZ/(2(n₁ + n₂))]. Asporin used the paired effect d_z. No pooled effect was estimated across datasets. Cell-identity effects were examined in per-mouse pseudobulks from the four uninjured in-house mice, retaining cell types with at least 30 cells and 200,000 UMIs, using pooled young epithelium as reference and cell type and age as model factors. Descriptive gene contributions were approximated as the log2 fold change of each clock gene in log2 counts per million between the regenerative state and its control, multiplied by the BayesianRidge coefficient divided by the training standard deviation of that gene’s log2(1 + 10⁴ × YuGene) input, and by 48 to give months. This approximation uses ordinary log2 fold changes rather than changes in the standardized YuGene inputs and ignores the nonlinearity and panel dependence of YuGene, so it does not decompose predicted age exactly; summed over all measured clock genes it correlated with the observed shifts (r = 0.99; eight contrasts with an observed shift in Supplementary Table 4) but reached a median of 55% (45% to 74%) of their magnitude. Identity genes differed between enterocyte and stem-cell pseudobulks of the four uninjured in-house mice (cell types with at least 30 cells; limma with mouse as block; FDR < 0.05 and absolute log2 fold change > 1). The identity share is the summed contribution of these genes divided by the summed contribution of all measured clock genes (c50_clock_contributions.py). Regenerative contrasts were also adjusted for an enterocyte score, the mean z score of 20 mature-enterocyte genes including *Alpi*, *Apoa4*, *Fabp2*, *Lct*, *Sis* and *Slc5a1*, and assessed within cell type in GSE165318.

### Intestinal gene signatures

#### Aging meta-signatures

Young-versus-old datasets were analyzed within study with limma (Ritchie *et al*, 2015). The *Lgr5*-high stem-cell meta-signature combined GSE122230 (Mihaylova *et al*, 2018), GSE198088/GSE198137 (Nefzger *et al*, 2022), GSE210240, GSE262228 and the corresponding cells from Pentinmikko *et al* (2019). *Lgr5*-low progenitor and Paneth-cell signatures combined GSE122230, GSE210240 and Pentinmikko *et al* (2019). For these sorted-cell signatures, the standard error of each study was derived from its log2 fold change and moderated limma P value as SE = |log2 fold change|/z, with z the upper P/2 quantile of the standard normal distribution (two-sided P bounded below at 10⁻3⁰⁰; no degrees of freedom are used, so for studies with few residual degrees of freedom this SE is larger than the moderated t-based SE; c1_correlations.py), and effects were combined per gene by random-effects meta-analysis with restricted maximum likelihood (Tyshkovskiy *et al*, 2026). Genes required measurements in at least two studies, with BH adjustment of meta-analysis P values. Tissue-level signatures included in-vivo mouse SI or mucosal, epithelial or crypt profiles from untreated or control-genotype animals, at least two independent samples per age group, a youngest group at most 6 months old and an oldest group at least 18 months old. Supplementary Table 1 lists candidates and exclusions. The SI tissue signature combined GSE39974 mucosa, GSE314817 intestinal segments, GSE132040 bulk SI, GSE182252 whole ileum and GSE234563 control intestine. The SI epithelial signature combined GSE190082 (Omrani *et al*, 2023), GSE213722, GSE129708 crypts, GSE182252 villous epithelium and per-animal epithelial pseudobulks from GSE210669, GSE210241, GSE274687 and GSE288291. Of these thirteen source profiles, eleven also contributed to clock training. GSE39974 and GSE210669 did not. Genes with at least 10 reads in 25% of samples were RLE-normalized and log-transformed. Within-study limma models related expression to age in months, adjusting for sex, segment, compartment or diet where applicable and blocking by mouse or pool when it contributed several profiles. Animals younger than 2 months were excluded. GSE314817 segments could not be linked to individual animals and were fitted separately, with their estimates averaged. Effects per 12 months were combined as above, using each study’s limma coefficient and moderated standard error directly (no conversion from P values).

#### Regeneration and developmental references

The epithelial regeneration meta-signature combined gene ranks from intestinal epithelial regeneration datasets using robust rank aggregation (Kolde *et al*, 2012), with signed −log10 aggregation scores as weights. Its candidate roster of 16 accessions included GSE165318 and GSE222505, but the per-dataset ranked lists, the contrasts and the exact contributing set were not retained and could not be recovered, and no gene met adjusted significance in the aggregation (smallest adjusted P = 1). The candidate roster and the complete aggregated list with scores and adjusted P values are given in Supplementary Tables 5 and 8. Scores derived from this signature in those datasets are descriptive. GSE279059, GSE131032, GSE152369, GSE152370, GSE262657, GSE329635 and GSE192482 were absent from the roster. Sensitivity analyses used 26 curated injury-repair/YAP-target genes and a 22-gene composite fetal/revival marker panel, both assembled independently of GSE165318 and the 5-FU series (Supplementary Table 5). The composite panel combines revival stem-cell markers (Ayyaz et al, 2019), covering 15 genes of the curated 25-gene revival set used for overlap tests, with fetal and repair-associated epithelial genes (*Spp1*, *Tnc*, *Krt8*, *Krt19*, *Sprr2a3*, *Anxa10* and *Gsdmc2*). It is not a list of revival stem-cell genes reported by Ayyaz et al (2019), and per-gene sources were not recorded. One unmatched, truncated RIKEN identifier was excluded from the source list. The whole-tissue regeneration reference combined DSS day 7 versus control from GSE131032 whole colon (Czarnewski *et al*, 2019) and GSE279059 colonic crypts (Xiao *et al*, 2025). Its 6,697 genes changed in the same direction in both contrasts and were weighted by the signed mean −log10 adjusted P. Comparisons of these discovery contrasts with the reference are marked in **Fig. 1I**. The fetal-versus-adult reference compared freshly sorted E16.5 (GSM7227026–GSM7227028) and adult (GSM7227023–GSM7227025) proximal SI epithelium in GSE230581 using DESeq2 (13,890 genes), with log2 fold changes as weights (Love *et al*, 2014, Pikkupeura *et al*, 2023). The fetal spheroid reference was mapped from microarray probes to gene symbols and weighted by log2 ratios of mean expression (Mustata *et al*, 2013). A second fetal reference, used for the overlap tests (**Fig. 1K**, **Extended Data Fig. 5D**), the fetal-like scores of **Extended Data Fig. 3**, the epithelial-metaspot fetal-like scores (**Fig. 4F**, **Extended Data Fig. 11B**) and the fetal-versus-adult intestine pathway reference (**Figs. 2E, 3E and 6B–D** and **Extended Data Fig. 10**), compared whole fetal small intestine (E12 to E18, eight samples) with adult ileum (two samples) in GSE115541 (counts per million, genes with a mean of more than one count per million in either group, Welch t test with BH adjustment, FDR < 0.05 and absolute log2 fold change > 1.5; 3,733 fetal-higher and 1,708 adult-higher genes). Gene sets used its 300 or 1,000 fetal-higher genes with the largest log2 fold changes, and the pathway ranking was the signed −log10 P of these 5,441 genes. This reference differs from GSE230581 in tissue, stages and adult sample size and agrees poorly with it (correlation of log2 fold changes r = 0.08); 426 of its fetal-higher genes have no adult counts, consistent with an annotation-mapping artifact, so scores based on it may partly reflect library depth. Results based on it are reported as computed (Supplementary Table 8).

#### Signature scores

For time courses, the fetal-like score was the mean z score of log2 counts per million over the 300 genes with the largest Wald statistics among genes increased in E16.5 relative to adult SI epithelium (adjusted P < 0.05, log2 fold change > 1, base mean > 50), and the regeneration score was the corresponding mean over the 300 highest-ranked increased genes of the epithelial regeneration meta-signature. Z scores were computed across the units of each series, population and age and centered on day 0.

### Comparisons with aging signatures

Seven contrasts were compared with intestinal aging, regeneration and developmental signatures (**Fig. 1I, Extended Data Fig. 4A,B**): TGF-β1 versus vehicle in organoids from uninjured intestine at day 7 and irradiated intestine at day 11 (GSE222505, three samples per group, Chen *et al*, 2023), DSS day 7 versus baseline whole colon (GSE131032, Czarnewski *et al*, 2019) or control colonic crypts (GSE279059, three per group, Xiao *et al*, 2025), LGK974 versus vehicle in EpCAM+ E16.5 epithelium matched by proximal, middle and distal regions (GSE183532, Maimets *et al*, 2022), spheroid-score-high epithelial cells at irradiation day 3 versus baseline epithelial cells below the baseline median of both the spheroid score and a score of 490 strongly fetal-higher genes (log2 fold change > 4) of the GSE115541 reference described above (GSE165318, 333 versus 585 cells, Lu *et al*, 2023), and donor-matched asporin versus vehicle organoids (E-MTAB-14662, four female mice, Iqbal *et al*, 2025). Spheroid-score-high cells exceeded the baseline 97.5th percentile of the Mustata *et al* (2013) score. GSE230581 organoid data were excluded because the deposited count matrix lacked gene identifiers. Sorted-cell aging signatures share training datasets with the epithelial clock, so these signature comparisons provide an analysis distinct from clock prediction but do not constitute an independent aging-data collection.

For count data, genes with at least one count per million in two or more samples were retained and expressed as log2(counts per million + 1). FPKM and RPKM matrices were analyzed as log2(value + 1) for genes with a mean of at least 1, and the deposited GSE279059 values, already log2-transformed FPKM, were used without further transformation. Log2 fold changes were differences in group means, or means of within-pair differences for asporin and of region-matched differences for LGK974. For enrichment, genes were ranked by a moderated t statistic with empirical Bayes variance shrinkage as in limma, using a one-sample paired statistic for asporin. The unreplicated GSE165318 contrast was ranked by log2 fold change. Spearman correlations between contrast log2 fold changes and signature weights were computed over the union of the 1,000 most significant genes of each pair (Tyshkovskiy *et al*, 2026); the plotted BH-adjusted correlation P values treat genes as independent and are nominal because genes are co-regulated. Adjustment covered the 56 reference–contrast pairs in **Fig. 1I**.

### Younger-state signature

The younger-state signature used nine regenerative comparisons from **Fig. 1H and t**he fetal-versus-adult comparison. For time courses, the time point with the lowest epithelial-clock estimate was used. Because this rule uses the clock, associations of the signature with epithelial-clock predictions are not independent of its construction. The clock-independent rule of **Fig. 1H** changes only GSE329635 (36 h instead of 60 h); with it, 570 of the 578 upregulated and 543 of the 550 downregulated genes were retained (Jaccard similarity 0.92 and 0.95; Supplementary Table 5). The confounded GSE222505 24-h re-plating contrast was excluded. Log2 fold changes were differences in mean log2 counts per million. Upregulated genes increased at least 1.5-fold in at least 60% of measured contrasts and were measured in at least seven. The downregulated set used the corresponding decrease, with genes ranked by recurrence and then mean effect. The resulting lists contained 578 upregulated and 550 downregulated genes. Gene symbols were matched directly. As a sensitivity check, mapping official symbols and unique synonyms to NCBI Gene identifiers merged aliases such as *Ctgf*/*Ccn2* and *Cyr61*/*Ccn1*, adding 23 upregulated and 28 downregulated genes and removing 5, with Jaccard similarity of 0.97 for each list (Supplementary Table 5). Analyses use the original symbol-matched lists. Full gene-level membership, per-contrast log2 fold changes, the time point used for each contrast and the other analyses that reuse the same data are given in Supplementary Table 5. Hallmark enrichment used pre-ranked GSEA with 1,000 permutations and seed 42. CollecTRI transcription-factor and PROGENy pathway activities were estimated from contrast log2 fold changes using decoupleR 2.1.6, respectively with univariate and multivariate linear models. PROGENy used target genes with adjusted P < 0.05 for each pathway. Heat maps display 22 terms with the largest activities and nominal per-contrast P values (Badia-i-Mompel *et al*, 2022, Müller-Dott *et al*, 2023, Schubert *et al*, 2018).

GSE66567 (YAP overexpression and knockout organoids) was screened but not used in **Extended Data Fig. 5F** because it has one sample per condition.

Overlap of the upregulated signature with reference sets was tested by one-sided hypergeometric test within the 11,293 genes measured in at least seven contrasts (**Fig. 1K and Extended Data Fig. 5D). The three-way overlap in Fig. 1K of the upregulated signature (T, 578 genes), the 300 top-ranked regeneration genes (R, 168 measured) and the top 1,000 fetal genes (F, 820 measured) was compared with 200,000 Monte Carlo draws (NumPy default_rng, seed 0), each sampling three sets of the same sizes uniformly and independently without replacement from the same 11,293-gene universe; P was (number of draws with at least the observed triple overlap + 1)/(200,000 + 1). A conditional one-sided hypergeometric test in the same universe asked whether the T∩R genes were enriched for F genes (7 of 50; fold 1.9, P = 0.068; Supplementary Table 5**). The references were the 300 top-ranked genes of the regeneration meta-signature (168 measured), the 1,000 top genes by robust rank aggregation (516 measured), all 8,969 upregulated genes of the regeneration reference (5,557 measured), a curated revival stem-cell marker set (25 genes, 22 measured; Ayyaz *et al*, 2019; fetal_revival markers of the coupling analysis) and a curated YAP-target set (21 symbols for 19 genes, 15 measured; Cheung *et al*, 2020; Gregorieff *et al*, 2015; Yui *et al*, 2018; both lists in Supplementary Table 5) and fetal references (820 measured genes in the top fetal reference). These curated sets provide biological annotation of the signature. Their assembly from the same injury literature should be considered when interpreting enrichment.

Signature scores in YAP perturbation and microarray datasets (**Extended Data Fig. 5E-G**) were the mean z score of upregulated genes minus that of downregulated genes, with z scores computed within each dataset over log2 counts per million (single-cell counts of GSE152370 summed per sample) or RMA log2 intensities, and were recomputed with the rebuilt signature. These comprised YAP overexpression (GSE218333), LATS1/2 deletion (GSE152370), inducible YAP activation over 96 h in *Apc*-mutant organoids and YAP5SA in *Apc*/*Kras*/*Trp53*-mutant organoids (GSE152376; deposited Salmon transcript quantifications (Patro *et al*, 2017) summed to gene-level estimated counts), and Sca1+ repairing versus homeostatic colon epithelium (E-MTAB-5249; Yui *et al*, 2018). Groups were compared by Welch t test. GSE218333 and GSE152370 also contributed to the signature, so their scores are not independent validation; GSE152376 and E-MTAB-5249 did not contribute.

### Association of epithelial age with regenerative and proliferative programs

For **Fig. 1J and Extended Data Fig. 5C, 464** epithelial profiles were assembled from GSE165318, GSE123516, GSE222505, GSE152369, GSE152370, GSE230581, GSE218333, E-MTAB-14662, GSE262657 and GSE329635. Profile-level metadata are in Supplementary Table 5. GSE123516 contributed 42 irradiated/control epithelial cell-type pseudobulks, GSE152369 contributed 22 DSS/control epithelial pseudobulks, and GSE222505 contributed nine in-vivo Ki67-RFP-positive profiles at 56 h after irradiation or without irradiation. Single-cell data were summed by epithelial type or as pooled epithelium, while organoid and crypt data contributed whole samples. GSE192482 and GSE279059 were excluded because they could not be restricted to defined epithelial populations. Three curated scores were calculated as mean z scores of log2 counts per million within dataset. The fetal/revival set contained *Ly6a*, *Anxa1*, *Anxa3*, *Tacstd2*, *Clu*, *Il33*, *Msln*, *Cldn4*, *Areg*, *Emp1*, *Basp1*, *Spp1*, *Tnc*, *Ctgf*, *Cyr61*, *S100a6*, *Krt8*, *Krt19*, *Sprr1a*, *Sprr2a3*, *Anxa10* and *Gsdmc2*. The regeneration set contained *Ctgf*, *Cyr61*, *Ankrd1*, *Amotl2*, *Cd44*, *Sox9*, *Plaur*, *Serpine1*, *Tnc*, *Timp1*, *Lgals3*, *Ereg*, *Fn1*, *Vim*, *Anxa2*, *Sfn*, *Tagln*, *Actn1*, *F3*, *Lamc2*, *Sprr1a*, *Il1rn*, *Mmp7*, *Areg*, *Il33* and *Clu*. The proliferation set contained *Mki67*, *Top2a*, *Pcna*, *Ccnb1*, *Cdk1*, *Birc5*, *Ube2c*, *Mcm3*, *Mcm5*, *Cenpa* and *Ccna2*. *Ctgf*/*Ccn2* and *Cyr61*/*Ccn1 were matched by Entrez identifiers 14219 and 16007*. Clock outputs and scores were standardized within dataset-by-cell-type strata before Pearson correlation. Confidence bands used 2,000 bootstrap resamples of whole strata. Proliferation-adjusted associations correlated residuals after regressing both age and regeneration score on proliferation score. A mixed model with dataset-specific random intercepts and slopes was fitted by REML (statsmodels MixedLM). Clock-gene overlap was quantified for each set and is reported in Supplementary Table 5. Correlation P values are nominal because profiles remain nested within studies and animals.

### Chromatin accessibility

The deposited consensus-peak count matrix of bulk ATAC-seq from GSE222505 (18 libraries; Chen *et al*., 2023) was analyzed with pychromVAR, a Python implementation of chromVAR (Schep *et al*, 2017). Peak sequences were extracted from the mm9 genome (the coordinate system of the deposited peaks), 50 GC- and accessibility-matched background peaks per peak were selected by nearest-neighbor search (pychromVAR default; the implementation exposes no random seed), and JASPAR 2024 CORE vertebrate motifs (Rauluseviciute *et al*, 2024) were matched with MOODS at a motif P-value threshold of 5 × 10⁻⁵ with a pseudocount of 10⁻⁴ (pychromVAR defaults). Per-library motif deviation scores were averaged across motifs whose transcription-factor names matched each family (AP-1, JUN, FOS and BATF; TEAD; SMAD; NF-κB, NFKB and REL; STAT3; and HNF4 and CDX, counted as one intestinal-identity group, HNF4/CDX). Organoids derived after irradiation and treated with TGF-β1 for 7 days were compared with non-irradiated vehicle-treated organoids (three libraries each) by Welch t test without multiplicity adjustment. This compound contrast does not separate irradiation history from TGF-β1 exposure.

### Irradiation time course

The irradiation series comprised 22,680 cells from 17 male C57BL/6J mice before and 1, 3, 7 and 14 days after 15 Gy abdominal irradiation, with four, four, three, three and three mice, respectively (GSE165318, Lu *et al*, 2023). Marker scores across the authors’ 55 clusters assigned eight epithelial populations and immune cells. Stem cells were defined by *Lgr5*, *Olfm4*, *Ascl2*, *Smoc2*, *Slc12a2* and *Rgmb* scores without requiring *Lgr5* detection. A separate cell-level annotation requiring *Lgr5* detection was used in **Fig. 6A, a**s described below. Within each day, pooled epithelial cells were randomly split into 24 bins. Stem-cell and enterocyte bins contained approximately 30 cells, with at most 18 bins per day. Populations with fewer than 25 cells were omitted. Bins were scored with epithelial, SI tissue and pan-intestinal clocks and fetal-like and regeneration scores, using day 0 of the same population as reference. Means and confidence intervals were calculated over bins. Mann–Whitney day contrasts were BH-adjusted within panel, and age–regeneration relationships used Spearman correlation across bins. PCA used standardized log2 counts per million for the 2,000 most variable genes with mean expression of at least 10 counts per million. PERMANOVA quantified day effects on Euclidean distances, with 4,999 permutations and BH-adjusted day-versus-baseline tests. For pathways, counts were summed by day and population, retaining genes with at least 10 counts per million in either group and calculating log2 fold changes with pseudocount 1. Because mice were pooled before binning, these tests and intervals describe pooled cell distributions and do not estimate between-mouse variation.

### 5-FU time-course analyses

The 5-FU metacells and per-mouse pseudobulks were scored with the epithelial clock using YuGene or quantile-normalized inputs and with SI tissue and pan-intestinal clocks. Time-course plots referenced each age and population to its own day 0. Fetal-like and regeneration scores followed the irradiation procedure. Mann–Whitney tests compared each day with day 0, BH-adjusted within age, population and measure. Age groups were compared within day using Cohen’s d and Mann–Whitney tests, and age–regeneration associations used Spearman correlations across metacells. These tests do not account for dependence within mice. Per-mouse clock models used summed epithelial counts or cell-type profiles with at least 30 cells and 200,000 UMIs, referenced to young day-0 mice; these thresholds also removed old day-3 stem cells of one mouse and day-4 enterocytes of one old and one young mouse, leaving one mouse in those groups. Ordinary least-squares models adjusted for log10 total UMIs and estimated day-versus-baseline contrasts within age, age-by-day interactions for shared days, and common day effects, with BH adjustment across days. One low-yield old day-1 sample (D1_Pink) was excluded from these per-mouse clock models only: of its 721 called cells, 371 exceeded 50% mitochondrial counts and 205 passed all filters (its pre-filter summary gave medians of 217 genes and 55% mitochondrial counts), and its 187 epithelial cells carried 0.52 million UMIs, compared with at least 2.27 million in every other mouse. Models with this sample included are given in Supplementary Table 4. It was retained in the metacell analyses, PCA and PERMANOVA, the differential-expression and pathway contrasts (**Fig. 3E** and **Extended Data Figs. 10 and 14**) and the per-mouse display of **Extended Data Fig. 8**C. PCA and PERMANOVA used all 22 per-mouse profiles without a cell-number threshold; PERMANOVA tested day (ages pooled) and age (days pooled) as separate one-factor tests with 4,999 permutations, and day-versus-baseline contrasts were BH-adjusted across days. Mice were distinct at each collection day, and each day’s samples were multiplexed in one capture, so day and capture batch are confounded. Adjustment for depth (log10 of the summed UMIs of each profile) accounts for library size but cannot separate injury day from capture batch. Pooled-epithelium metacells contained 0.09-1.78 million UMIs (0.09-0.12 million for the low-yield old day-1 sample) and cell-type metacells 0.02-1.12 million UMIs (1st-99th percentile 0.08-0.67 million). Coverage benchmarks from Tyshkovskiy *et al* (2026) use reads, which are distinct from UMI counts. Cell-type metacell analyses support description of the sampled populations, with animal-level inference based on per-mouse models (Supplementary Table 4).

For pathway analysis, per-mouse pseudobulk counts of each cell type were compared between each sampled day and day 0 within age (two mice per group). Genes with at least 10 counts and 1 count per million in two or more samples were retained, log2 counts per million (prior count 2) were fitted with a two-group linear model, and gene-wise variances were moderated by empirical Bayes shrinkage as in limma (Smyth, 2004). The moderated t statistic ranked genes for enrichment. Pooled epithelium at day 4 was compared with day 0 within age using DESeq2 as implemented in PyDESeq2 (Muzellec *et al*, 2023), with genes ranked by the Wald statistic. Transit-amplifying and tuft-cell contrasts were not analyzed because of low cell numbers, and young day 3 was unavailable.

### Revival-marker-high cells and ligand-receptor analysis

GSE165318 counts were normalized to 10,000 per cell and log-transformed. Revival-marker-high epithelial cells exceeded the baseline 97.5th percentile of a six-gene score (*Ly6a*, *Clu*, *Anxa1*, *Msln*, *Spp1*, *Tacstd2*), yielding 467 cells. *Ly6a*-positive cells contained at least one *Ly6a* UMI (4,190 of 13,450 epithelial cells). The two analyses used different annotations of the same 22,680 GSE165318 cells. The original revival-marker-high analysis, run earlier in the project with an AI research agent, assigned the deposited clusters of Lu et al (2023) to 11 classes inferred from the publication, including stromal (1,342 cells) and endothelial (988 cells) classes; the later *Ly6a* analysis used a re-annotated object with eight epithelial classes (13,450 cells) and a single immune class for the remaining 9,230 cells, without stromal or endothelial labels. Differences in predicted partners between **Extended Data Fig. 4C and 4D** therefore reflect the annotation, not an absence of stromal or endothelial cells. In the 5-FU series, *Ly6a*-positive epithelial metacells were analyzed with per-sample cell-type pseudobulks as partners, with expression defined as at least 10 counts per million (28 young and 34 old metacells, approximately 35 cells each). LIANA 1.7.1 rank_aggregate used the mouse OmniPath consensus resource, minimum expression proportion 0.1, at least five cells per group, 1,000 permutations and seed 42 (Dimitrov *et al*, 2024, Dimitrov *et al*, 2022, Türei *et al*, 2021). Pairs with magnitude rank below 0.05 were summarized by sender and receiver. Ligand–target relationships were drawn from a curated literature list, not inferred with NicheNet. *Tgfb1* targets were *Smad7*, *Serpine1*, *Fn1*, *Ctgf*, *Col1a1*, *Cdkn1a*, *Jun* and *Snai1*. Targets were compared between marker-high and other epithelial cells by Mann–Whitney test, requiring BH FDR below 0.05 and more than 1.5-fold higher expression. In 5-FU, *Ly6a*-positive and negative metacells were compared by Mann–Whitney tests, followed by Wilcoxon signed-rank tests of within-sample contrasts across samples. The links in **Extended Data Fig. 4F** represent prior knowledge and expression associations among the sampled populations. The 5-FU libraries were prepared from EpCAM⁺CD31⁻CD45⁻ sorted cells. Partner pseudobulks labeled endothelial, smooth muscle, mast or immune (5,493 of 44,792 cells; 9% to 44% of the cells of a sample) expressed *Epcam* and *Krt8* at levels similar to epithelial pseudobulks and lacked canonical markers of these lineages (*Acta2*, *Myh11*, *Des*, *Tpsb2* and *Cma1* at or near zero; *Pecam1* and *Ptprc* low), so these labels mark epithelial cells assigned non-epithelial labels, residual contaminating cells, doublets or sorting leakage rather than niche populations (Supplementary Table 5). Interactions involving these labels are not interpreted as stromal, vascular or immune signals, and the 5-FU preparations do not provide a census of the tissue niche. These six-gene and *Ly6a* definitions differ from the 22-gene fetal/revival panel, the 300-gene fetal-like score, the GSE115541 fetal reference, the 28-gene oncofetal panel of the spatial analyses and the Mustata fetal-spheroid score used elsewhere.

### Comparison of regenerative programs

Gene-level log2 fold changes for six regeneration contrasts (GSE222505 bulk TGF-β1 organoids, GSE152370, GSE279059, GSE218333, GSE262657 and E-MTAB-14662) and the fetal-versus-adult contrast (GSE230581) were taken from the consensus-signature analysis; the GSE222505 re-plated-organoid 24-h contrast is not used (no time-matched vehicle). For the **Fig. 6A** contrasts (*Lgr5*⁺ stem cells and enterocytes at day 3 versus day 0), each GSE165318 cell was assigned individually to the highest of seven standardized marker-gene scores (stem, *Lgr5*, *Olfm4*, *Ascl2*, *Smoc2*, *Slc12a2* and *Rgmb*; enterocyte, *Alpi*, *Apoa1*, *Apoa4*, *Fabp1*, *Fabp2*, *Apob*, *Rbp2* and *Ckb*; and transit-amplifying, goblet, Paneth, tuft and enteroendocrine sets), requiring a score of at least 0.3 and a margin of 0.15 over the next score. Stem cells with detectable *Lgr5* were classed as *Lgr5*⁺ stem cells, yielding 128 and 45 *Lgr5*+ stem cells and 1,144 and 172 enterocytes at days 0 and 3. Counts were summed per cell type and day, and log2 fold changes were computed from counts per million (pseudocount 1) for genes with at least 10 counts per million in either sample. For 5-FU, day 4 was compared with day 0 within each age in per-mouse pooled-epithelium profiles (limma-trend on log-transformed RLE-normalized counts, two mice per group). As for **Fig. 1I, S**pearman correlations were computed over the union of the 1,000 genes with the largest absolute log2 fold change in each of the two contrasts (most contrasts have no replicate-based P value). BH-adjusted Spearman P values are displayed across contrast pairs. Because genes are co-regulated, these are nominal measures based on gene-level units and are not tests across independent biological samples.

### Gene-set enrichment and pathway-level correlation

Pre-ranked gene-set enrichment analysis (Subramanian *et al*, 2005) used MSigDB mouse Hallmark and Reactome collections (v2024.1.Mm), KEGG legacy sets (v2024.1) and the GO biological-process sets present in the reference matrix described below, restricted to sets of 15 to 500 genes. The KEGG legacy sets are defined with human genes and were mapped to mouse genes through one-to-one human–mouse orthologs (the ortholog table of the tAge pipeline; Tyshkovskiy *et al*, 2026), which retained a median of 94% of the genes of each set. Enrichment was computed with a custom Python implementation of pre-ranked GSEA (fastgsea2.py), not with fgsea. Enrichment scores used the weighted (p = 1) running-sum statistic, with the absolute ranking statistic as weight. For each gene-set size, 20,000 random gene sets of identical size formed the null distribution (gene-set permutation; in each draw, the sets of every size were the leading genes of one random ordering of the ranked genes; seed 42). The normalized enrichment score (NES) was the enrichment score divided by the mean of the null scores of the same sign, the nominal P value was (number of same-sign null scores at least as extreme + 1)/(number of same-sign null scores + 1), and the false discovery rate was controlled by the Benjamini-Hochberg method within each contrast and collection (Hallmark, KEGG, Reactome and GO). The implementation was not benchmarked against a reference GSEA implementation in this study. For **Extended Data Fig. 4A,B, H**allmark sets and 20 KEGG metabolic pathways (10 to 500 genes) were analyzed in the same way. The rows of **Extended Data Fig. 15A** were chosen by a fixed rule, following Tyshkovskiy *et al* (2026) in requiring significance in at least one profile: among Reactome and KEGG pathways evaluated in the published reference matrix, those with FDR < 0.05 in at least one of the ten post-surgical contrasts were ranked within each direction of their mean NES by the smallest FDR (ties broken by the largest absolute NES), a pathway was skipped when more than half of the genes of the smaller set were shared with a higher-ranked pathway, and the first 12 pathways of each direction are shown.

GSEA rankings were DESeq2 Wald statistics for liver and pooled 5-FU epithelium, moderated t statistics for 5-FU cell types and replicated clock-independent contrasts, log2 fold changes for unreplicated regeneration contrasts and GSE165318, signed −log10 meta-analysis P values for intestinal aging, and signed aggregation or differential-expression statistics for regeneration and fetal references. Gene-wise meta-analyses required measurements in at least two studies. Published reference NES came from two generations of signatures. Correlation matrices in **Figs. 2E, 3E, 6B and 7E and Extended Data Figs. 10 and 15B** used the 2,943-pathway matrix of tissue-aging and lifespan-intervention signatures from Tyshkovskiy et al (2019, 2023), excluding its species-lifespan, brain and kidney signatures. The Hallmark heatmap in **Fig. 6C** used the 2026 rodent chronological-age, mortality, maximum-lifespan and species-aging signatures (Tyshkovskiy *et al*, 2026). The selected Reactome and KEGG pathways in **Fig. 6D** used the aging and lifespan-intervention references from the earlier matrix (Tyshkovskiy *et al*, 2019, 2023). The two generations cover different pathway libraries and were not mixed in one correlation matrix. Similarity was Spearman correlation across jointly evaluated pathways (1,519-2,943). Displayed significance in **Figs. 2E, 3E, 6B and 7E and Extended Data Figs. 10D and 15B** uses BH adjustment across the pairs specified in the legends. These P values treat overlapping pathways as independent and are nominal. **Extended Data Fig. 10A-C** reports coefficients without tests.

### Spatial transcriptomics

Visium data were analyzed for irradiated SI from mice on standard diet, without irradiation (0 dpi) and 3 days after 10 Gy irradiation (GSE227742; Das *et al*, 2025), *Il10*-deficient colon at weeks 10 and 14 and at week 14 with rectal prolapse with a wild-type week-6 reference (GSE256234; Fransson *et al*, 2026), and AOM/DSS colon (GSE227598, below). Each condition was represented by a single Visium section (one capture area). Spots with fewer than 500 counts and genes detected in fewer than 10 spots were excluded. Each spot was assigned the cell type with the highest marker-gene score (the score_genes function in Scanpy) among curated SI or colon epithelial, stromal, immune and muscle marker sets, and this dominant label was used for cell-type comparisons. Because Visium spots contain several cells, the label reflects the dominant expression program rather than a deconvolved cell-type proportion.

Spatial prediction adapted the reference-relative approach of stAge (Vicente-Alvarez *et al*, 2025). The epithelial clock was applied to individual spots and to aggregates restricted to epithelial regions. Bulk, pseudobulk and epithelial-metaspot applications computed YuGene over the 9,670-gene training panel. For per-spot estimates, computing YuGene over the training panel or over all genes of the spot gave nearly identical values (Pearson r = 1.00; median difference 0.02 months), and recomputation from the deposited GSE227742 matrices agreed with the reported spot estimates (r = 0.985; c66_spot_panel_check.py). All applications used log2(1 + 10⁴ × YuGene), training means and standard deviations, and centering on the relevant reference section. Missing clock genes were set to zero, and predictions were multiplied by 48. Per-spot estimates were sensitive to coverage. Median counts per in-tissue spot were 6,256 and 8,288 in the GSE227742 standard-diet sections (0 and 3 dpi) and 271 to 882 in the GSE256234 colon sections (735 to 1,346 among epithelial-compartment spots; c67_spot_counts.py). In GSE227742, spot depth and epithelial ΔtAge were correlated (Spearman ρ approximately −0.26 to −0.32). For **Fig. 4E,F and Extended Data Fig. 11, e**pithelial spots assigned to crypt-base, transit-amplifying, differentiated or revival compartments were pooled by k-means clustering of full-resolution coordinates, with k equal to total counts/300,000 rounded to an integer and seed 0. Raw counts were summed and metaspots with fewer than 200,000 counts were excluded; this cutoff follows the metacell depth used by Tyshkovskiy et al (2026) and is an analysis choice, not a validated threshold. Regeneration and fetal-like scores were calculated per metaspot and z-scored across metaspots; metaspot scores used the 300 top-ranked genes of the epithelial regeneration meta-signature and the 300 top fetal-higher genes of the GSE115541 reference. Per-spot scores (**Fig. 4A,B**, **Fig. 5** and **Extended Data Fig. 12**) and Visium HD scores (**Fig. 4G**, **Extended Data Fig. 13**) instead used Scanpy score_genes on normalized, log-transformed counts, with a 28-gene curated oncofetal epithelial panel for the fetal-like score and the 200 top-ranked genes of a core intestinal regeneration meta-signature for the regeneration score; the core signature was rebuilt by robust rank aggregation from 13 regeneration-versus-control contrasts with retained per-dataset rankings (build_regen_metasignatures.py). Both gene lists and the 13-contrast roster are in Supplementary Table 5. GSE227742 used GSM7106693/GSM7106694, and GSE256234 used GSM8091664-GSM8091667. Estimates were referenced to the 0-dpi SI section or wild-type week-6 colon section. For *Il10*-deficient colon, chronological time since week 6 was subtracted from spot and metaspot estimates (0.93 months at week 10 and 1.87 months at week 14). Program hotspots exceeded the reference-section 95th percentile. Section contrasts were summarized by mean differences, with depth-adjusted estimates from linear regression on log sequencing depth. Regeneration-score deciles were defined within sections and compared with reference spots, and age–regeneration associations used Spearman correlations. Cell-type comparisons used the same dominant label in injured and reference sections. Plotted spot and metaspot comparisons used Mann–Whitney tests, and hotspot proportions used Fisher’s exact tests, with BH adjustment in the families specified in the legends. As each condition comprised one section, these tests are nominal, as plotted: they describe differences among subsamples of those sections and do not estimate between-animal variation. Maps used seven-nearest-neighbor smoothing for display only.

Visium HD jejunum data comprised male C57BL/6J mice before and 1-6 days after 12 Gy whole-body irradiation on four capture areas (GSE303705, Ofer *et al*, 2026). GEO lists 23 mice in the deposited experiment. Using the authors’ bin-level day and mouse labels (GitHub tht38/intestinalspatialregen; Zenodo 21348331), 19 mice carried an identifier, the labeled day-2 piece of capture area 2 without an identifier was treated as one further mouse and the unlabeled day-4 piece of that capture area was not analyzed, giving 20 mice in the present analysis (day 0, n = 5; days 1, 2, 3 and 5, n = 3; day 4, n = 2; day 6, n = 1), of which 17 contributed metaspots with at least 200,000 counts. Each capture area included baseline tissue. Space Ranger 8-µm bins were assigned day and mouse using deposited metadata (GitHub tht38/intestinalspatialregen, Zenodo 21348331). Nineteen mice had identifiers. A labeled day-2 piece without an identifier was treated as a further mouse, and an unlabeled day-4 piece was excluded, yielding 20 mice (day 0, n = 5, days 1, 2, 3 and 5, n = 3, day 4, n = 2, day 6, n = 1). Bins required at least 10 counts and agreement between marker-based epithelial identity and the authors’ epithelial-region annotation. Within each mouse, epithelial bins were pooled into 320-µm squares, retaining aggregates with at least 200,000 counts. This yielded 897 metaspots from 17 mice, as the shallow first capture area contributed no qualifying baseline or day-1 aggregate. Epithelial and SI tissue clocks used their respective training transformations, with YuGene calculated over the epithelial training panel or all detected genes, respectively, and all day-0 metaspots as reference. Program scores were z-scored across metaspots. For display, 24-µm aggregates with at least 1,000 counts were referenced to baseline in the same capture area and smoothed across seven neighbors. Inference used per-mouse means fitted as y ∼ day + capture area, with BH adjustment across day contrasts within readout. The single day-6 mouse was not tested. Per-mouse estimates were also compared with estimates from all epithelial bins pooled within mouse.

### Colitis-associated tumorigenesis

AOM/DSS colon Visium sections at days 0, 22 and 43 on standard diet (GSE227598, Das *et al*, 2025) were jointly filtered and scored relative to day 0. The expression-based dysplasia score contrasted *Axin2*, *Lgr5*, *Ascl2*, *Notum*, *Nkd1*, *Sox9*, *Cd44*, *Mki67*, *Top2a*, *Ly6a*, *Tacstd2*, *Cldn2*, *Reg3b*, *Reg3g* and *Olfm4* with differentiated epithelial genes *Car1*, *Car4*, *Ceacam1*, *Slc26a3*, *Aqp8*, *Guca2a*, *Guca2b*, *Muc2*, *Fabp2*, *Krt20* and *Slc26a2*. It was not a histological classification. Map hotspots exceeded the day-0 95th percentile. **Fig. 5F** compared dysplasia, fetal-like and regeneration hotspots (2,530, 3,016 and 4,851 of 11,470 spots) with all other spots using log2 ratios of mean counts per 10,000 plus one. For **Fig. 5E,G, u**pper and lower score tertiles among epithelial-type spots defined high-score and reference regions. **Fig. 5G** compared high-score spots of each day with the same dominant cell type in low-score regions pooled across days 22 and 43. Mann–Whitney tests with BH adjustment were calculated across spots. Hotspot proportions, score-decile comparisons and age–regeneration correlations followed the spatial methods above. **Fig. 5D** used Cohen’s d and intervals calculated as described for regenerative comparisons, with Welch t tests across spots. All sections had one sample per condition, so spot-level tests and intervals do not estimate between-animal variation.

Single-cell profiles of AOM/DSS lesions and normal colon and of *Apc*-mutant tdTomato+ tumors (GSE224679; Bala, 2023) were annotated by marker scoring and summarized in cell-type pseudobulk bins within each tissue. Cell types within a tissue were compared with enterocytes by one-way ANOVA with Dunnett’s test, and lesion and normal cells of the same type by Welch t test, on random bins of pooled cells from one sample per group; these tests compare bins within single samples and do not test differences between samples (**Extended Data Fig. 12C,D**).

### Liver regeneration

GSE185705 comprised 36 young and old C57BL/6JN livers before and 48, 72, 96, 120 and 240 h after two-thirds partial hepatectomy, with three livers per age and timepoint (4 and 18 months, both sexes, Yang *et al*, 2023). Raw featureCounts were processed with the tAge pipeline (Tyshkovskiy *et al*, 2026). Genes required at least 10 counts in 20% of samples and were mapped to Entrez identifiers, RLE-normalized, log-transformed, scaled or YuGene-transformed, and centered on young pre-surgical liver. Published coefficients were applied for 24 multi-species, rodent multi-tissue, mouse multi-tissue and rodent liver clocks covering chronological age, normalized age and mortality, with scaled or YuGene input, and for rodent module clocks. Chronological outputs were multiplied by 48 and normalized-age outputs by 100. The **Fig. 7B** output retains the intercept and constant contribution of absent features, so expression centering does not make the young baseline prediction zero; it is therefore shown as a reference-centered clock output rather than as ΔtAge from young pre-surgical liver. Within-age post-surgical-minus-baseline differences do not require recalibration. Post-surgical effects were differences from pre-surgical predictions within the same age. Planned contrasts used the pooled residual variance of six groups per age (12 degrees of freedom), with BH adjustment over ten post-surgical contrasts per clock. Module adjustment covered 23 modules and the all-module estimate across ten contrasts (240 tests per panel). Welch tests with the same correction families provided a sensitivity analysis, reported in Supplementary Table 7. Baseline old-versus-young comparisons used Welch tests. Age-dependent responses were assessed as the difference between age groups in the mean of five post-surgical groups minus baseline, using pooled variance (24 degrees of freedom) and BH adjustment over 24 clocks. Module effects were divided by the standard deviation across all livers. DESeq2 compared timepoints within age and baseline old versus young liver. The within-cohort aging contrast was shown in **Fig. 7F** but not correlated with regeneration responses because they share reference animals. Shared-reference effects were evaluated in 500 null simulations using observed gene-wise residual variances, limma and the union of the top 1,000 genes. Gene-level comparisons instead used independent ITP liver chronological-age and mouse multi-tissue aging signatures (Tyshkovskiy *et al*, 2026). Liver, hepatocyte and old-liver regeneration meta-signatures used robust rank aggregation (Kolde *et al*, 2012) from a candidate collection comprising partial hepatectomy (GSE282210, GSE243302, GSE263879), carbon tetrachloride (GSE176042, GSE180366), acetaminophen (GSE277619, GSE291946, GSE252192) and organoids (GSE308085). Individual ranks, contrast definitions and exact contributor rosters were not retained and could not be recovered; the aggregated lists (robust rank aggregation scores, adjusted P values, ranks and the signed −log10 score weights used for GSEA), the candidate roster and the fetal-liver identity signature are given in Supplementary Table 8. GSE185705 was absent from that collection. The fetal-liver identity signature compared E14.5-E16.5 liver with other fetal tissues in ENCODE (ENCSR867YNV, ENCSR611PTP, ENCSR826HIQ), using DESeq2 one-versus-all contrasts, adjusted P < 0.01, log2 fold change > 1 and protein-coding genes absent from lung, kidney, forebrain and limb signatures. Its 2,888 genes were ranked by Wald statistic. This signature measures fetal liver identity relative to other fetal organs, not fetal-versus-adult liver development.

### Statistics and reproducibility

Tests, units and multiple-testing families are specified in the figure legends. Tests were two-sided unless stated otherwise, with BH adjustment within the stated families. Symbols denote * P or q < 0.05, ** < 0.01, *** < 0.001 and **** < 10⁻⁴, with ns for non-significance. Gene-set enrichment heatmaps additionally use ^ for FDR < 0.1. Effect sizes are mean differences or Cohen’s d with the intervals described above. Intervals calculated over bins or spots quantify variation among those subsamples. Cells, metacells, bins and spots from the same animal, pooled sample or section are not independent biological replicates. Their plotted tests are descriptive and do not estimate between-animal variation. Gene- and pathway-correlation tests likewise use non-independent features, so their adjusted P values are nominal. Animal-level inference for the 5-FU series used per-mouse models (Supplementary Table 4), with two mice per age and sampled day before the stated exclusions. No day-2 or young day-3 samples were analyzed. Irradiation GSE165318 cells were pooled across mice before binning. Visium HD inference used mouse-level means with capture area as a covariate. No statistical method predetermined sample size.

### Software

The 5-FU pipeline used Python 3.12, Scanpy 1.11.4, anndata, harmonypy, Scrublet through Scanpy, scikit-learn 1.7.1, decoupleR 2.1.6 and LIANA 1.7.1. Other expression, clock and statistical analyses used Python 3.13.16, NumPy 2.5.3, pandas 2.3.3, SciPy 1.18.1, statsmodels 0.15.0, scikit-learn 1.7.1, PyDESeq2 0.5.4 and InMoose 0.9.1. Chromatin analysis used pychromVAR 0.0.4, MOODS and Biopython. Empirical-Bayes linear models (Smyth, 2004), restricted-maximum-likelihood meta-analysis following metafor rma.uni, and pre-ranked GSEA were implemented in the supplied Python modules revlib.py and fastgsea2.py. Microarray normalization and earlier signature construction used R 4.3.3 with Bioconductor 3.18. Figures used matplotlib 3.11.2 and CairoSVG 2.9.1.

Dataset retrieval, intestinal and liver regeneration rank-aggregation signatures, the initial LIANA analysis and curated ligand–target comparison, and **Extended Data Fig. 6** activity estimates were generated with an AI research agent (Edison Scientific) under author direction. The authors computed aging signatures, clock training and applications, time-course and spatial analyses, pathway correlations and figure statistics using the accompanying code. The *Ly6a* analysis in **Extended Data Fig. 4D-F** was recomputed with LIANA 1.7.1, Scanpy 1.10.4 and anndata 0.11.4.

## Data availability

Single-cell RNA-sequencing data from the 5-FU time course will be deposited in GEO before journal submission, with a reviewer token, and the accession will be given here. All other datasets are publicly available under the accessions given in the Methods and figure legends. The Visium HD irradiation data are available from GEO (GSE303705), and their bin-level day and mouse labels from GitHub (tht38/intestinalspatialregen) and Zenodo (record 21348331). Source data (the per-sample, per-bin and per-spot values behind every quantitative panel) accompany this version.

## Code availability

Code for clock training, validation and application, model coefficients, reference expression profiles and gene sets will be made available upon publication.

## Author contributions

S.I.: conceptualization, funding acquisition, experiments, data analysis, writing of the original draft, and review and editing. V.N.G.: conceptualization, discussion, supervision, funding acquisition, and manuscript review and editing.

