## Supplemental Figures for "Transcriptomic rejuvenation during intestinal and liver regeneration in mice"

Extended Data figures with legends. Each figure is on its own page and is followed by its legend on the next page.

**Extended Data Fig. 1 | Pooled and study-level validation of the intestinal clocks and age ranking in additional cohorts**

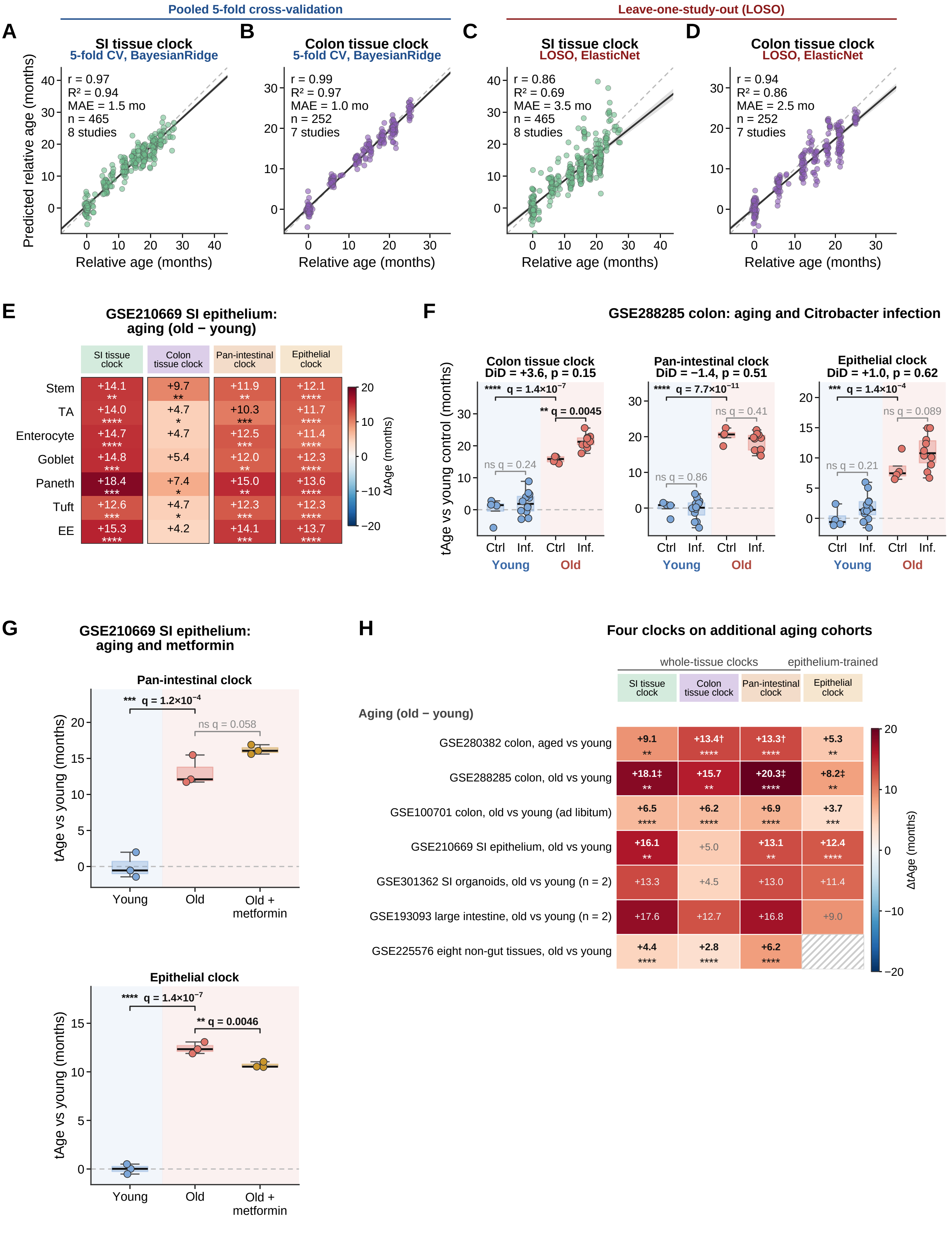

### Extended Data Fig. 1 | Validation of intestinal clocks and age ordering in additional cohorts.

**(A,B)** SI tissue and colon tissue clocks under pooled five-fold cross-validation with BayesianRidge (BR), using 465 samples from eight studies and 252 samples from seven studies, respectively. **(C,D)** The same samples under leave-one-study-out (LOSO) validation with ElasticNet (EN). A and B differ from Fig. 1A,B in validation scheme, while C and D differ in final estimator. Ages are relative to the youngest group of each study. Points represent samples, the solid line and band show a least-squares fit and 95% bootstrap confidence interval, and the dashed line indicates identity. **(E)** Old-minus-young ΔtAge by epithelial cell type in GSE210669. Counts were summed per mouse and cell type, requiring at least 15 cells, with three mice per age. Stars show BH-adjusted planned contrasts across cell types within clock. Residual variance was pooled across young, old, rapamycin-treated old and metformin-treated old groups. Exact permutation P values range from 0.1 to 0.3, with a minimum attainable P of 0.1. **(F)** Mid-colon before and after Citrobacter rodentium infection (GSE288285), on colon tissue, pan-intestinal and epithelial clocks. Young and old control groups contain four samples each. Infected groups contain 12 young and 10 old samples pooled across days 3-10. Estimates are relative to young controls. DiD denotes the age-by-infection difference-in-differences. **(G)** Young, old and metformin-treated old SI epithelium (GSE210669), with three mice per group. F and G use pooled-variance planned contrasts and BH-adjusted P values. Permutation results are provided in Source Data. Boxes show medians and interquartile ranges, whiskers extend to 1.5 times the interquartile range, and points represent samples. **(H)** Old-minus-young ΔtAge in seven additional cohorts. The GSE288285 and GSE210669 rows repeat the control comparisons in F and G. GSE225576 contains eight non-intestinal tissues from four male mice per tissue and age, comparing 24 and 30 months with 6 months using tissue as a fixed effect. Deposited metadata do not establish whether tissues came from the same mice. All 27 evaluated contrasts predict older states. Eighteen have $q < 0.05$, including 13 of the 22 contrasts independent of the indicated clock's training data. † marks a training cohort and ‡ another cohort from the same training publication. Hatched cells were not evaluated. Welch P values were BH-adjusted within clock, with stars shown only for groups of at least three samples. Gray values indicate $q \geq 0.05$. Stars indicate P or $q < 0.05$ (*), < 0.01 (**), < 0.001 (***) or $< 10^{-4}$ (****). ns, not significant.

**Extended Data Fig. 2 | Individual observations and temporal profiles underlying the regenerative comparisons**

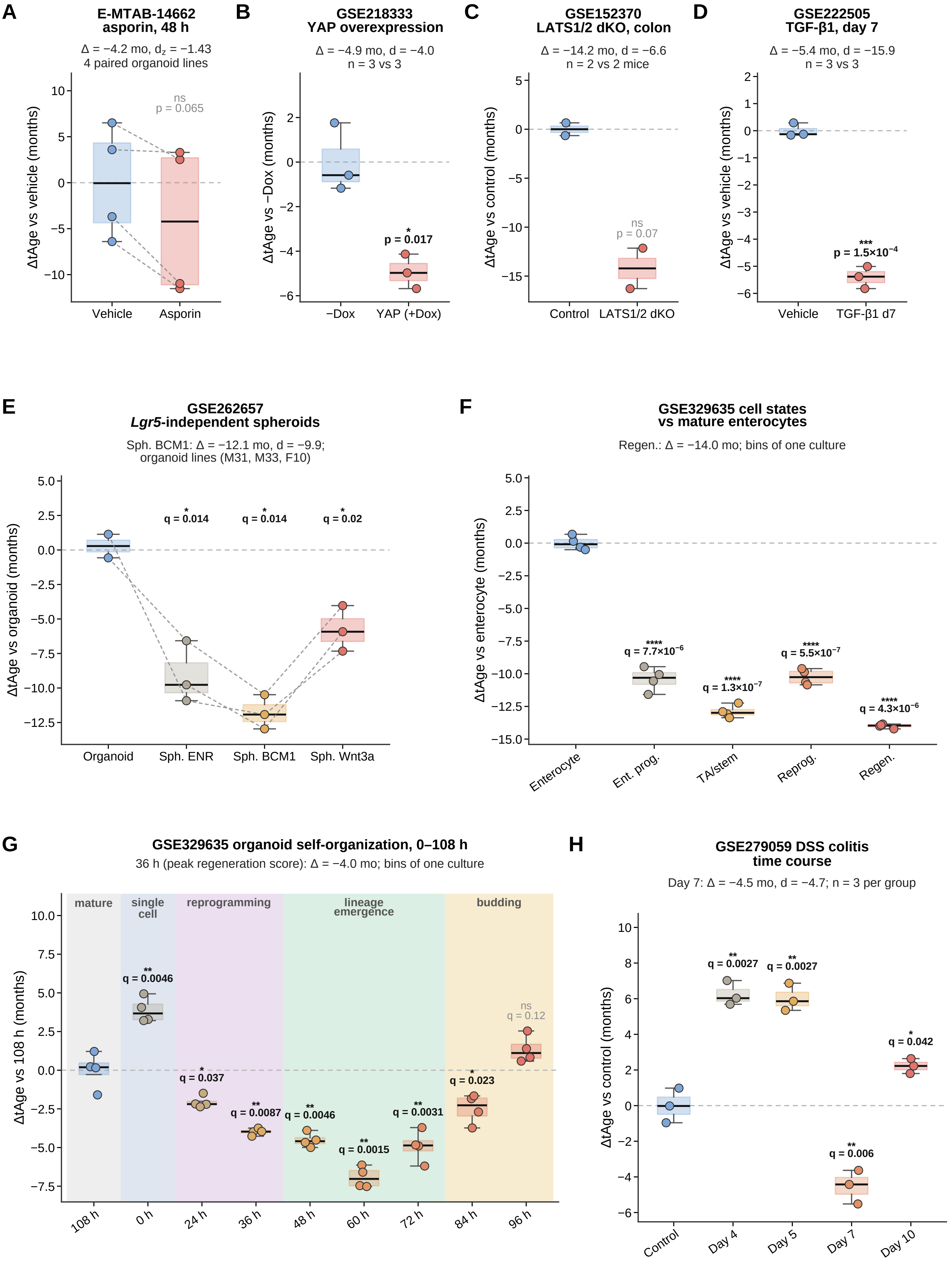

### Extended Data Fig. 2 | Individual observations and time courses underlying regenerative comparisons.

Epithelial-clock estimates are relative to the indicated reference group. **(A)** Recombinant asporin versus vehicle after 48 h in organoids from four donor mice (E-MTAB-14662). Lines connect donor-matched pairs, tested by paired t test. **(B)** YAP overexpression versus uninduced organoids (GSE218333, three samples per group). **(C)** LATS1/2-deficient versus control colon epithelium (GSE152370, two mice per group). **(D)** TGF-β1-treated versus vehicle organoids at day 7 (GSE222505, three samples per group). B-D use Welch tests. **(E)** *Lgr5*-independent spheroids in three media compared with organoids from lines M31, M33 and F10 (GSE262657). Gray lines connect matching lines. F10 has no organoid sample, so each Welch comparison uses three spheroid and two organoid line means, with BH adjustment across the three media. **(F)** Cell states versus mature enterocytes during organoid self-organization (GSE329635). Ent. prog., enterocyte progenitors. TA/stem, transit-amplifying and stem-cell progenitors. Reprog., reprogramming. Regen., regenerative. Labels follow the deposited data (Schwayer et al, 2026). Differences between these states include cell-identity effects. **(G)** Organoid self-organization, referenced to the mature 108-h condition. Shading marks phases defined by Schwayer et al, 2026. The header gives the 36-h contrast selected for Fig. 1H by its regeneration score. F and G use random bins from one culture per time point, with Welch tests and BH adjustment within panel. These tests describe bins from the sampled cultures and do not provide replication between cultures. **(H)** DSS colitis time course in bulk colonic crypts (GSE279059), with three mice per time point, referenced to control. Mice received 3% DSS for five days followed by water. Welch comparisons were BH-adjusted across sampled days. Boxes show medians and interquartile ranges, whiskers extend to 1.5 times the interquartile range, and points represent samples or bins. Δ, mean difference. d and d_z, independent-sample and paired Cohen's effect sizes. *Star* thresholds are as in Extended Data Fig. 1.

### Extended Data Fig. 3 | Age and expression-program responses across regenerative conditions

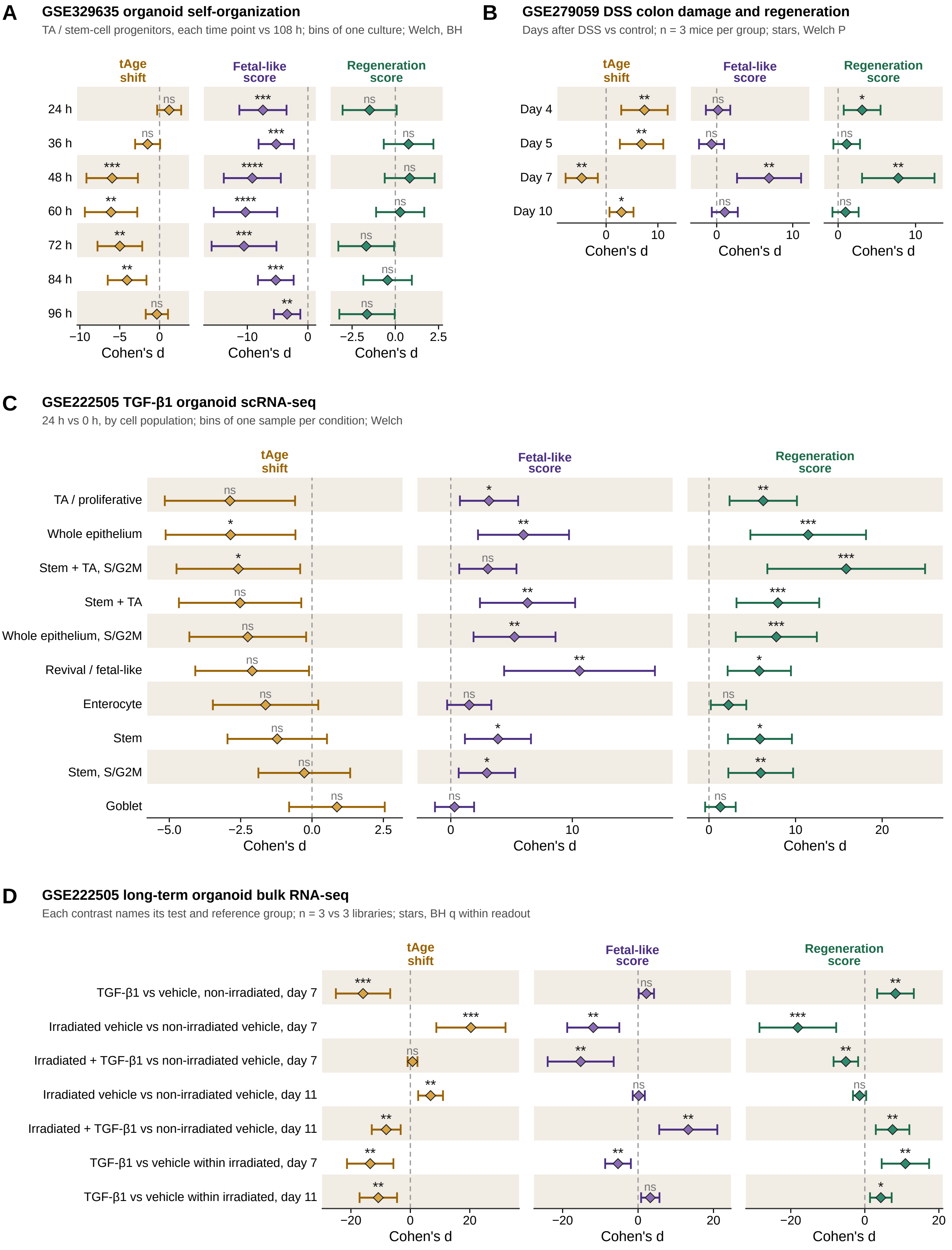

### Extended Data Fig. 3 | Epithelial age and expression programs across regenerative conditions.

Diamonds show Cohen's d and 95% confidence intervals for epithelial-clock age, fetal-like score (300 top fetal-higher genes of the GSE115541 reference; Methods) and regeneration score on the same units. The epithelial clock uses YuGene input and BayesianRidge. **(A)** Transit-amplifying and stem-cell progenitors during organoid self-organization, compared with 108 h (GSE329635). **(B)** DSS colonic crypts versus control, with three mice per group (GSE279059). **(C)** Organoids 24 h after re-plating with TGF-β1, compared with 0 h (GSE222505). Whole-epithelium, stem/progenitor and S/G2M groups overlap and are not independent populations. This comparison has no time-matched vehicle control. A and C use random bins from one sample per time point. Welch tests therefore compare bins within the sampled cultures, with BH adjustment within panel in A. B and C show unadjusted Welch P values. **(D)** Long-term organoid cultures (GSE222505), with three libraries per group. Each row names its treatment and reference, either non-irradiated vehicle organoids at day 7 or vehicle organoids of the same irradiation history and day. Welch tests were BH-adjusted across seven contrasts within each readout. The day-7 TGF-β1-versus-vehicle comparison repeats Extended Data Fig. 2D. *Star* thresholds are as in Extended Data Fig. 1.

**Extended Data Fig. 4 | Expression pathways and candidate niche signals associated with regenerative states**

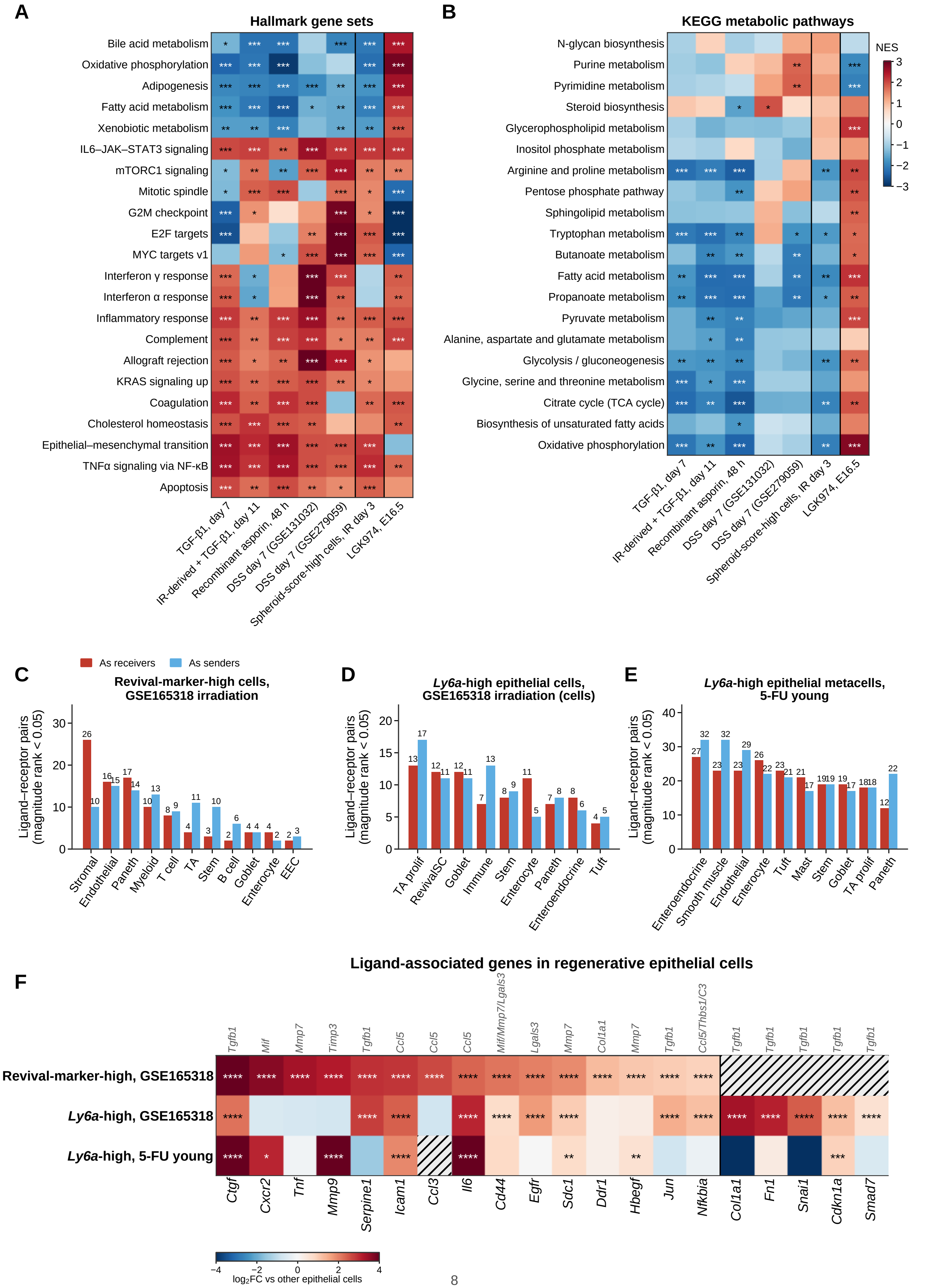

### Extended Data Fig. 4 | Expression pathways and candidate niche signals associated with regenerative states.

**(A)** Hallmark normalized enrichment scores (NES) for the seven contrasts in Fig. 1I, showing the 22 sets with the largest mean absolute NES. Contrasts include TGF-β1-treated organoids, irradiation-derived organoids treated with TGF-β1 (GSE222505), asporin-treated organoids (E-MTAB-14662), DSS colon (GSE131032 and GSE279059), spheroid-score-high cells at irradiation day 3 (GSE165318) and fetal epithelium treated with LGK974 (GSE183532). Genes were ranked by moderated t statistics, paired for asporin. The pooled irradiation contrast used $\log_2$ fold changes. GSEA used 20,000 random gene sets, with BH adjustment within contrast and collection. Stars in the irradiation column assess the ranking within a pooled profile, not reproducibility between animals. **(B)** Twenty KEGG metabolic pathways analyzed as in A. NES describes expression programs, not metabolic flux. **(C)** LIANA-predicted ligand-receptor pairs with magnitude rank < 0.05 involving 467 revival-marker-high cells among 22,680 cells in GSE165318. Red indicates these cells as receivers and blue as senders. **(D)** The corresponding analysis for 4,190 *Ly6a*-positive cells among 13,450 epithelial cells, using a later annotation of the same cells with no stromal or endothelial label (all 9,230 non-epithelial cells form one immune class), so differences in partners from C reflect the annotation (Methods). Positivity requires at least one *Ly6a* UMI. The annotation used in D includes epithelial and immune cells but no stromal or endothelial categories. **(E)** Young 5-FU samples analyzed using 28 *Ly6a*-positive epithelial metacells, comprising 1,136 cells, and per-sample pseudobulks of partner cell types. The libraries were EpCAM CD31 CD45 sorted; partner labels named for non-epithelial lineages express epithelial markers and do not represent a census of stromal, vascular or immune cells (Supplementary Table 5). Expression required at least 10 counts per million. **(F)** Expression of genes associated with the candidate ligands indicated above each column, compared with other epithelial cells. Ligand-target assignments are curated from prior knowledge. Color shows $\log_2$ fold change. Stars mark increases above 1.5-fold with BH-adjusted Mann-Whitney P values on cells or, for 5-FU, metacells. Hatched cells were not tested or were unavailable in the original analysis. Additional curated TGF-β1 targets appear to the right of the divider. The 5-FU differences were not consistent within mice. For example, *Ctgf* was higher in *Ly6a*-positive metacells in one of ten mouse-day samples. C-F describe expression-based signaling candidates in pooled cells or metacells and do not establish ligand action or animal-level replication. *Star* thresholds are as in Extended Data Fig. 1.

**Extended Data Fig. 5 | Consensus younger-state signature: recurrent genes, adjustment for a proliferation score, overlap with regeneration programs and response to YAP activity**

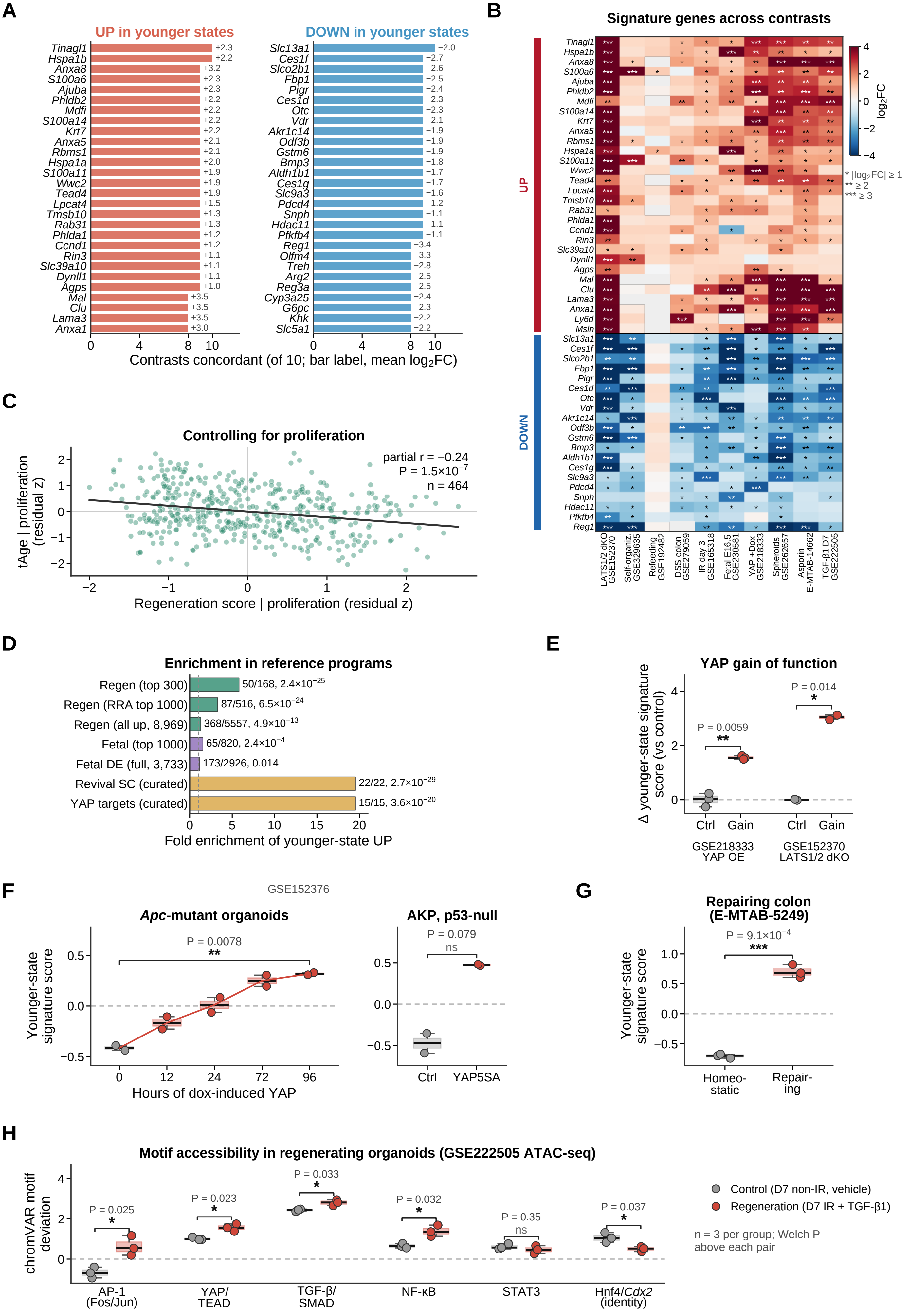

### Extended Data Fig. 5 | Genes and regulatory programs associated with younger epithelial states.

**(A)** The leading 28 upregulated and 28 downregulated younger-state genes, ranked by recurrence across ten contrasts. Bar labels show mean $\log_2$ fold change. The full signature contains 578 upregulated and 550 downregulated genes, requiring detection in at least seven contrasts and a change of at least 1.5-fold in the same direction in at least 60% of eligible contrasts. **(B)** $\text{Log}_2$ fold changes for 30 upregulated and 20 downregulated genes across the ten contrasts. Asterisks indicate absolute $\log_2$ fold changes of at least 1, 2 or 3, not statistical significance. Time-course contrasts use the lowest clock estimate, including organoid self-organization at 60 h versus 108 h, DSS day 7, irradiation day 3 and refeeding versus ad libitum feeding. Fig. 1H instead selects time points by regeneration score. **(C)** Association of epithelial age with regeneration score after removing their linear relationships with the proliferation score, across 464 profiles. Partial r = -0.24, nominal P = $1.5 \times 10^{\ 7}$. **(D)** Enrichment of younger-state upregulated genes in regeneration, fetal (GSE115541 reference; Methods), revival stem-cell and YAP-target sets, tested by one-sided hypergeometric tests. Labels give overlap, measured reference-set size and P. The curated revival set contains 25 genes, of which 22 were measured; it is distinct from the 22-gene composite fetal/revival panel of Fig. 1J (gene lists in Supplementary Table 5). The YAP set contains 21 symbols representing 19 genes, of which 15 were measured. Fig. 1K shows the three-way overlap. **(E)** Younger-state signature scores after YAP overexpression (GSE218333, three samples per group) and LATS1/2 deletion (GSE152370, two mice per group). These datasets contributed to signature discovery. **(F)** Signature scores during inducible YAP activation in *Apc*-mutant organoids, with two samples per time point and a 96-h-versus-0-h Welch comparison, and after YAP5SA expression in *Apc*/*Kras*/*Trp53*-mutant organoids, with two samples per group (GSE152376; the two deposited replicate samples of each condition are the units, and GEO does not state whether they are independent organoid lines or cultures of one line). This dataset did not contribute to signature discovery. **(G)** Sca1 repairing versus homeostatic colon epithelium (E-MTAB-5249), measured by microarray with three samples per group. E-G use Welch tests. **(H)** chromVAR motif-accessibility deviations in irradiation-derived organoids treated with TGF-β1 versus non-irradiated vehicle organoids (GSE222505 ATAC-seq, three libraries per group). Unadjusted Welch P values are shown for six transcription-factor families. The contrast combines irradiation history and TGF-β1 exposure. Boxes in E-H show medians and interquartile ranges, with individual samples plotted. *Star* thresholds are as in Extended Data Fig. 1.

**Extended Data Fig. 6 | Inferred transcription-factor and pathway activities across the signature-derivation contrasts**

**A**

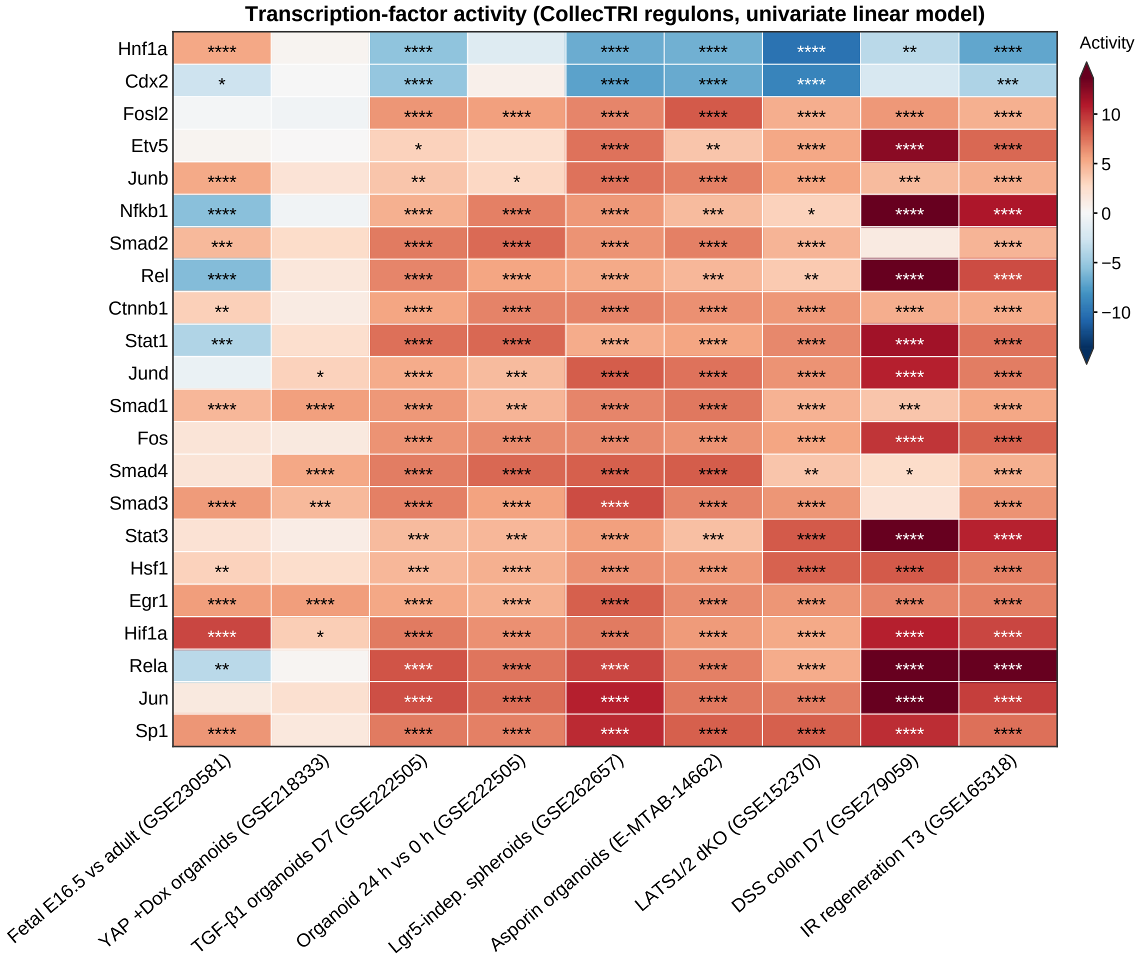

**B**

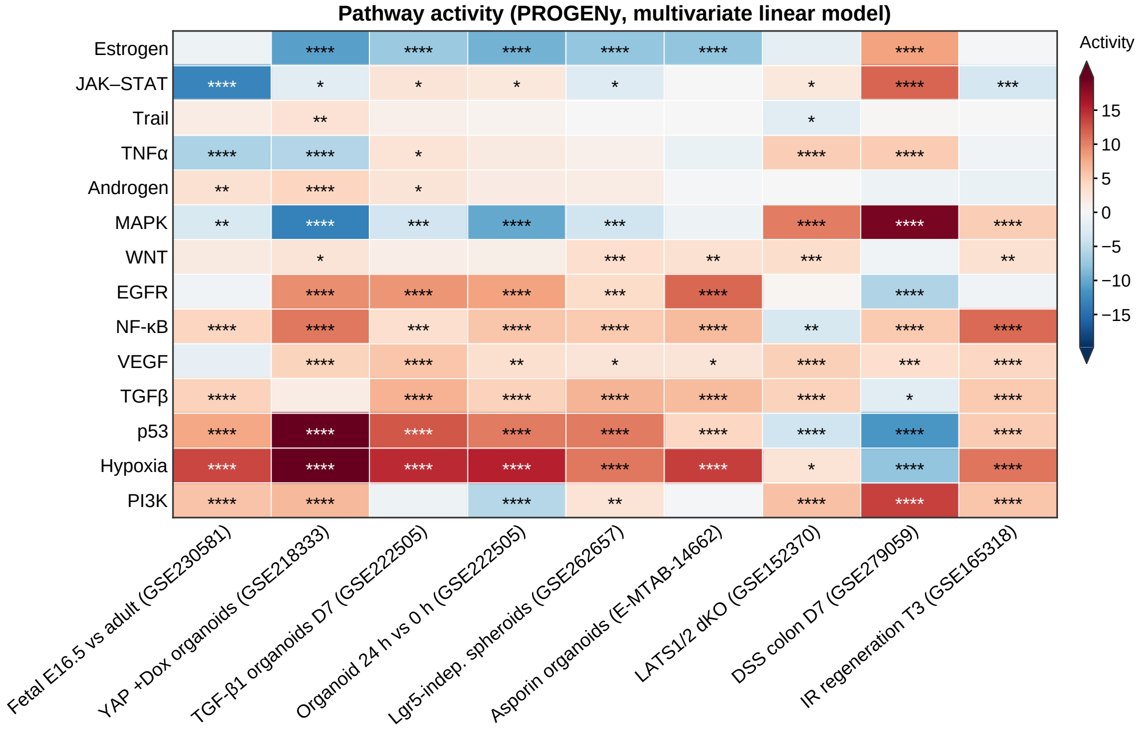

### Extended Data Fig. 6 | Inferred transcription-factor and pathway activities in regenerative and developmental contrasts.

**(A)** CollecTRI transcription-factor activity estimated from $log_2$ fold changes with the decoupleR 2.1.6 univariate linear model. **(B)** PROGENy pathway activity estimated with the multivariate linear model. Red indicates higher inferred activity in the regenerative or fetal state. Stars indicate unadjusted per-contrast model P values. Color scales saturate at ±13.6 in A and ±19.8 in B. The nine displayed contrasts include the signature-derivation comparisons other than self-organization (GSE329635) and refeeding (GSE192482), plus the 24-h re-plating comparison (GSE222505) shown for context. The latter does not contribute to the younger-state signature. IR regeneration T3 denotes pooled epithelium at day 3 versus baseline in GSE165318. Values are provided in Source Data. *Star* thresholds are as in Extended Data Fig. 1.

**Extended Data Fig. 7 | Stem-cell revival markers and sensitivity of the 5-FU epithelial trajectory to normalization and clock**

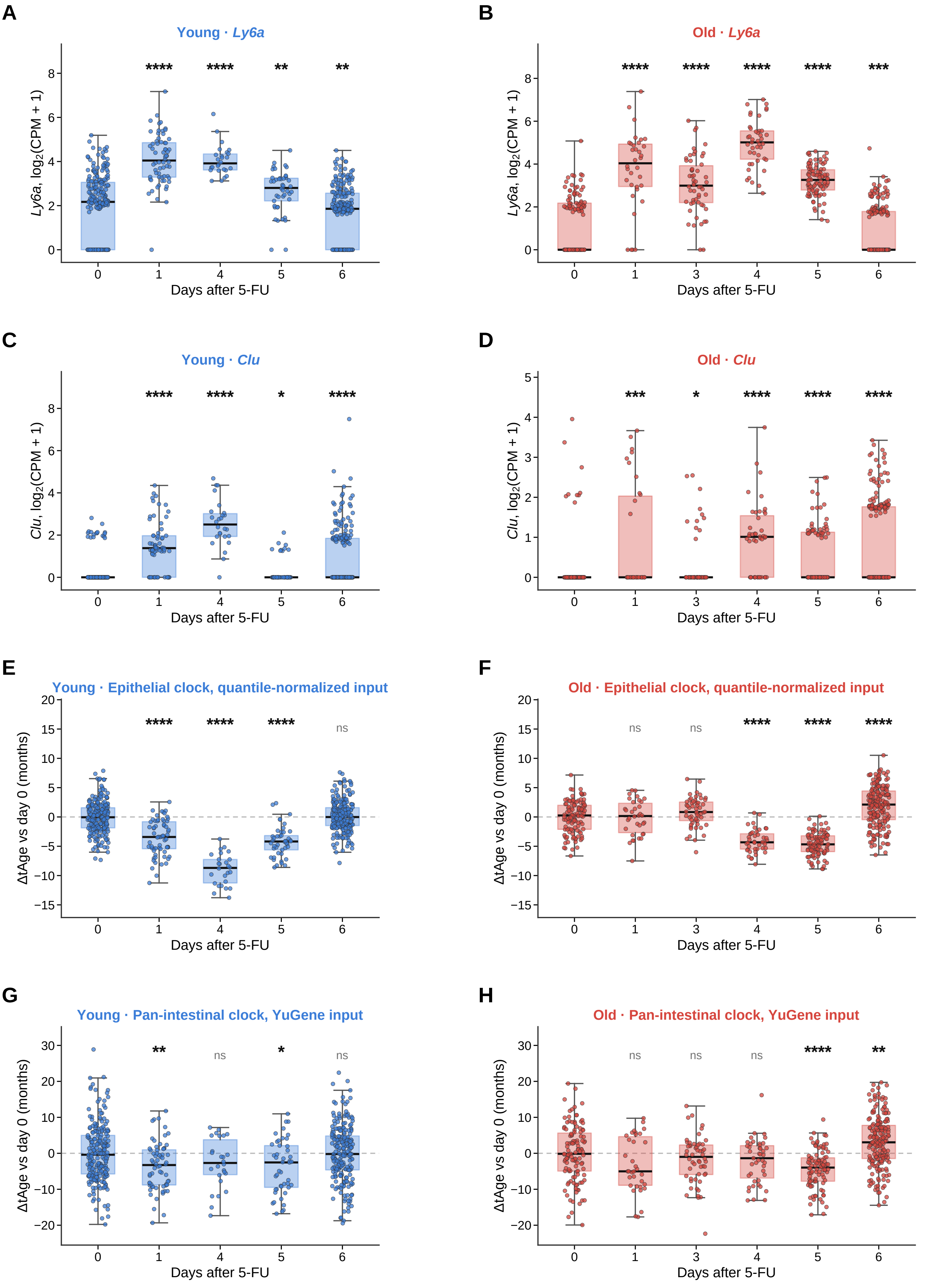

### Extended Data Fig. 7 | Revival-associated genes and sensitivity of the 5-FU age response to normalization and clock.

Metacells of approximately 35 cells from young and old epithelium, with each day compared with day 0 of the same age. **(A,B)** *Ly6a* expression in young and old epithelium. **(C,D)** *Clu* expression in young and old epithelium. Gene expression is shown as $\log_2$(counts per million + 1). **(E,F)** Epithelial-clock ΔtAge after quantile normalization of input profiles. **(G,H)** Pan-intestinal-clock ΔtAge with YuGene input, using the validated model of Fig. 1C. Boxes show medians and interquartile ranges, whiskers extend to 1.5 times the interquartile range, and points represent metacells. Stars denote Mann-Whitney comparisons with day 0, BH-adjusted within panel. Metacells derive from two mice per age and day, so tests describe metacell differences and do not provide animal-level inference. Young day 3 yielded insufficient cells. Per-mouse estimates from the primary YuGene epithelial clock appear in Extended Data Fig. 8C and Supplementary Table 4. *Star* thresholds are as in Extended Data Fig. 1.

### Extended Data Fig. 8 | Cell-type and animal-level age trajectories after 5-FU injury

In-house single-cell RNA-seq, two young and two old mice per day (young day 3 not analyzed). A, B, D: metacells; C: per-mouse pseudobulk.
Stars, Mann–Whitney test versus day 0 across metacells of two mice per age and day, BH-adjusted.

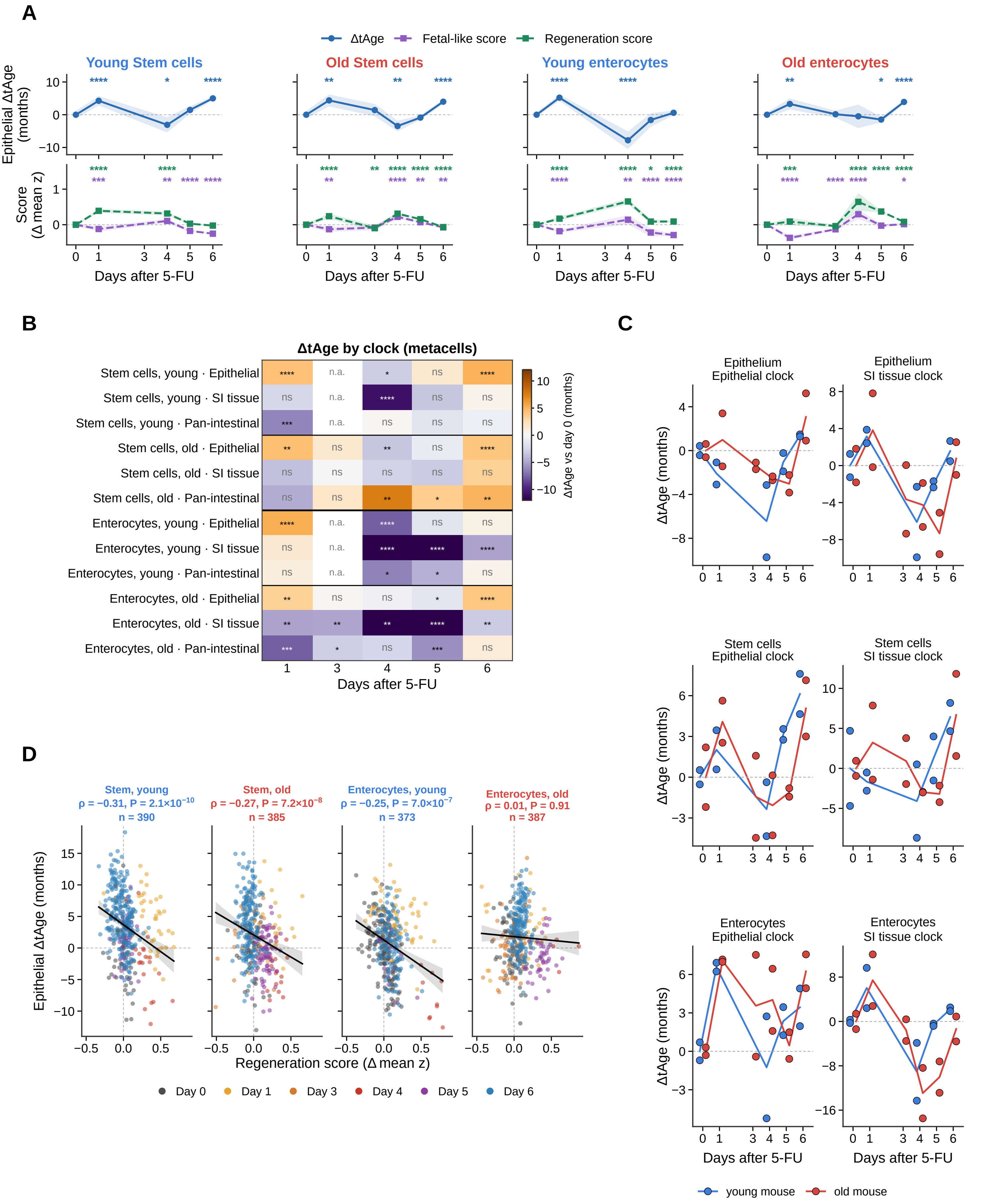

Points, individual mice (including the low-yield day-1 old sample); lines, daily means

### Extended Data Fig. 8 | Cell-type and animal-level age responses after 5-FU injury.

**(A)** Epithelial ΔtAge and fetal-like and regeneration scores in stem cells and enterocytes, relative to day 0 of the same age and population. Lines and bands show metacell means and 95% confidence intervals, with a common y range within each row. **(B)** Mean ΔtAge on epithelial, SI tissue and pan-intestinal clocks for the same populations. Stars in A and B show Mann-Whitney comparisons with day 0, BH-adjusted within panel. n.a., not analyzed. **(C)** Per-mouse pseudobulk estimates for pooled epithelium, stem cells and enterocytes on epithelial and SI tissue clocks, referenced to day 0 of the same age. Points represent mice and lines show daily means, with young mice in blue and old mice in red. C shows all 22 mice as unadjusted differences from day 0 of the same age, including the low-yield old day-1 sample and cell-type profiles below the 30-cell and 200,000-UMI thresholds; the animal-level clock models exclude these profiles (Supplementary Table 4). No tests are shown in C. **(D)** Epithelial ΔtAge versus regeneration score across stem-cell and enterocyte metacells, separated by age. Lines and bands show least-squares fits and 95% confidence intervals. Spearman ρ, nominal P and metacell counts are given. Day colors follow Fig. 3. A, B and D analyze metacells from the sampled mice and do not provide animal-level inference. Animal-level models are reported in Supplementary Table 4. *Star* thresholds are as in Extended Data Fig. 1.

### Extended Data Fig. 9 | Epithelial age after 5-FU on a clock trained without the developmental-reversion and regeneration programs and on the original clock capped at its training range

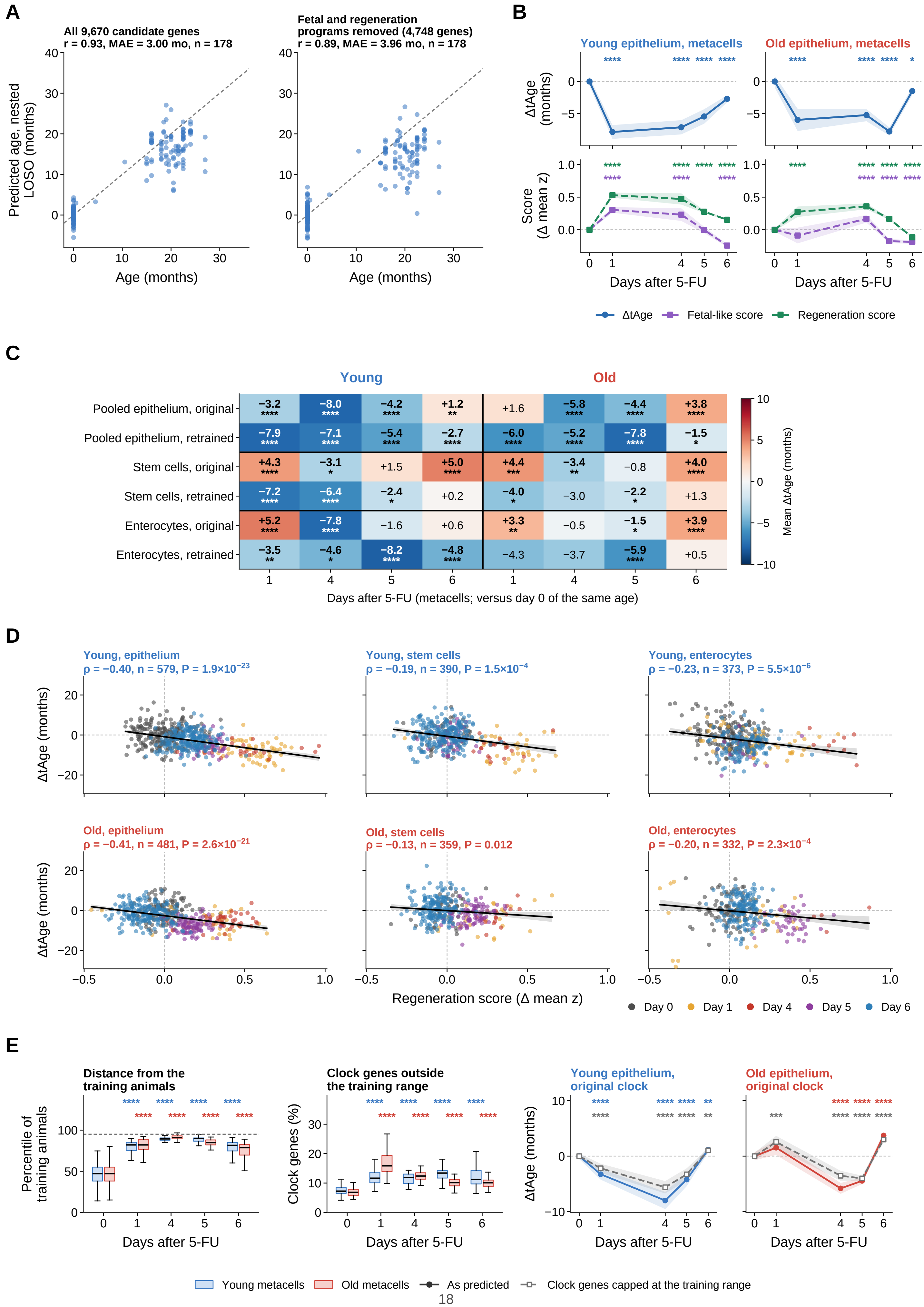

### Extended Data Fig. 9 | Age predictions after excluding developmental and regenerative genes or restricting inputs to the training range.

The epithelial clock was retrained after removing all genes differentially expressed between E16.5 and adult SI epithelium, the leading epithelial regeneration genes, the curated repair and fetal/revival sets and the younger-state signature. This removed 4,922 of 9,670 candidate genes, leaving 4,748, before variable-gene selection. Training and nested leave-one-study-out validation followed Fig. 1. **(A)** Predictions for 178 training animals from models retrained on all candidates or the reduced gene set. Pearson r and mean absolute error are shown. The dashed line indicates identity. **(B)** Reduced-gene-clock ΔtAge and fetal-like and regeneration scores across pooled epithelial metacells, relative to day 0 of the same age. Lines and bands show means and 95% confidence intervals. **(C)** Mean ΔtAge on the original and reduced-gene clocks for pooled epithelium, stem cells and enterocytes. Each population is referenced to day 0 of the same age. **(D)** Reduced-gene-clock ΔtAge versus regeneration score in the three populations. Lines and bands show least-squares fits and 95% confidence intervals. Spearman ρ, nominal P and metacell counts are given. **(E)** Original-clock inputs compared with those of 178 training animals. Left, Mahalanobis distance in the first 20 training principal components, expressed as a percentile of training-animal distances. The dashed line marks the 95th percentile. Middle, percentage of 2,000 clock genes outside the training range. Boxes show medians and interquartile ranges, with whiskers extending to 1.5 times the interquartile range. Right, ΔtAge before and after capping each clock-gene input at its training minimum and maximum, shown as means and 95% confidence intervals. B, C and E use Mann-Whitney comparisons with day 0, BH-adjusted within age, population and readout as applicable. Gray stars in E refer to capped predictions. Metacells derive from the sampled mice, so these tests and correlations are descriptive of metacells, not animal-level replication. Old day 3 is omitted because of low cell yield. Per-mouse models are provided in Supplementary Table 6. *Star* thresholds are as in Fig. 1.

**Extended Data Fig. 10 | Pathway-level correlation of the two regeneration time courses with aging, regeneration and fetal references**

Spearman ρ between pathway normalized enrichment scores (NES) of each injury contrast (versus day 0 of the same age) and of the reference signatures; no P values are shown because overlapping pathways are not independent. Boxed, the injury contrasts. A: GSE165318; B–D: in-house 5-FU single-cell RNA-seq.

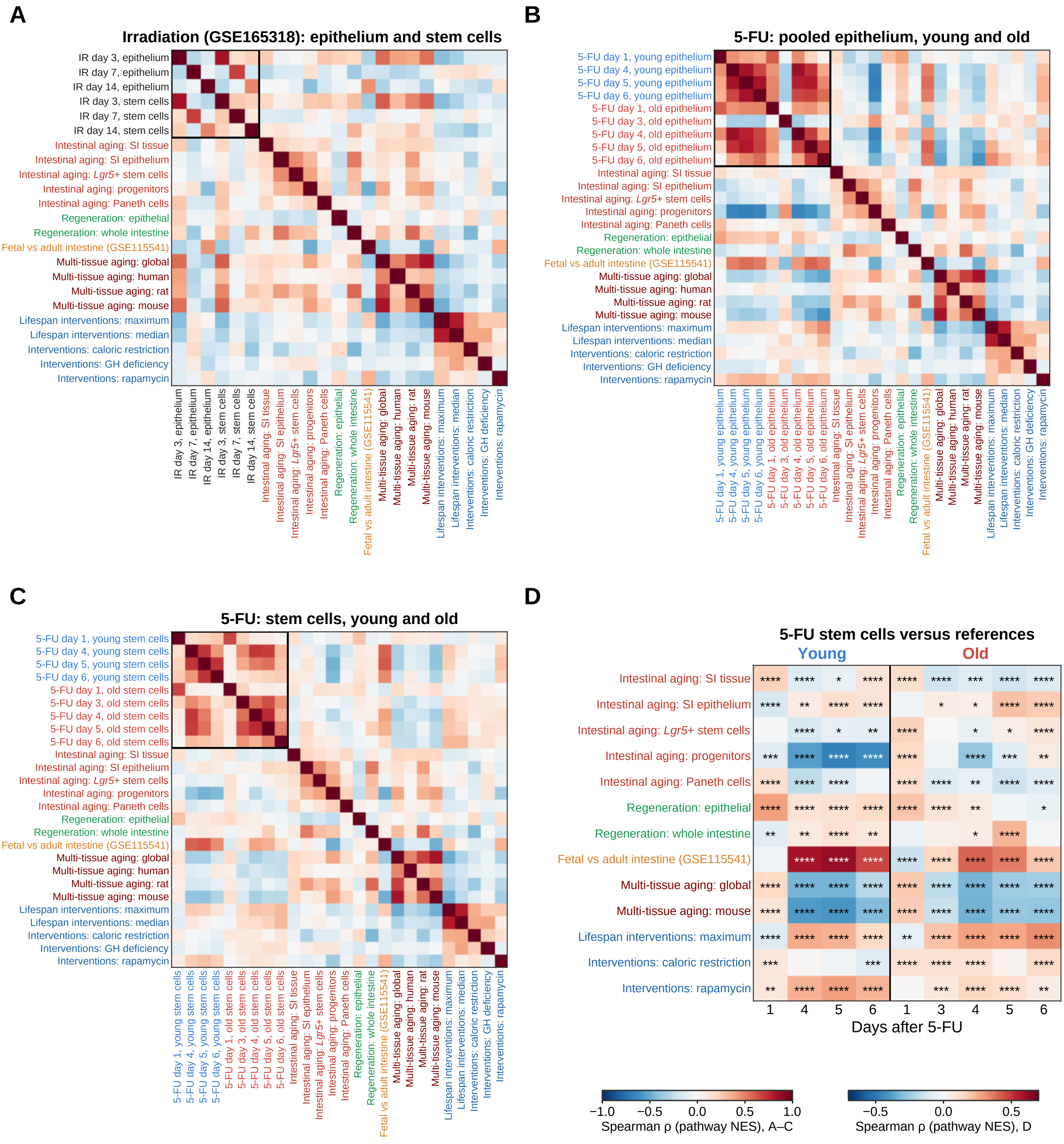

### Extended Data Fig. 10 | Pathway relationships between regeneration, aging and fetal expression.

Spearman correlations of pathway normalized enrichment scores (NES) for injury responses and intestinal aging, regeneration, fetal, multi-tissue aging and lifespan-intervention references. Boxes identify injury-response contrasts. **(A)** Pooled epithelium and stem cells at irradiation days 3, 7 and 14 versus day 0 (GSE165318). Genes were ranked by $\log_2$ fold changes in pooled day pseudobulks. **(B)** Young and old pooled epithelium at each sampled day after 5-FU versus baseline. This is the full matrix underlying Fig. 3E. **(C)** Corresponding 5-FU stem-cell responses. B and C use per-mouse pseudobulk differential-expression contrasts. **(D)** Stem-cell responses against selected reference signatures. A-C show correlations without P values. In D, stars are BH q values across all 468 contrast-reference pairs of the two time courses, the family used in Fig. 2E; the q values saved for the A–C matrices (BH within each matrix) are not displayed. These calculations treat overlapping pathways as independent and should be read descriptively. The reference signatures and gene-ranking methods are specified in Methods. *Star* thresholds are as in Extended Data Fig. 1.

**Extended Data Fig. 11 | Spatial epithelial age after irradiation at metaspot resolution**

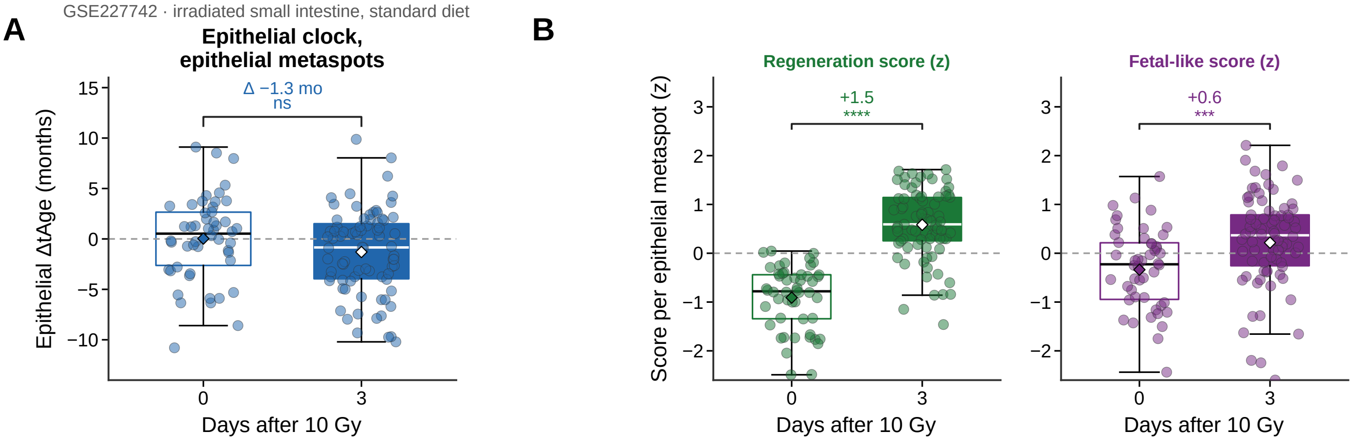

### Extended Data Fig. 11 | Spatial epithelial age after irradiation at metaspot resolution.

**(A, B)** Irradiated small intestine (GSE227742) before and 3 days after 10 Gy irradiation (dpi) on standard diet. **(A)** Epithelial ΔtAge of epithelial metaspots (spots of the crypt-base, transit-amplifying, differentiated and revival compartments pooled by k-means on their coordinates; at least 200,000 counts; 50 to 78 metaspots per section), relative to the 0-dpi section. **(B)** Regeneration and fetal-like scores of the epithelial metaspots in A (z-scored across metaspots; fetal-like score from the GSE115541 reference, Methods). Boxes, median and interquartile range; whiskers, 1.5 × interquartile range; dots, metaspots; diamonds, means; Δ, mean difference from the reference section; stars, Mann–Whitney test across metaspots (* $P < 0.05$, ** < 0.01, *** < 0.001, **** < 0.0001; ns, not significant). Each condition is represented by one section, so the Mann–Whitney tests compare metaspots within single sections and do not test differences between sections or animals.

**Extended Data Fig. 12 | Epithelial age predictions in colitis and tumorigenesis by disease stage and cell type**

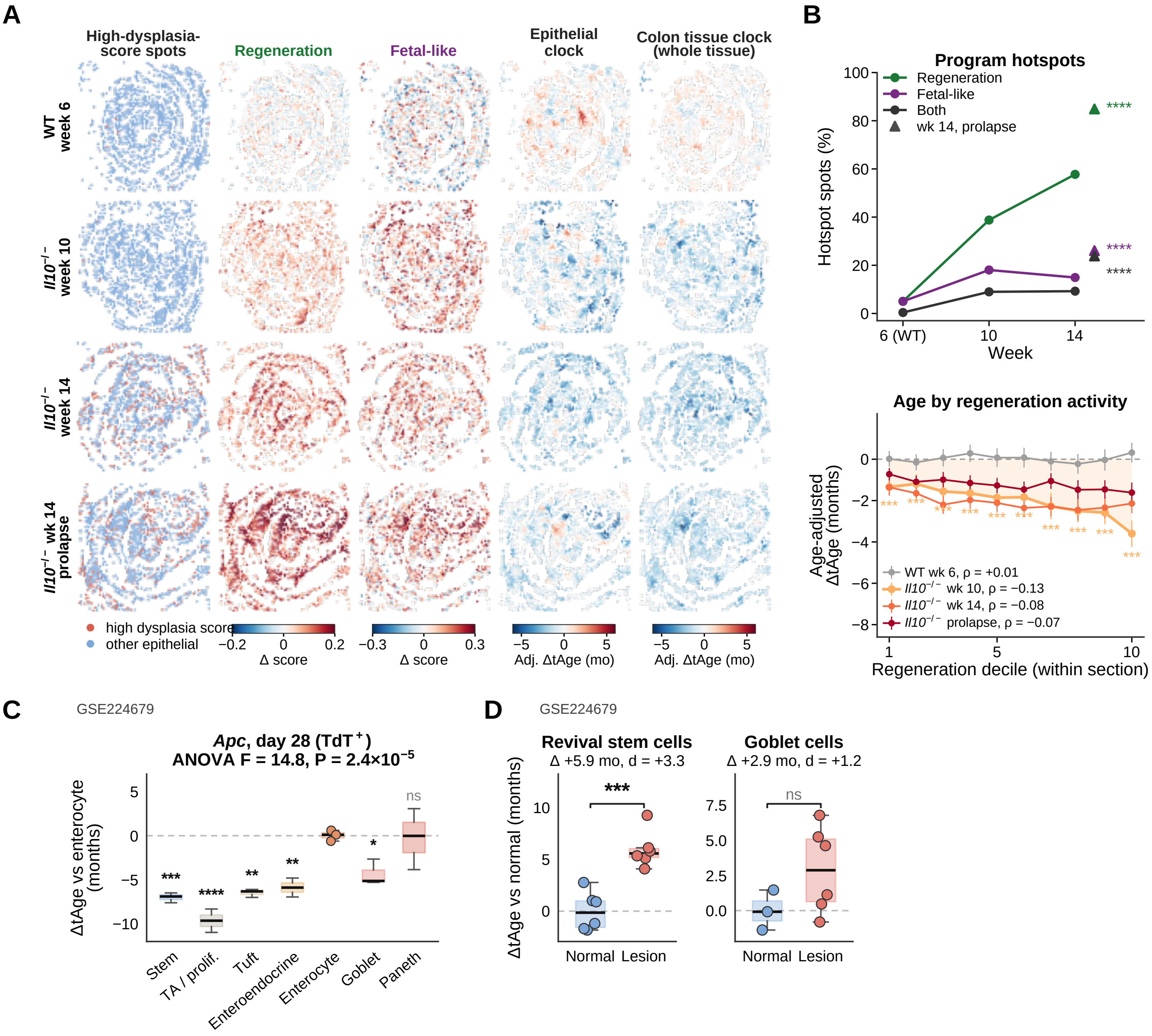

### Extended Data Fig. 12 | Epithelial age predictions in colitis and tumorigenesis by disease stage and cell type.

**(A)** *Il10*-deficient colon (GSE256234), showing maps of high-dysplasia-score spots, regeneration and fetal-like scores, and epithelial and colon tissue clock estimates relative to the wild-type week-6 section (scores minus the week-6 mean; age estimates minus the chronological age elapsed since week 6, 0.93 months at week 10 and 1.87 months at week 14, and smoothed over seven nearest spots for display). **(B)** Quantification of A. Top, proportions of spots above the 95th percentile of the week-6 section. Bottom, age-adjusted epithelial ΔtAge (as in A) by within-section regeneration-score decile (legend, Spearman ρ between regeneration score and ΔtAge across the spots of each section). **(C)** Cell-type pseudobulk estimates within *Apc*-mutant (tdTomato ) tumors from GSE224679, relative to enterocytes of the same tissue (random bins of pooled cells; one-way ANOVA with Dunnett comparison against enterocytes; bins of one sample per group, so not a test between samples). **(D)** AOM/DSS lesion-versus-normal comparisons within revival stem and goblet cells (GSE224679; random bins of pooled cells; Welch t test; bins of one sample per group, so not a test between samples). In B, stars for the hotspot proportions compare the week-14 prolapse section with week 6 (Fisher's exact test, BH; every week-10 and week-14 proportion also exceeds week 6, $q < 10^{25}$), and decile stars compare week-10 deciles with week-6 spots (Mann–Whitney, BH). Panels B–D use spots or bins from one section or sample per group, so their tests compare spots or bins within single samples and do not test differences between animals. Stars as in Extended Data Fig. 1.

**Extended Data Fig. 13 | Spatial epithelial age across the irradiation time course in replicated Visium HD jejunum**

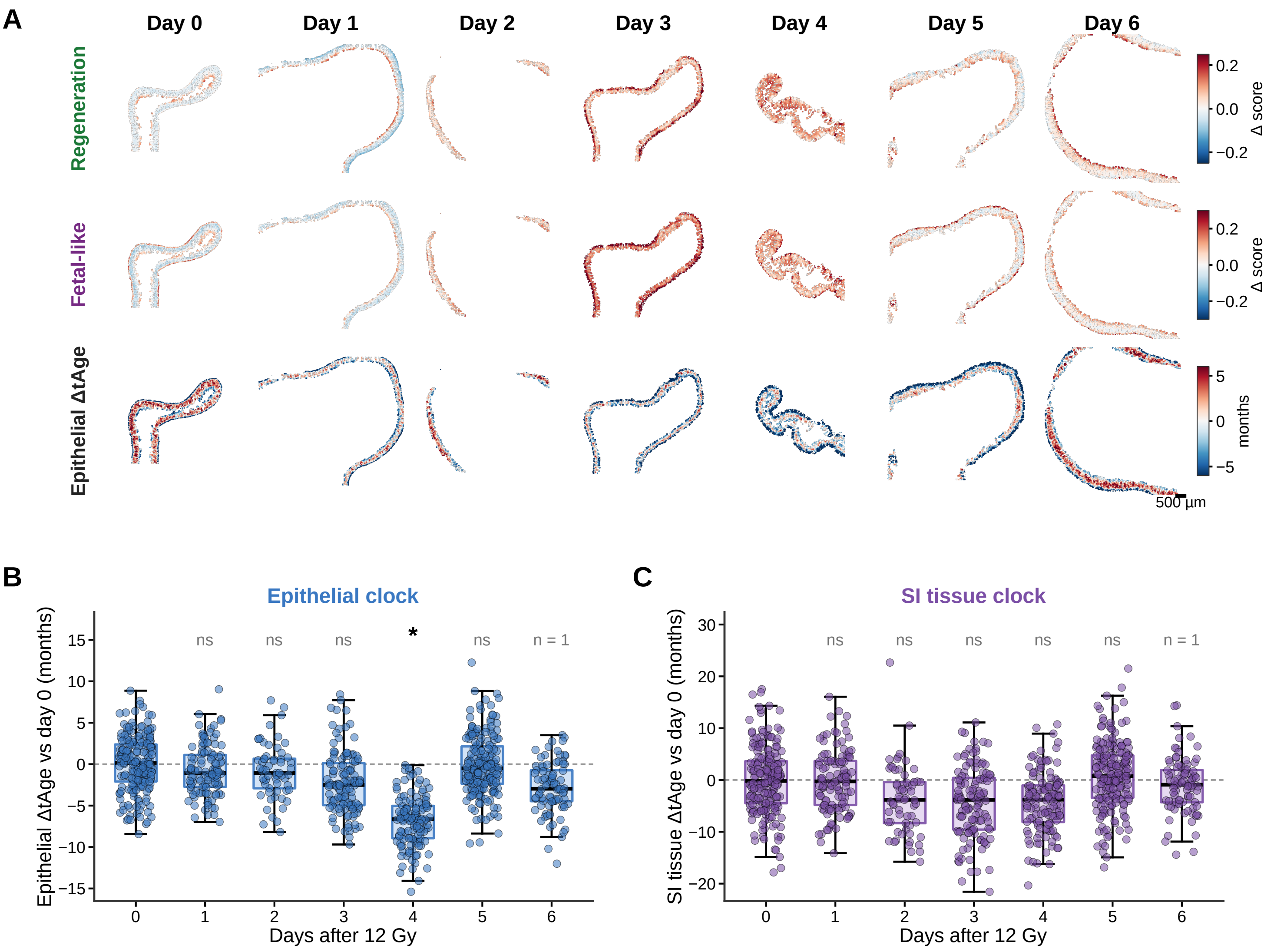

### Extended Data Fig. 13 | Spatial epithelial age across the irradiation time course in replicated Visium HD jejunum.

Jejunum before (day 0) and 1 to 6 days after 12 Gy whole-body irradiation, profiled by Visium HD on four capture areas that each contain a day-0 mouse (GSE303705; 23 mice in the deposited experiment, 20 assignable from the authors' bin-level labels and analyzed, 17 with metaspots of at least 200,000 counts). Twenty mice were retained at the metadata stage, and 17 contributed epithelial metaspots passing the count threshold. **(A)** Maps of one mouse per day (days 0, 1, 3 and 5, capture area 2; days 2, 4 and 6, capture area 4) in epithelial units of 3 × 3 eight-µm bins (24 µm): regeneration and fetal-like scores minus the mean of the day-0 units of the same capture area, and epithelial ΔtAge relative to those units (smoothed over seven nearest neighbors for display). Scale bar, 500 µm. **(B, C)** Epithelial-clock (B) and SI-tissue-clock (C) ΔtAge of epithelial metaspots (320 µm, at least 200,000 counts; dots), relative to all day-0 metaspots and adjusted for capture area; boxes, median and interquartile range; whiskers, 1.5 × interquartile range. Metaspots (mice): day 0, 195 (3); day 1, 112 (2); day 2, 61 (3); day 3, 123 (3); day 4, 128 (2); day 5, 193 (3); day 6, 85 (1). In B and C, each mouse is summarized by the mean of its metaspots, and each day is compared with day 0 by ordinary least squares with capture area as a covariate (y ~ day + capture area; Benjamini–Hochberg adjustment across days within each readout); the day-6 mouse is not tested. Stars as in Extended Data Fig. 1.

**Extended Data Fig. 14 | Cell-type-resolved functional enrichment after 5-FU injury in young and old intestine**

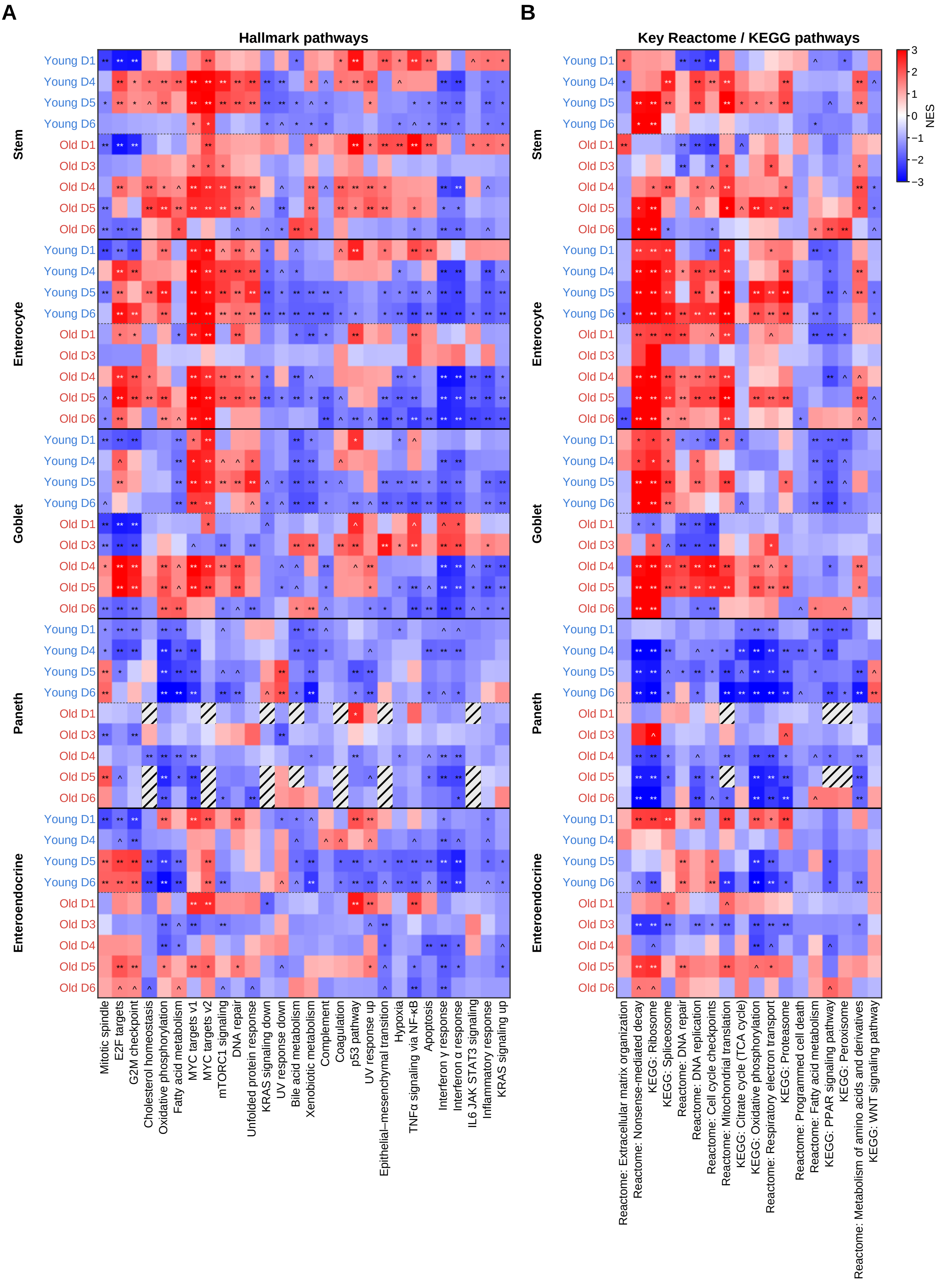

### Extended Data Fig. 14 | Cell-type-resolved functional enrichment after 5-FU injury in young and old intestine.

NES for 28 Hallmark sets (**A**; the sets with the largest mean absolute NES among those evaluated in more than 85% of contrasts, after excluding the eight sets HALLMARK_PANCREAS_BETA_CELLS, HALLMARK_SPERMATOGENESIS, HALLMARK_HEDGEHOG_SIGNALING, HALLMARK_ESTROGEN_RESPONSE_EARLY, HALLMARK_ESTROGEN_RESPONSE_LATE, HALLMARK_ANDROGEN_RESPONSE, HALLMARK_MYOGENESIS and HALLMARK_HEME_METABOLISM) and for a curated set of Reactome and KEGG pathways (**B**; chosen by the authors to cover protein synthesis and RNA processing, extracellular matrix organization, DNA replication and repair, mitochondrial metabolism, proteostasis, cell death, lipid and amino-acid metabolism and Wnt signaling, not by a significance rule) for stem, enterocyte, goblet, Paneth and enteroendocrine cells at each sampled day after 5-FU versus day 0 in mice of the same age (blue labels, young; red labels, old). Pseudobulk counts per mouse (two mice per group, including the low-yield old day-1 mouse, which was excluded only from the animal-level clock models) were ranked by a moderated t statistic; pre-ranked GSEA with 20,000 random gene sets as the null. ** FDR < 0.01, * FDR < 0.05, ^ FDR < 0.1 (BH within contrast and collection); hatched, pathway not evaluated. Young day 3 was not available. Paneth-cell yields were uneven at baseline (1,156 and 86 cells in the two young mice and 97 and 54 in the two old mice) and low after injury (2 to 55 cells per mouse on days 1 to 6), limiting the precision of the Paneth-cell estimates.

**Extended Data Fig. 15 | Pathway and module responses of young and old liver during regeneration**

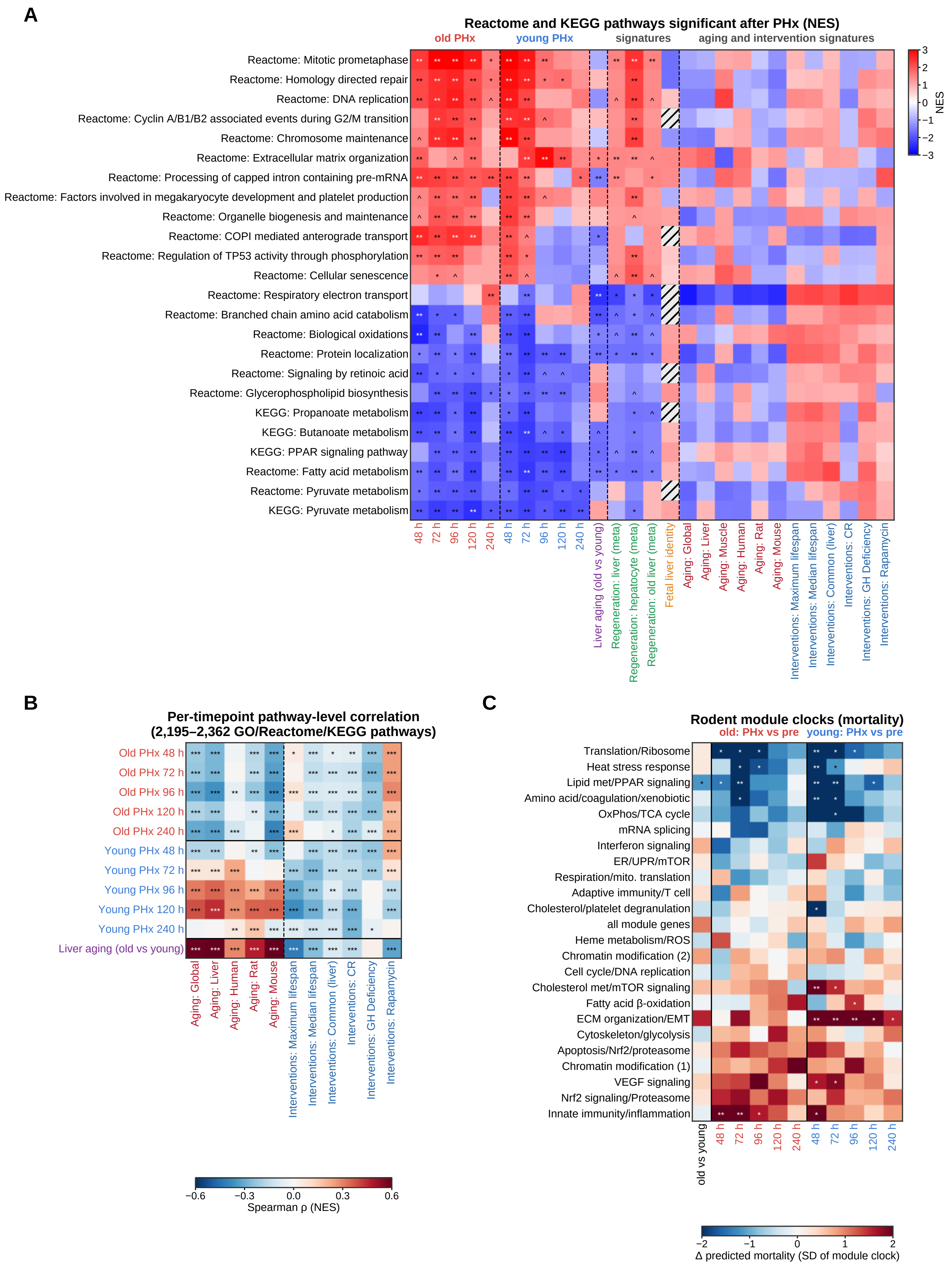

### Extended Data Fig. 15 | Pathway and module responses of young and old liver during regeneration.

Bulk liver RNA-seq after two-thirds partial hepatectomy (GSE185705; n = 3 livers per age group and timepoint); each PHx timepoint is compared with pre-PHx liver of the same age. **(A)** NES for 24 Reactome and KEGG pathways chosen by a fixed rule (evaluated in the reference matrix and FDR < 0.05 in at least one post-PHx contrast; the 12 with the smallest FDR in each direction, skipping a pathway that shares more than half of its genes with a higher-ranked one; Methods) for each post-PHx timepoint, pre-surgical liver aging, liver regeneration meta-signatures (liver, hepatocyte and old liver; aggregated lists in Supplementary Table 8; per-dataset contributors and contrasts were not retained), the fetal liver identity signature (fetal liver versus other fetal tissues) and published aging and intervention signatures (pre-ranked GSEA, 20,000 random gene sets as the null; KEGG sets mapped to mouse genes through one-to-one orthologs; ** FDR < 0.01, * < 0.05, ^ < 0.1; reference NES without FDR; hatched, not available). **(B)** Spearman correlation of pathway NES for each post-PHx timepoint and for pre-surgical aging with published aging signatures and intervention signatures (2,195–2,362 pathways per comparison; stars, BH q across pairs; the tests treat pathways as independent, so the q values are descriptive and do not account for genes shared between pathways). **(C)** Rodent mortality module clocks, labeled by their dominant pathways and shown as in Fig. 7D (change from pre-PHx liver in standard deviations of each module estimate; stars, pooled-variance contrast with Benjamini–Hochberg q across the 24 rows (23 modules and the all-module-gene estimate) and ten post-PHx contrasts; old-versus-young column, Welch P). Module-level significance in C depends on the pooled-variance assumption, with Welch-test sensitivity results provided in Supplementary Table 7.
